# Evolutionary dynamics of bipartite begomoviruses revealed by complete genome analysis

**DOI:** 10.1101/2020.06.25.171728

**Authors:** César A.D. Xavier, Márcio T. Godinho, Talita B. Mar, Camila G. Ferro, Osvaldo F.L. Sande, José C. Silva, Roberto Ramos-Sobrinho, Renato N. Nascimento, Iraildes Assunção, Gaus S.A. Lima, Alison T.M. Lima, F.Murilo Zerbini

**Author notes:** Department of Entomology and Plant Pathology, North Carolina State University, Raleigh, NC 27695, USA. Embrapa Trigo, Rodovia BR 285, Km 294, s/n, Passo Fundo, RS 99050-970, Brazil. Dep. de Fitopatologia e Nematologia, Universidade de São Paulo, Av. Pádua Dias 11, Piracicaba, SP 13418900, Brazil. School of Plant Sciences, University of Arizona, 1140 E. South Campus Drive, Tucson, AZ 85721, USA.

## Abstract

Several key evolutionary events marked the evolution of geminiviruses, culminating with the emergence of bipartite genomes represented by viruses classified in the genus *Begomovirus*. This genus represents the most abundant group of multipartite viruses, contributing significantly to the observed abundance of multipartite species in the virosphere. Although aspects related to virus-host interactions and evolutionary dynamics have been extensively studied, the bipartite nature of these viruses has been little explored in evolutionary studies. We performed a parallel evolutionary analysis of the DNA-A and DNA-B components of New World begomoviruses. A total of 239 full-length DNA-B sequences obtained in this study, combined with 292 DNA-A and 76 DNA-B sequences retrieved from GenBank, were analyzed. The results indicate that the DNA-A and DNA-B respond differentially to evolutionary processes, with the DNA-B being more permissive to variation and more prone to recombination than the DNA-A. Although a clear geographic segregation was observed for both components, differences in the genetic structure between DNA-A and DNA-B were also observed, with cognate components belonging to distinct genetic clusters. DNA-B coding regions evolve under the same selection pressures than DNA-A coding regions. Together, our results indicate an interplay between reassortment and recombination acting at different levels across distinct subpopulations and components.

## Introduction

Although structurally simple, viruses possess complex evolutionary histories (1–5). Throughout their evolutionary process, viruses incorporated several unique features, exhibiting a wide diversity of genome organizations, mechanisms of replication and gene expression strategies (6–8). This diversity allows viral populations to be dynamic, with a high adaptive capacity, infecting hosts in the three domains of life (1, 9, 10). An intriguing aspect is the emergence of viruses with segmented genomes (11–13). Segmented genomes can be found in families of DNA and RNA viruses infecting animals, plants and fungi (14). The biological and evolutionary significance of genome segmentation remains unclear (15–18).

A special case of genome segmentation are viruses with multipartite genomes, which have their genome segments packed into separate particles. The existence of viruses with multipartite genomes presents further challenges to the biological and evolutionary purpose of genome segmentation (19). While each segment reaches complete independence (since there is no physical connection between them), they need to act together to complete the infectious cycle. Therefore, multipartite genomes can suffer a high adaptative cost, especially during intercellular movement and transmission between hosts, because in order for all segments to be present in the next cell or transmitted to a new host, all segments must be in a high multiplicity of infection (MOI) (19–23). Alternatively, it has been suggested that the lower number of errors introduced per replication round in small genome segments may counterbalance the disadvantageous effects of multipartite genomes (12). In addition, Ojosnegros *et al*. (24) proposed that increased capsid stability due to the encapsidation of smaller genomic components is a factor that can at least in part contribute to the maintenance of multipartite genomes. Evolutionarily, it has been proposed that reassortment (the exchange of genomic components between strains or related species) in segmented viruses may represent a form of “sexual reproduction”, allowing the maintenance of genomic integrity through the elimination of deleterious mutations (25) and increasing population-level genotypic diversity (26).

A number of studies aiming to understand the evolutionary dynamics of different components at population level have been conducted (27–32). Given the trade-offs between functional complementation and independence among the distinct components in multipartite viruses, it would be expected that the different components would be in an intimate process of co-evolution, with similar evolutionary histories. In support of this hypothesis, some of the above-mentioned studies indicated that reassortants between isolates of multipartite plant viruses did not become established in the population (27). However, other studies suggested that the different components of multipartite genomes may undergo distinct selection pressures, and thus experience different evolutionary histories (28, 30, 31). These conflicting results could be related to varying degrees of interdependency among the proteins encoded by genomic segments of distinct viruses. For example, genomic segments that encode components of the viral replicase would be strongly dependent of each other, while being relatively independent of another segment that encodes a movement protein.

The genus *Begomovirus* (family *Geminiviridae*) is comprised of viral species with one or two genomic components of circular, single-strand DNA (ssDNA) of about 2,700 nucleotides, encapsidated in geminate icosahedral particles and transmitted to dicotyledonous plants by whiteflies of the *Bemisia tabaci* cryptic species complex (33). Begomoviruses constitute are responsible for severe diseases in crops and ornamentals plants of economic importance worldwide (34). Beyond their economic importance, two main characteristics make begomoviruses an attractive model to study the evolutionary dynamics of viral populations. First, begomoviruses evolve at rates which are comparable to those of ssRNA viruses, with estimates of substitution rates in the order of 10^−3^-10^−4^ substitutions/site/year (35–37). Second, begomoviruses include viruses with non-segmented as well as bipartite genomes. The bipartite nature of these viruses has been little explored in evolutionary studies, and little is known about the evolutionary dynamics of the different components.

The two genomic components of bipartite begomoviruses are referred to as DNA-A and DNA-B. The two components share no significant sequence identity, except for an intergenic region (IR) of approximately 200 nucleotides. The IR includes the replication origin (*ori*), a conserved stem-loop structure with the invariable nonanucleotide TAATATT//AC, and conserved repeat sequences (iterons) that are specifically recognized by the viral replication-associated protein, Rep (38–40). The IR is important for maintaining the integrity of the bipartite genome, allowing both components to be replicated by Rep, since Rep proteins show high specificity for their cognate *ori* (38).

Although the mechanism by which segmented and multipartite genomes have emerged is unclear, evidence suggests they may originate by fragmentation of the genome of non-segmented progenitors, with defective segments becoming infectious by complementation (20). Specifically for begomoviruses, Briddon *et al*. (31) suggested that the DNA-B could have originated from a satellite molecule captured by the monopartite progenitor of all begomoviruses. Possibly, this association provided greater adaptability to the monopartite progenitor, and consequently was maintained over the evolutionary process.

Understanding the processes driving the patterns of population genetic differentiation and genetic variability of viruses has been a long-standing focus in the area of evolutionary virology. A number of studies have been conducted to determine and quantify the contribution of the mechanisms dictating the evolutionary dynamics of begomovirus populations (35–37, 41–47). However, these studies were based on analysis of non-segmented viruses or of the DNA-A component of bipartite viruses. Considering the important role played by the DNA-B in the infection process, evolutionary studies that consider this component are needed. The analysis of the complete genome can provide a more accurate picture of the evolution of begomovirus and a better understanding of the evolution of multipartite genomes in general.

Here, using a population genetics approach, we performed a parallel evolutionary analysis of the DNA-A and DNA-B component of NW begomoviruses from cultivated and non-cultivated hosts. The two components respond differentially to evolutionary processes, with the DNA-B being more variable and more prone to recombination than the DNA-A. A clear geographic segregation was observed for both components, but differences between the genetic structure of cognate DNA-A and DNA-B components could be observed. Interestingly, DNA-B coding regions evolve at least under the same selection pressures than DNA-A coding regions. Our results indicate an interplay between reassortment and recombination acting at different levels across distinct subpopulations and genomic components.

## Results

A total of 239 full-length DNA-B sequences were obtained in this study, all from samples of cultivated and non-cultivated plants from which a DNA-A had been previously cloned and analyzed (41, 42, 48). These sequences were combined with 53 DNA-A and 76 DNA-B sequences retrieved from GenBank. Detailed information on the samples and the corresponding DNA-A and DNA-B sequences are presented in Figure 1 and Suppl. Tables S1 and S2. The global data set consisted of a total of 292 DNA-A and 315 DNA-B sequences: 117 DNA-A and 123 DNA-B sequences of *Bean golden mosaic virus* (BGMV), 30 DNA-A and 41 DNA-B sequences of *Blainvillea yellow spot virus* (BlYSV), 50 DNA-A and 53 DNA-B sequences of *Euphorbia yellow mosaic virus* (EuYMV), 21 DNA-A and 24 DNA-B sequences of *Macroptilium yellow spot virus* (MaYSV) and 74 DNA-A and 74 DNA-B sequences of *Tomato severe rugose virus* (ToSRV) (Figure 1). For each species data set, the number of DNA-A and DNA-B sequences was similar (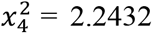, *P* = 0.6911).

**Figure 1.**
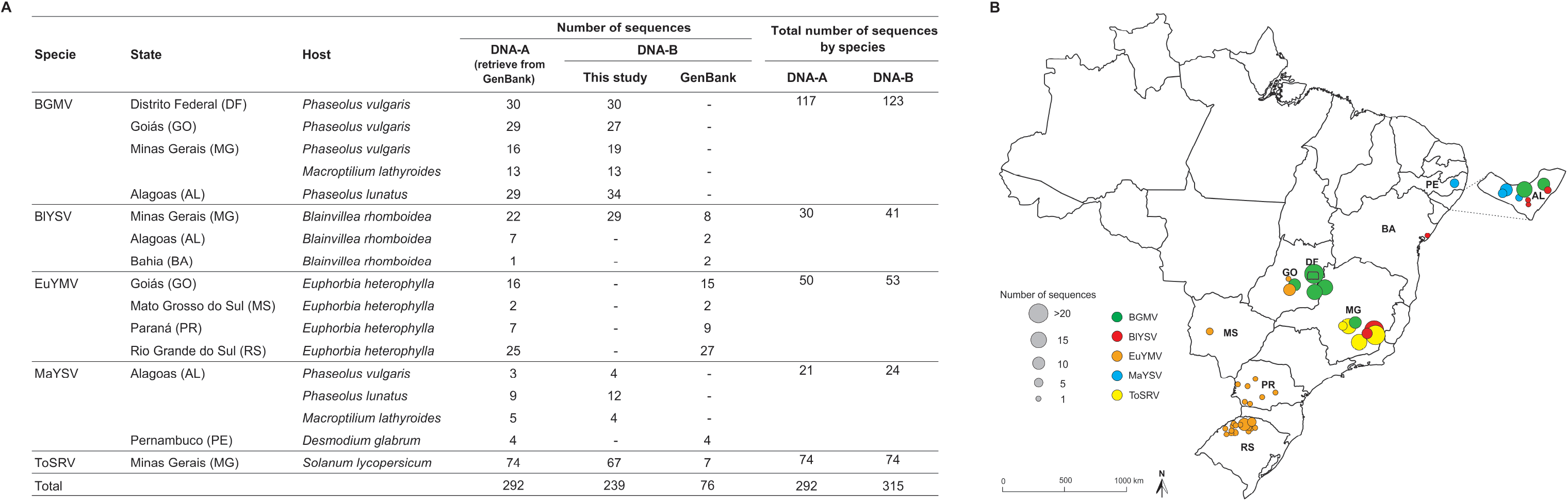
(**A**) Begomovirus sequences analyzed in this study. The map of Brazil (**B**) shows the origin of the sequences sampled in different regions of country. Circles indicate the location where samples were collected. Circle size represents the number of sequences (DNA-A and DNA-B) obtained from samples collected at each location. AL, Alagoas; BA, Bahia; DF, Distrito Federal; GO, Góias; MG, Minas Gerais; MS, Mato Grosso do Sul; PE, Pernambuco; PR, Paraná; RS, Rio Grande do Sul.

### Phylogenetic analysis

Bayesian phylogenetic trees based on full-length DNA-A and DNA-B nucleotide sequences were constructed for each species data set (Figure 2). Although the PACo analysis provided significant evidence for global congruence between DNA-A and DNA-B trees (*P*≤0.001; Table 1), the residual square sum (*m^2^_XY_*) values were variable across the different data sets (0.1079 – 0.6039; Table 1), indicating that congruence levels are variable among them. Visual analysis of phylogenetic trees indicated the formation of large clades according to geographical region, for both components (Figure 2), providing initial evidence of population subdivision based on geography.

**Figure 2.**
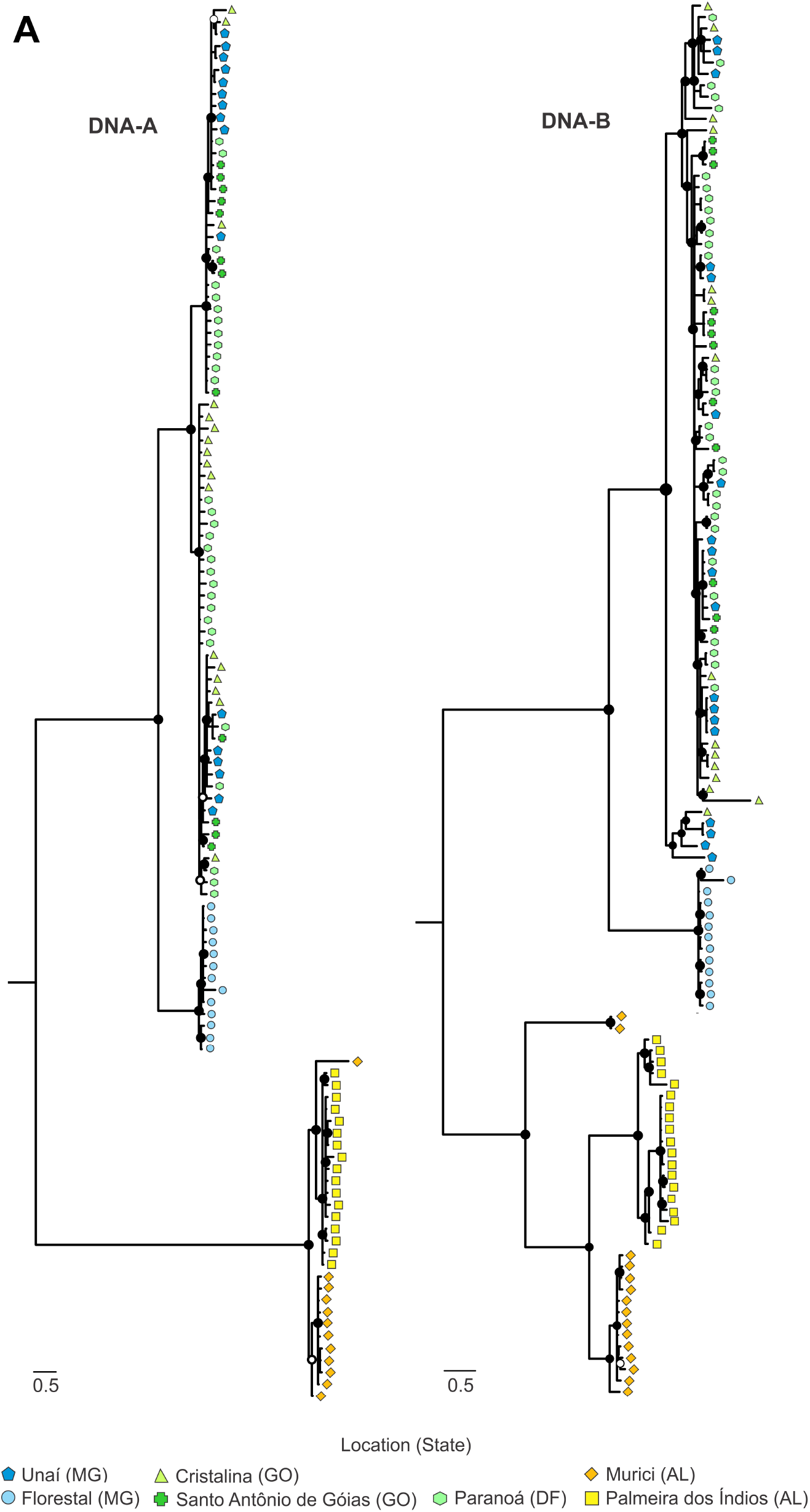
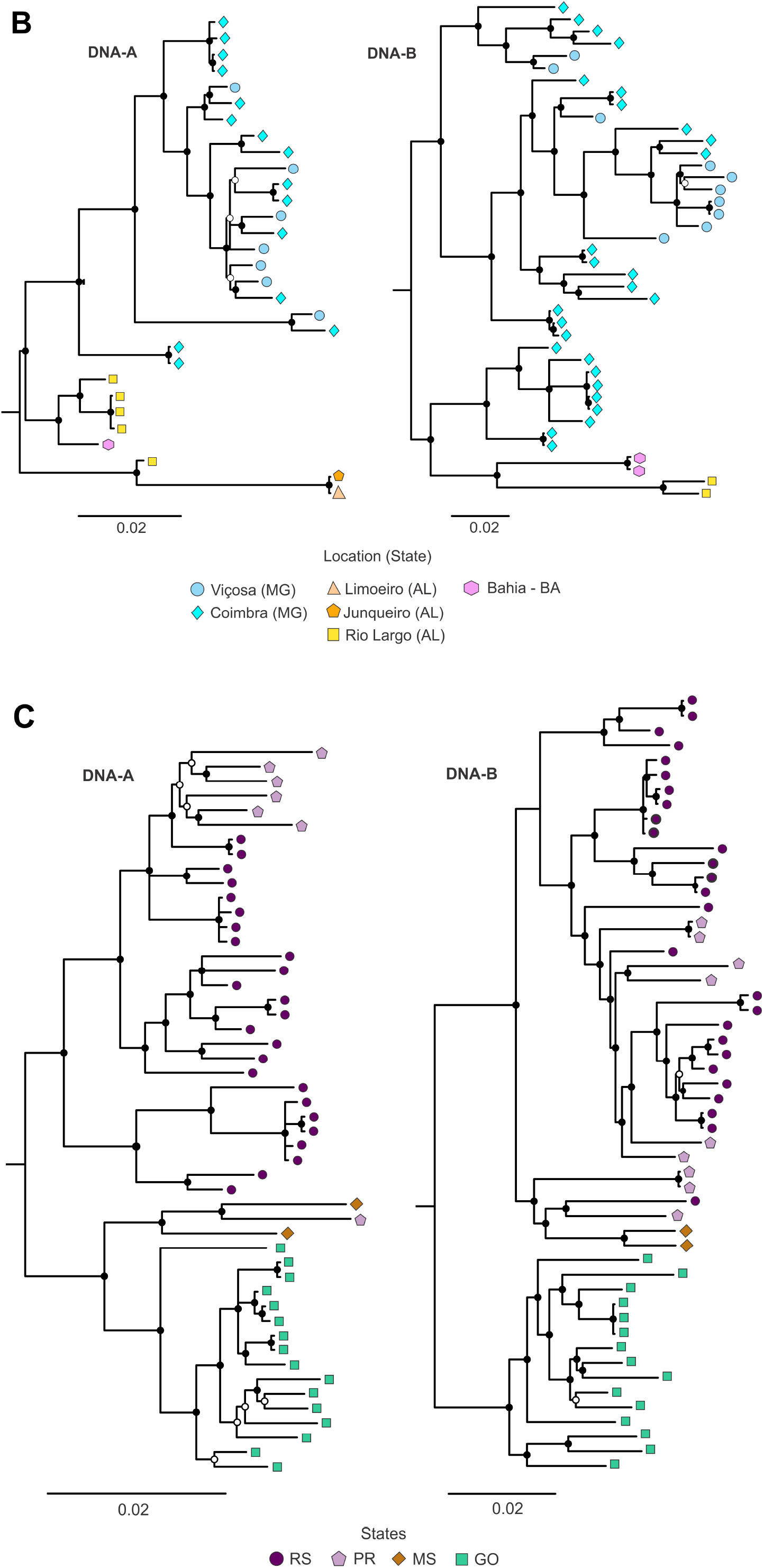
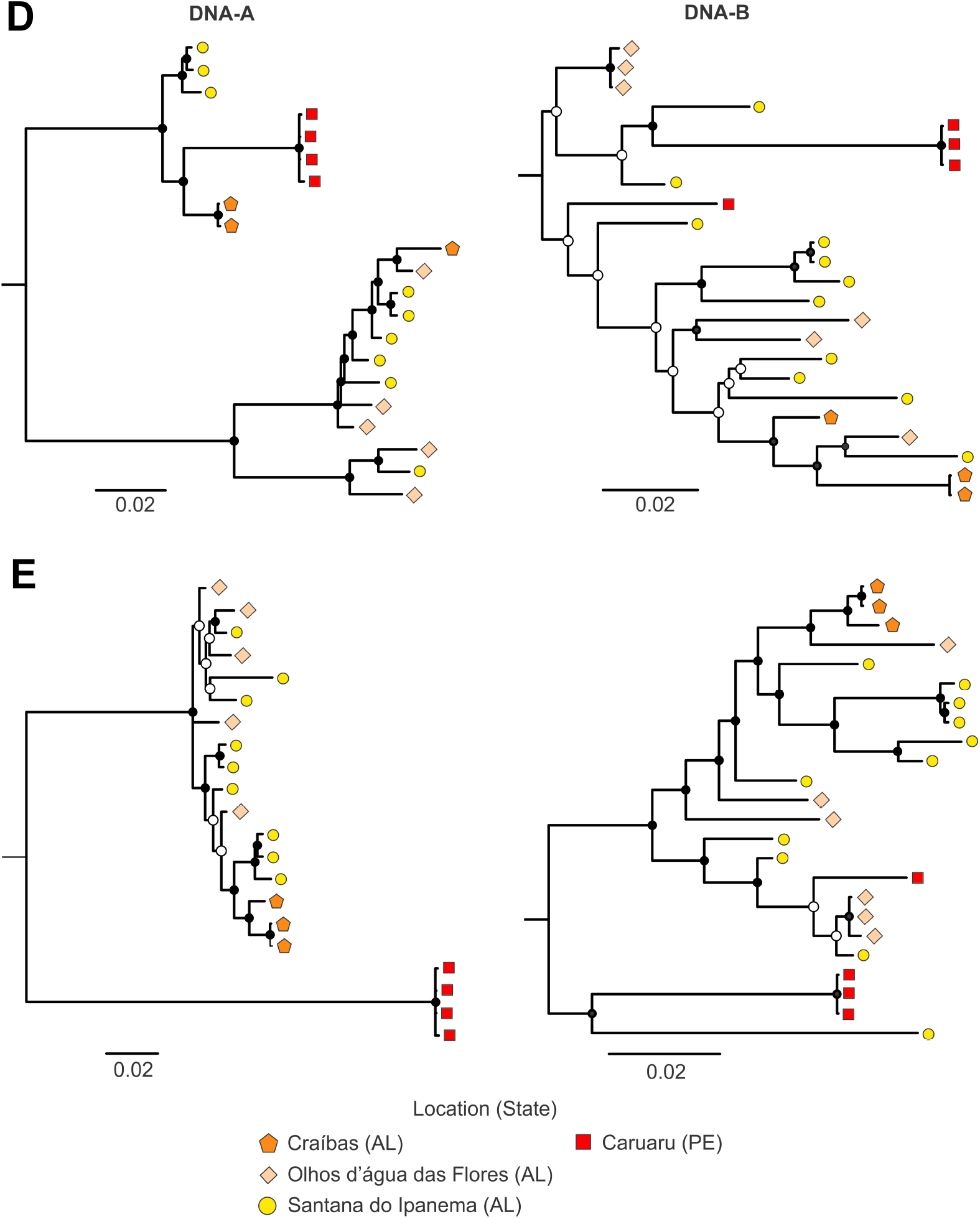
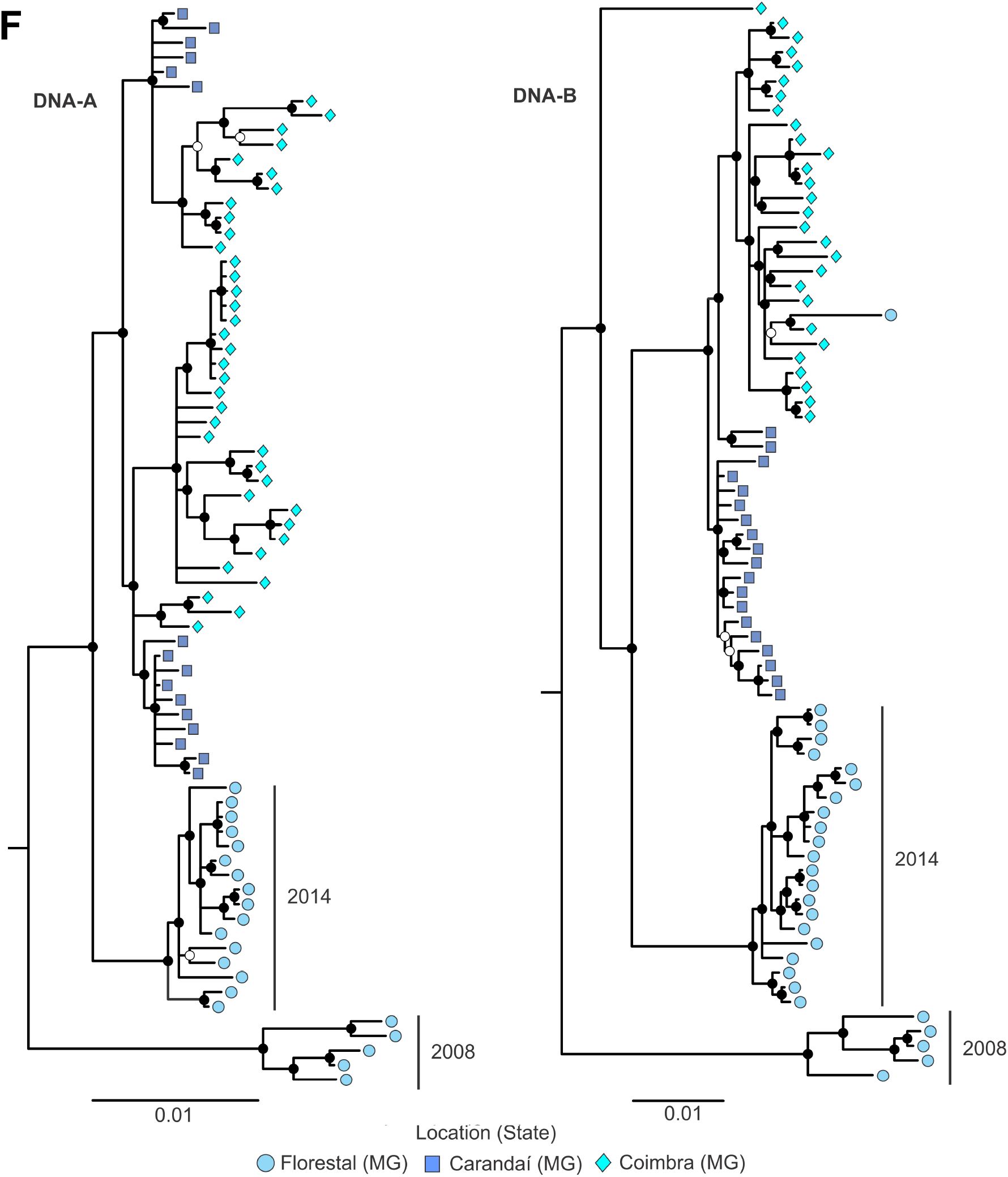
Midpoint-rooted Bayesian phylogenetic trees based on full-length nucleotide sequences of the DNA-A and DNA-B segments of (**A**) *Bean golden mosaic virus* (BGMV), (**B**) *Blainvillea yellow spot virus* (BlYSV), (**C**) *Euphorbia yellow mosaic virus* (EuYMV), (**D**, **E**) *Macroptilium yellow spot virus* (MaYSV), and (**F**) *Tomato severe rugose virus* (ToSRV). **E**, Phylogenetic tree inferred using a data set in which recombinant blocks were excluded. Nodes with posterior probability values between 0.50 and 0.80 are indicated by empty circles and nodes with values equal to or greater than 0.81 are indicated by filled circles. The scale bar represents the number of nucleotide substitutions per site. Isolate color indicates sampling location.

**Table 1.** Results of co-phylogeny analysis between begomovirus DNA-A and DNA-B segments using Procrustean Approach to Cophylogeny (PACo) for *Bean golden mosaic virus* (BGMV), *Blainvillea yellow spot virus* (BlYSV), *Euphorbia yellow mosaic virus* (EuYMV), *Macroptilium yellow spot virus* (MaYSV) and *Tomato severe rugose virus* (ToSRV) data sets.

| Virus | Global goodness-of-fit |  |
| --- | --- | --- |
| | $m^2_{xy}$* | P-value |
| BGMV<sup>#</sup> | 0.4992 | 0.001 |
| BGMV<sup>&</sup> | 0.5411 | 0.001 |
| BIYSV<sup>#</sup> | 0.1682 | 0.001 |
| BIYSV<sup>&</sup> | 0.1623 | 0.001 |
| EuYMV<sup>#</sup> | 0.3059 | 0.001 |
| EuYMV<sup>&</sup> | 0.2970 | 0.001 |
| MaYSV<sup>#</sup> | 0.6039 | 0.001 |
| MaYSV<sup>&</sup> | 0.6613 | 0.001 |
| ToSRV | 0.1079 | 0.001 |

\* Residual square sum for global test, which is inversely proportional to the level of global congruence between phylogenies.
<sup>#</sup> Alignment includes recombinant blocks.
<sup>&</sup> Alignment does not include recombinant blocks.

DNA-A and DNA-B phylogenetic trees for BGMV were perfectly symmetric, yielding “mirror image” trees (Figure 2A). In both trees it is possible to visualize two major clades supported by high posterior probability values and separated by long branches (Figure 2A). The first clade (DF/GO/MG) is comprised of isolates collected in Distrito Federal (DF), Góias (GO) and Minas Gerais (MG) states, and the second clade (AL) is comprised of isolates sampled in Alagoas (AL) state. In addition, well-defined subclusters can be observed for both components, suggesting the existence of a substructure within a smaller geographical scale. The DF/GO/MG cluster can be subdivided into isolates from Florestal (MG) and from Paranoá (DF), Cristalina (GO), Santo Antônio de Goiás (GO) and Unaí (MG). The AL cluster also presented two subclusters, one comprised of isolates sampled in Murici and another by isolates from Palmeira dos Índios (Figure 2A).

Although the BlYSV data set contains isolates sampled in distant geographical regions (approximately 1800 km apart), phylogenetic trees do not display a clear and consistent pattern of geography-based clustering as noted for other species data sets (Figure 2B). There was a tendency in isolates collected in the northeastern states (AL and Bahia, BA) to form separate clades from isolates collected in the southeast (MG), however short internal branches can be observed, suggesting a low level of differentiation. In addition, recombination events were detected in DNA-A components among isolates of the northeast with putative major parents in the southeast (Suppl. Table S3). For the DNA-B, isolates sampled in the southeast showed recombination events with putative minor parents in the northeast (Suppl. Table S3). These results suggest that, despite the greater distance between sampled sites, some connection between the two regions exists, resulting in an incipient pattern of differentiation. Sampling of intermediate regions between AL and MG may provide a better picture of the genetic make-up of the BlYSV population.

In agreement with Mar *et al*. (44), a near perfect segregation based on geography was observed in the EuYMV DNA-A and DNA-B trees (Figure 2C). For the DNA-A, two major clades were well supported, with isolates sampled in Rio Grande do Sul (RS) and Paraná states (PR) clustered separately from those sampled in Góias (GO) and Mato Grosso do Sul (MS). The same clusters could be observed in the DNA-B tree, except that isolates sampled in MS clustered with those from RS and PR, suggesting a reassortment event.

Due to the complex recombination profile of the MaYSV data set (see below), DNA-A and DNA-B phylogenetic trees showed a high degree of incongruence and displayed no obvious pattern of segregation (Figure 2D). For the DNA-A tree two well-supported major clades can be observed, while the DNA-B tree shows several poorly supported clades (Figure 2D). To minimize the effect of recombination, phylogenetic trees were reconstructed based on recombination-free DNA-A and DNA-B data sets (Figure 2E). Although MaYSV isolates were sampled in a smaller geographical scale (maximum distance among fields of approximately 200 kilometers) and a smaller number of sequences were obtained compared to the other data sets, two well-supported clades based in geographical location can be observed in the DNA-A tree, with isolates sampled in Pernambuco (PE) state clustered separately from those collected in neighboring AL (Figure 2E). Although two well-supported clades were also observed in the DNA-B tree, these were not perfectly congruent with the DNA-A tree (Figure 2E). Only two DNA-B haplotypes were sampled in PE, with the first (isolates BR-PE-Cau-22-3, BR-PE-Cau-22-4 and BR-PE-Cau-22-5), clustering together with BR-Sti-34-11 sampled in AL, and the second (BR-PE-Cau-23-1) clustering with the remaining sequences also sampled in AL. Therefore, a larger number of haplotypes sampled in PE will be necessary to obtain a better resolution of the pattern of DNA-B segregation.

Phylogenetic trees for ToSRV DNA-A and DNA-B were not completely congruent (Figure 2F). A clear geographical clustering was observed for the DNA-B tree, with three clades comprised almost exclusively of isolates from Coimbra, Carandaí and Florestal/2014. Moreover, a fourth cluster contained isolates from Florestal/2008, indicating a remarkable temporal structure considering the short period of time between samplings (only six years). The DNA-A tree displayed the same topology as the DNA-B tree for the isolates from Florestal, but isolates from Coimbra and Carandaí were not segregated (Figure 2F). Interestingly, the DNA-B of isolate BR-Flo01-14 from Florestal clustered together with isolates sampled in Coimbra, due to a recombination event involving parental sequences from these two locations (Suppl. Table S3). This indicates that the exchange of components between isolates from Coimbra and Carandaí has occurred.

### Genetic structure of populations

To compare the genetic structure based on DNA-A and DNA-B components, a nonparametric multivariate statistical analysis was performed using DAPC (Figure 3; Suppl. Table S4). Since DAPC requires predefined groups, the analysis was run sequentially with *k* values varying from 2 to 10, with the best number of genetic clusters chosen based on the minimum number of *k* after which the Bayesian Information Criterion (BIC) decreased by an insignificant amount. When a clear limit after which BIC values decreased by an insignificant amount could not be observed, the percentage of variation among genetic clusters inferred by *k*-means was plotted against BIC according to analysis of molecular variance (AMOVA) (Suppl. Figure S1). For most data sets it was possible to define clear genetic clusters according to the criteria mentioned above. However, for BlYSV DNA-A and DNA-B and MaYSV DNA-B component data sets, in agreement with phylogenetic analysis, it was not possible to define consistent genetic clusters (Suppl. Figure S1). Therefore, for these three component data sets all individuals were considered to constitute a single population in later analyses.

**Figure 3.**
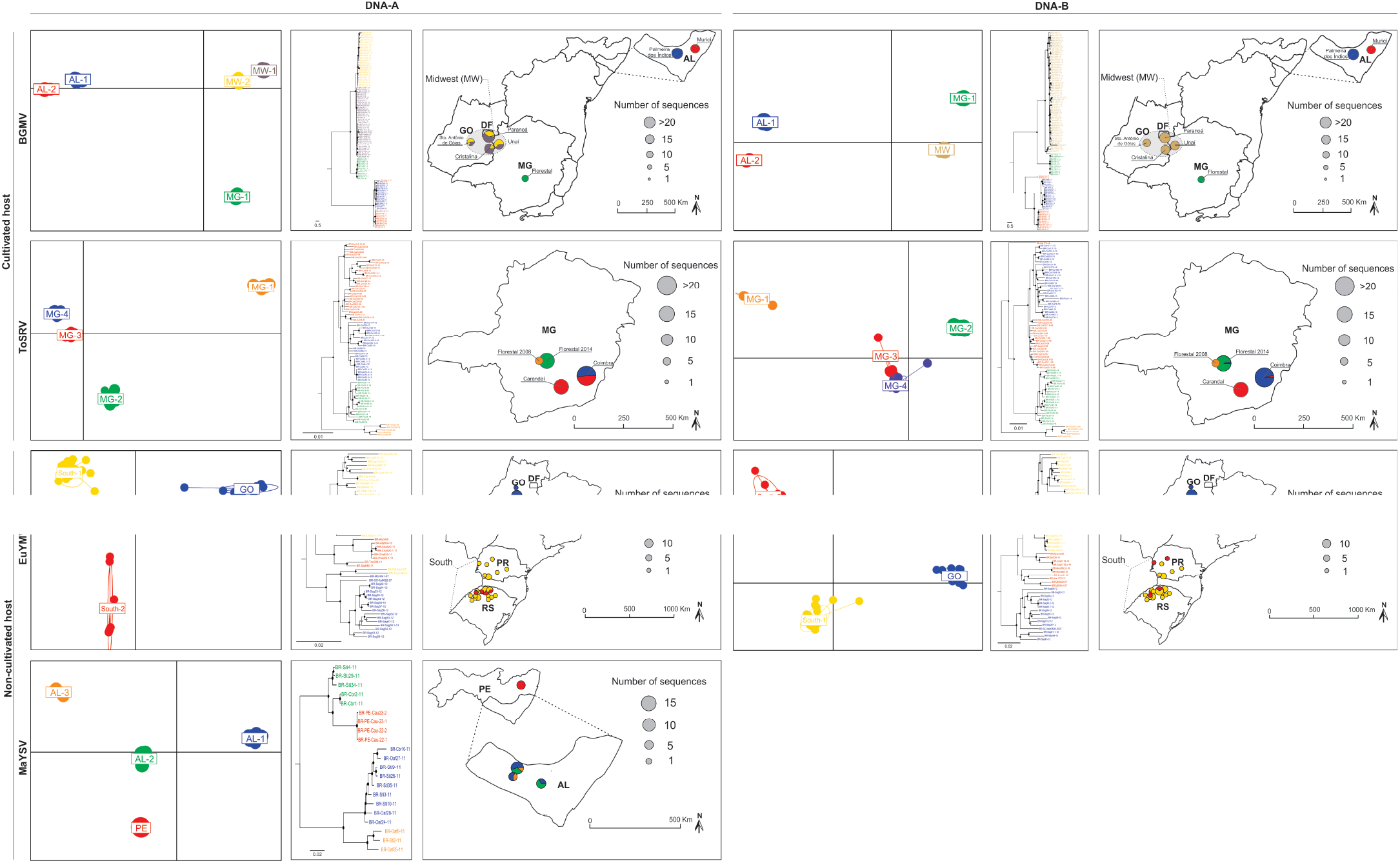
Analysis of population structure using Discriminant Analysis of Principal Components (DAPC) for *Bean golden mosaic virus* (BGMV), *Euphorbia yellow mosaic virus* (EuYMV), *Macroptilium yellow spot virus* (MaYSV) and *Tomato severe rugose virus* (ToSRV) data sets. DAPC scatterplots (left panels) show the first two principal components with ellipses representing genetic clusters and dots representing each isolate. Isolates in the Bayesian phylogenetic trees (center panels) are colored based on genetic clusters identified in the DAPC analysis. Pie-charts in the maps (right panels) represent the genetic composition of subpopulations according to DAPC analysis. Pie-chart size represents the number of sequences obtained from samples collected at each location. Although there are genetics clusters with the same names in the DNA-A and DNA-B data sets of some viral species, the composition of isolates in these clusters is different.

In general, DAPC analysis across all data sets consistently discriminated all pre-defined clusters based on *k*-means (data not shown). Although a clear geographical segregation can be observed for the two components, especially for BGMV, EuYMV and ToSRV (consistent with phylogenetic analysis), slight differences between the genetic structure based on DNA-A and DNA-B could be observed, with cognate components belonging to distinct genetic clusters (Figure 3).

For BGMV, DAPC analysis discriminated five and four genetics clusters for DNA-A and DNA-B, respectively, primarily based on geography (Figure 3). The clusters named AL-1, AL-2 and MG-1 were observed for both components, displaying a perfect pattern based on sampling location (Figure 3). For isolates sampled in the midwestern region, two genetic clusters (MW-1 and MW-2) and a single cluster (MW) were detected for DNA-A and DNA-B, respectively (Figure 3). Interestingly, no clear pattern was observed to explain the formation of MW-1 and MW-2 clusters, with all four subpopulations from the midwest including DNA-A sequences from both genetic clusters (Figure 3). Although the difference in genetic structure between DNA-A and DNA-B could be due to the different number of clusters assigned in the *k*-means analysis, such difference was observed even when the same *k* number was tested for both components. To try to understand what could be influencing this process, we checked which sites in the DNA-A were contributing the most to the observed structure. We found 22 contributing sites located between the C-terminal region of REP and the C-terminal region of CP, mostly displaying synonymous substitutions. This suggests the action of genetic drift, rather than positive selection, leading to a change in the frequency of the two genotypes in the different subpopulations.

Similar to the patterns observed for BGMV, a clear geographical segregation was also observed for both EuYMV components, with only minor differences in the assignment of some DNA-A and DNA-B components to the three genetic clusters (South-1, South-2 and GO) (Figure 3).

Although it was not possible to define clear genetic clusters for MaYSV DNA-B components based on DAPC analysis, the DNA-A components were split into four groups (Figure 3). Isolates from PE clustered separately from isolates sampled in AL (Figure 3). Moreover, isolates from AL grouped into three separate genetic clusters (AL-1, AL-2, AL-3) with no association with sampling location or host (Figure 3). When the data were analyzed without recombinant blocks, only two clusters were formed, with isolates sampled in AL forming a single cluster separately from those sampled in PE (Suppl. Figure S2).

For ToSRV it was possible to discriminate four clusters for both components. However, there was no perfect correspondence between DNA-A and DNA-B clusters in the same locations (Figure 3). The MG-1 and MG-2 clusters showed the same composition for both components, comprised of isolates sampled in Florestal in 2008 and 2014, respectively. The DNA-A components sampled in Coimbra were split into two genetic clusters (MG-3 and MG-4), while DNA-A components from Carandaí were assigned only to the MG-3 cluster (Figure 3). For the DNA-B, in agreement with the phylogenetic analysis, isolates sampled in Carandaí and Coimbra were assigned to MG-3 and MG-4 clusters, respectively, except for BR-Coi179-14 from Coimbra which was assigned to the MG-3 cluster (Figure 3). These results suggest that asymmetric flow of DNA-A components can occur from Carandaí to Coimbra, but the opposite is not necessarily true, since the Carandaí subpopulation was entirely comprised of individuals belonging to same genetic cluster (Figure 3). In addition, there is not a large mixture of DNA-B components, suggesting that the exchange may be restricted to the DNA-A.

To assess the consistence of the genetic clusters inferred by DAPC analysis, the differentiation degree among the clusters was estimated by means of the coefficient of nucleotide differentiation (Nst) (Table 2). In general, high values of pairwise Nst were observed among different clusters, indicating great differentiation, and fully supporting the subdivision proposed by DAPC analysis.

**Table 2.** Results of subdivision test performed for *Bean golden mosaic virus* (BGMV), *Euphorbia yellow mosaic virus* (EuYMV), *Macroptilium yellow spot virus* (MaYSV) and *Tomato severe rugose virus* (ToSRV) data sets.

| Virus | Genomic component |  |  |  |  |  |
| --- | --- | --- | --- | --- | --- | --- |
|  | DNA-A* |  |  | DNA-B* |  |  |
|  | Cluster 1 | Cluster 2 | Nst<sup>#</sup> | Cluster 1 | Cluster 2 | Nst |
| BGMV | MW-2 | MW-5 | 0.704 | MW | MG-1 | 0.892 |
|  | MW-2 | MG-1 | 0.922 | MW | AL-2 | 0.868 |
|  | MW-2 | AL-2 | 0.966 | MW | AL-1 | 0.925 |
|  | MW-2 | AL1-1 | 0.975 | - | - | - |
|  | MW-5 | MG-1 | 0.908 | - | - | - |
|  | MW-5 | AL-2 | 0.964 | - | - | - |
|  | MW-5 | AL-1 | 0.973 | - | - | - |
|  | MG-1 | AL-2 | 0.971 | MG-1 | AL-2 | 0.911 |
|  | MG-1 | AL-1 | 0.981 | MG-1 | AL-1 | 0.962 |
|  | AL-2 | AL-1 | 0.695 | AL-2 | AL-1 | 0.680 |
| EuYMV | GO | SO-1 | 0.487 | GO | SO-2 | 0.363 |
|  | GO | SO-2 | 0.640 | GO | SO-1 | 0.448 |
|  | SO-1 | SO-2 | 0.442 | SO-2 | SO-1 | 0.215 |
| MaYSV | AL-2 | AL-1 | 0.834 | - | - | - |
|  | AL-2 | PE | 0.827 | - | - | - |
|  | AL-2 | AL-3 | 0.801 | - | - | - |
|  | AL-1 | PE | 0.929 | - | - | - |
|  | AL-1 | AL-3 | 0.726 | - | - | - |
|  | PE | AL-3 | 0.895 | - | - | - |
| ToSRV | MG-3 | MG-4 | 0.420 | MG-3 | MG-4 | 0.316 |
|  | MG-3 | MG-1 | 0.718 | MG-3 | MG-1 | 0.830 |
|  | MG-3 | MG-2 | 0.623 | MG-3 | MG-2 | 0.778 |
|  | MG-4 | MG-1 | 0.752 | MG-4 | MG-1 | 0.785 |
|  | MG-4 | MG-2 | 0.703 | MG-4 | MG-2 | 0.709 |
|  | MG-1 | MG-2 | 0.776 | MG-1 | MG-2 | 0.811 |

\* Although there are subpopulations with the same names in the DNA-A and DNA-B data sets of some viral species, the composition of isolates in these clusters is different. AL, Alagoas; GO, Goiás; MG, Minas Gerais; MW, Midwest; PE, Pernambuco; SO, South
<sup>#</sup> Values from 0 to 0.05 indicate little genetic differentiation; from 0.05 to 0.15, moderate differentiation; from 0.15 to 0.25, great differentiation; > 0.25, high differentiation.

### Genetic variability

Although there is evidence that DNA-B components are more variable than DNA-A components (31) a quantitative and comparative analysis using large intraspecific data sets has not yet been performed. To address this issue, we calculated the average pairwise number of nucleotide differences (nucleotide diversity index, π) for the DNA-A and DNA-B components from the five species data sets. The π values for the DNA-B were statistically higher than those for the DNA-A for almost all species data sets (Figure 4A; Suppl. Tables S5 and S6). The exception was MaYSV, for which the value for the DNA-A (π=0.07170) was statistically higher than for the DNA-B (π=0.05567) (Figure 4A; Suppl. Table S5 and S6). These results confirm the highest variabilty of the DNA-B in relation to the DNA-A at the populational level. However, this is not an absolute rule, as exemplified by MaYSV, which has a more variable DNA-A than DNA-B.

**Figure 4.**
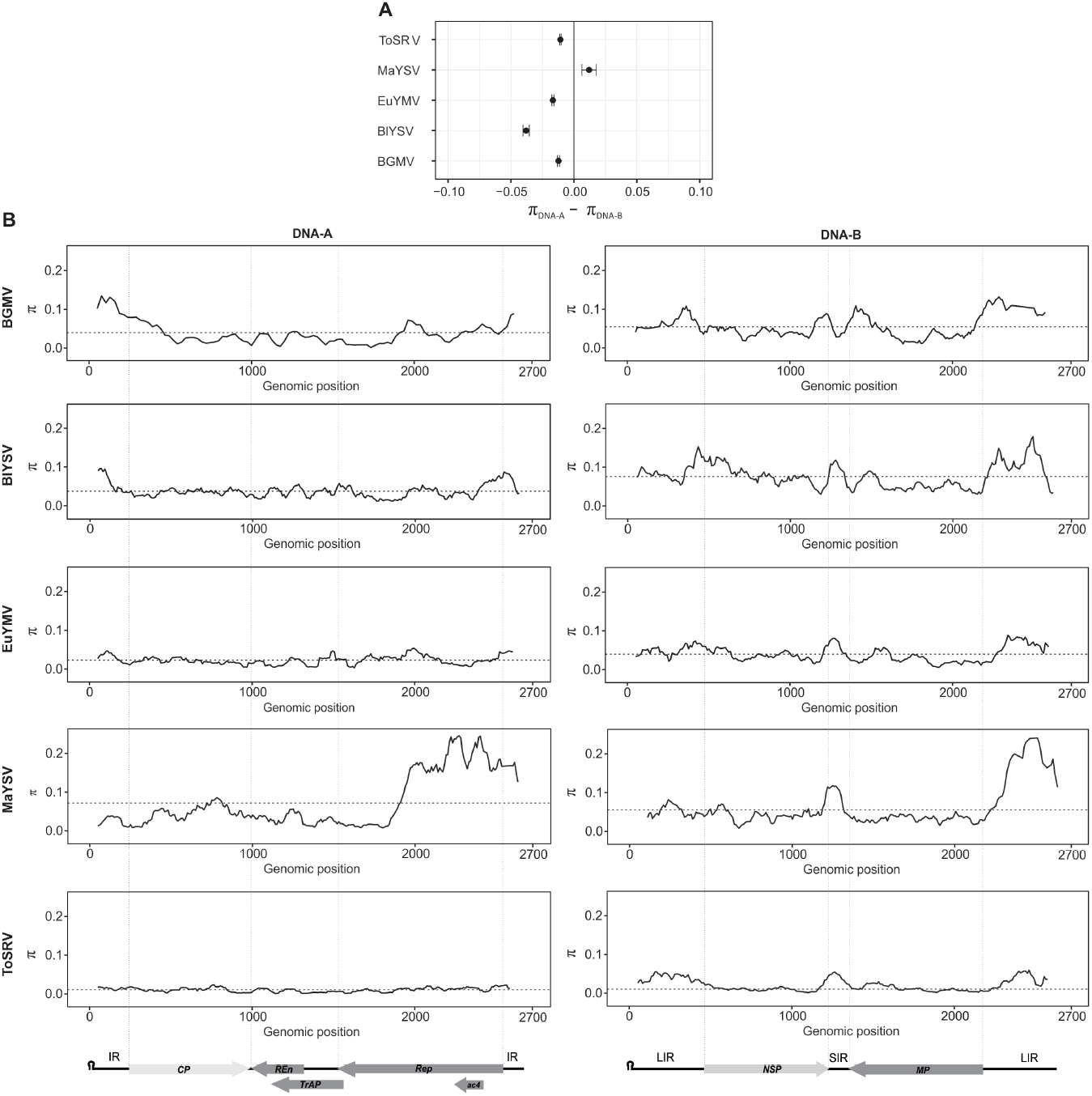
Genetic variability based on full-length nucleotide sequences of the DNA-A and DNA-B segments of *Bean golden mosaic virus* (BGMV), *Blainvillea yellow spot virus* (BlYSV), *Euphorbia yellow mosaic virus* (EuYMV), *Macroptilium yellow spot virus* (MaYSV) and *Tomato severe rugose virus* (ToSRV). (**A**) Statistical significance of the differences between the average pairwise number of nucleotide differences per site (nucleotide diversity - π) calculated for the DNA-A and DNA-B of each virus. Ninety-five percent bootstrap confidence intervals (CIs) for the difference between π values were estimated from 1,000 nonparametric simulations. Confidence intervals which include the value “zero” denote no statistically significant difference between the means. (**B**) Average pairwise number of nucleotide differences per site (nucleotide diversity - π) calculated on a 100-nucleotide sliding window with a step size of 10 nucleotides across the full-length nucleotide sequences of the DNA-A and DNA-B. Horizontal dotted lines represent the average pairwise number of nucleotide differences per site. Genome maps for the DNA-A and DNA-B are presented at the bottom.

The nucleotide diversity index was also calculated on a sliding window across the DNA-A and DNA-B for each species data set (Figure 4B). The results show an uneven distribution of variation along the components for almost all data sets. The exception was the ToSRV DNA-A (and to a lesser extent also its DNA-B), which in addition to presenting a very low degree of variation, this variation is evenly distributed along the genome (Figure 4B). To verify the statistical significance of the uneven distribution of variation along the genomic components, the genome was partitioned and nucleotide sequences of coding (*CP, Rep, TrAP, REn ac4, NSP* and *MP*) and intergenic regions (IR-A, LIR-B and SIR-B) were compared for each species data set (Suppl. Figure S4). Unsurprisingly, non-coding regions were more variable compared to coding regions (Suppl. Figure S4). The SIR-B was more variable than the LIR-B, which in turn was more variable than the IR-A. There was no consistent patttern for coding regions, with the most variable region being highly dependent on the data set analyzed (Suppl. Figure S4). The intragenic variation for nucleotide sequences of *CP*, *Rep*, *NSP* and *MP* and intraintergenic variation for IR-A and LIR-B were also analyzed (Suppl. Figure S5). As well as differences in the variation levels between genomic components and between regions within each component, there are clear differences in intragenic and intraintergenic variation levels (Suppl. Figure S5).

### Recombination analysis

It is well established that recombination plays an important role in the diversification and evolution of begomoviruses (43, 49, 50). However, most studies were based on analysis of non-segmented viruses or of the DNA-A component of bipartite viruses, and little is known about the effect of recombination on the DNA-B component (51). To investigate the effect of recombination on this component and possible differences in recombination patterns between the DNA-A and DNA-B, the intraspecific data sets were analyzed using RDP4.

There is a clear correlation between number of recombination events and genetic variability. BlYSV and MaYSV, which showed the higher number of events for both genomic components, are also the viruses with the higher genetic variability. Likewise, viruses with low genetic variability (BGMV, EuYMV and especially ToSRV) had few or no recombinantion events detected.

For all species data sets, the DNA-B was more prone to recombination than the DNA-A, with a higher number of unique, well-supported recombination events (detected by at least four methods and a *P*<0.001) (Figure 5A, B; Suppl. Table S3). Considering all unique events, 32 recombination events were detected in the DNA-B data sets and only 10 in the DNA-A data sets. No recombination events were detected for the DNA-A data sets from BGMV, EuYMV and ToSRV, whereas 3 and 7 unique events were detected for the DNA-A data sets from BlYSV and MaYSV, respectively (Figure 5A). The MaYSV DNA-B was the most prone to recombination, with 13 unique recombination events, followed by BlYSV, EuYMV, BGMV and ToSRV, with 9, 5, 4 and 1 unique events, respectively (Figure 5B). In DNA-A data sets the frequency of sequences that has at least one recombination event was higher for MaYSV (76%) followed by BlYSV (37%). In contrast, for the DNA-B the frequency of recombinant sequences was higher for BlYSV (98%), followed by MaYSV (75%) and EuYMV (40%). For BGMV and ToSRV DNA-B data sets, the frequency of recombinant sequences was similar (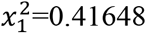, *P*=0.5187) even though it was extremely low, with only 4% for BGMV and 1.3% for ToSRV. These results demonstrate an asymmetric distribution of recombination events between genomic components, with a higher propensity of the DNA-B to recombine compared to the DNA-A, and also suggest that populations of begomoviruses from non-cultivated hosts are more prone to recombination, as shown for MaYSV and BlYSV and to a lesser extent for EuYMV DNA-B.

**Figure 5.**
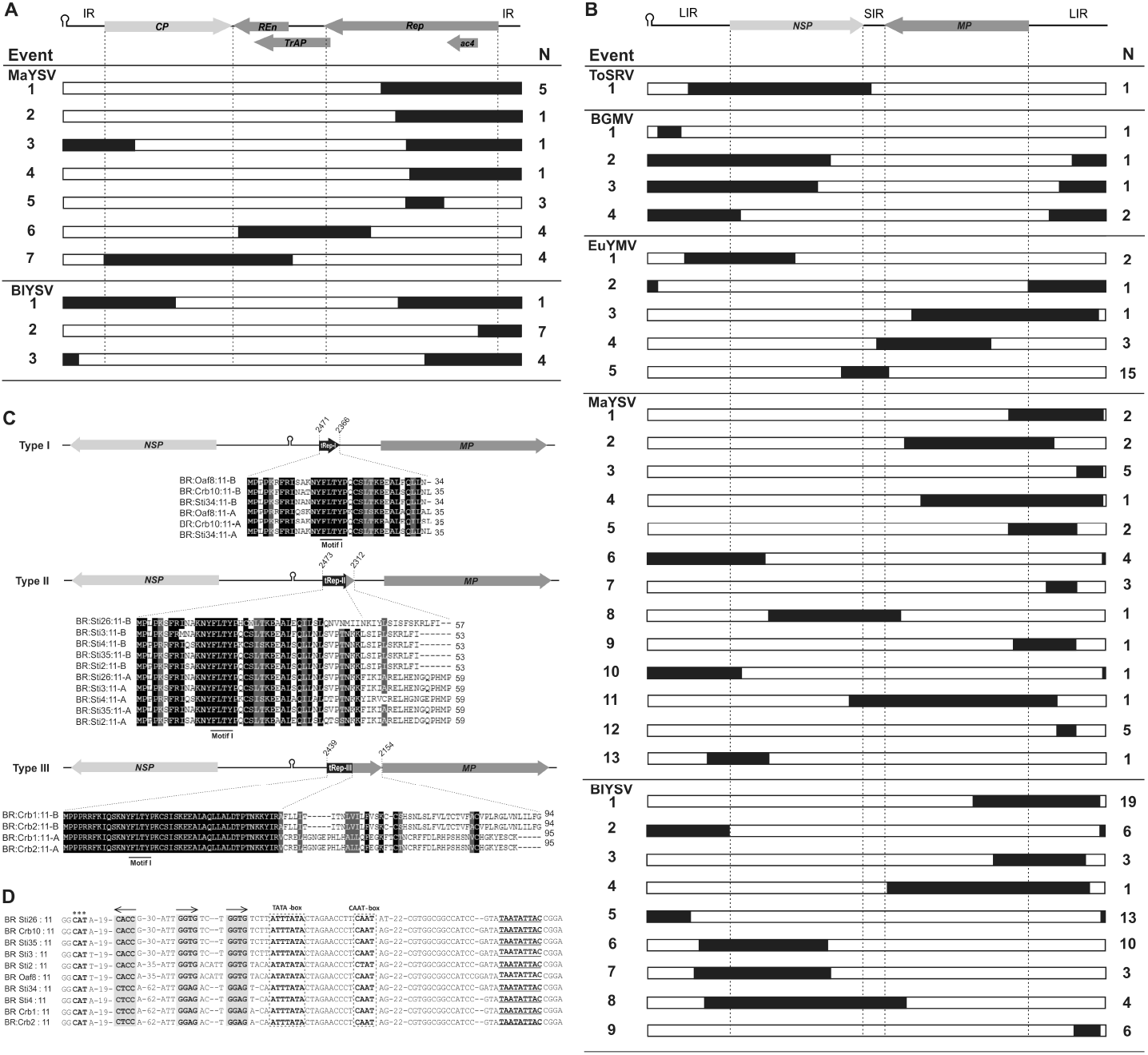
Recombination events detected in the (**A**) DNA-A and (**B**) DNA-B segments of begomoviruses using RDP4 (101). The genome map at the top of the figure corresponds to the schematic representations of the sequences below. Regions highlighted in black correspond to the donated (minor parent) portion, while the remaining portion corresponds to the receiving (major parent) sequence (see Suppl. Table S4 for more details on recombination events). IR, intergenic region in the DNA-A; LIR, large intergenic region and SIR, small intergenic region, both in the DNA-B. N, Number of isolates containing each event. (**C**) Truncated Rep (tRep) ORFs present in the DNA-B of *Macroptilium yellow spot virus* (MaYSV) isolates. For each type of tRep insertion (type I, type II and type III), the DNA-B is represented with genes indicated by arrows (*MP* in the viral sense, *NSP* in the complementary sense), and the portion of the tRep ORF filled in black represents homologous sequences to the N-terminal portion of the Rep protein in the cognate DNA-A. Each DNA-B map indicates the amino acid sequence alignments of the tRep ORF and the Rep protein N-terminal region of the cognate DNA-A (identical residues in black, similar residues in gray). (**D**) Nucleotide sequence alignment of the DNA-B LIR containing *cis*-acting elements. The nonanucleotide at the origin of replication is highlighted in bold and underlined. Putative Rep-binding elements (iterons) are shaded in gray and their orientation is indicated by arrows. Putative *cis*-acting elements found in eukaryotic promoters located in the LIR are marked by dotted boxes located at the right side of the tRep start codon (indicated in bold with asterisks).

Recombination breakpoints were mapped across of the DNA-A and DNA-B. Overall, a non-random distribution of recombination breakpoints was observed along both components. In agreement with previous studies (50), most recombination events detected in the DNA-A involved breakpoints located in the *Rep* gene (10 our of 20), followed by intergenic region (IR-A; 6/20). Conversely, few recombination breakpoints were located in the *CP* (2/20) and at the interface between the *TrAP* and *REn* genes (2/20). Most recombination events in the DNA-B have breakpoints in the LIR-B (37 out of 64), especially upstream to the initiation codon of the *MP* gene. In contrast, the SIR-B showed only 2/64 recombination breakpoints. This low number could be, at least in part, a consequence of its high nucleotide diversity, which impairs the capacity of RDP to detect reliable recombination events. Differently from the asymmetric breakpoint distribution observed for the *CP* and *REP* genes, the *NSP* and *MP* genes displayed a very homogeneous distribution of recombination breakpoints and a lower number of events compared to the intergenic regions (12/64 and 13/64 breakpoints, respectively). Despite approximately one third of the DNA-B length being comprised of intergenic regions, the number of breakpoints maped to intergenic and coding regions was similar (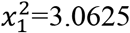, *P*=0.08012).

### Fragments of *Rep* ORFs in the large intergenic regions of the DNA-B (LIR-B) of MaYSV

During an ORF finder analysis to verify the integrity of ORFs encoded by the DNA-B, wes observed the presence of a large number of small ORFS in the LIR-B. To investigate the coding potential of these small ORFs, a BLASTn analysis was performed for all species data sets. Interestingly, small ORFs in the complementary-sense strand of the LIR-B of several MaYSV isolates are homologous to the N-terminal region of the *Rep* gene located in its DNA-A (Figure 5C; Table 3). These small ORFs were named truncated Rep (tRep) and, based on their common features, were grouped into three types named tRep I, II and III (Table 3). tRep ORFs were detected in 10 MaYSV isolates (42%) collected in three different regions (Table 3), ruling out the possibility of them being cloning artifacts. Despite their size ranging from 35 to 95 amino acids, the conserved Motif I (FLTYP) and iteron related domains were detected in all three types (Figure 5C). Interestingly, all tRep ORFs were located upstream of the start codon and downstream of the promoter of the *MP* gene (Figure 5D), suggesting that they may be transcribed and translated, and possibly interfering with expression of the *MP* gene. The ratio of non-synonymous to synonymous substitutions (dN/dS) for homologous regions (105 nt/35 aa) of tRep ORFs was 0.261, indicating negative selection acting in this region and suggesting that these ORFs, if expressed, are functional. Although a recombination analysis was not performed (it was not possible to obtain a good alignment between non-homologous components), the presence of these tRep ORFs in the LIR-B is a strong indication of inter-component recombination. An analysis of the LIR-B from the other four begomoviruses failed to identify any truncated ORFs homologous to the N-terminal region of the *Rep* gene as observed in MaYSV.

**Table 3.** Features of truncated Rep (tRep) ORFs located in the complementary-sense strand of the DNA-B large intergenic region (LIR-B) of *Macroptilium yellow spot virus* (MaYSV) isolates.

| Isolate | Type | Length of ORF (nt/aa) | Location in the genome* |  | Coverage (% nt/aa)<sup>#</sup> | Identity (% nt/aa) | E-value (nt/aa) |
| --- | --- | --- | --- | --- | --- | --- | --- |
|  |  |  | Start | End |  |  |  |
| BR:Oaf8:11 | I | 105/35 | 2471 | 2366 | 97/100 | 91/88 | 4e<sup>-31</sup>/6e<sup>-12</sup> |
| BR:Crb10:11 | I | 108/36 | 2472 | 2364 | 98/100 | 97/94 | 2e<sup>-41</sup>/1e<sup>-13</sup> |
| BR:Sti34:11 | I | 105/35 | 2468 | 2363 | 97/100 | 98/94 | 6e<sup>-41</sup>/1e<sup>-12</sup> |
| BR:Sti2:11 | II | 162/54 | 2473 | 2312 | 82/96 | 92/80 | 2e<sup>-44</sup>/4e<sup>-18</sup> |
| BR:Sti3:11 | II | 162/54 | 2470 | 2308 | 82/81 | 97/93 | 3e<sup>-54</sup>/9e<sup>-12</sup> |
| BR:Sti4:11 | II | 162/54 | 2473 | 2312 | 71/79 | 90/79 | 3e<sup>-34</sup>/7e<sup>-17</sup> |
| BR:Sti26:11 | II | 174/58 | 2486 | 2312 | 62/87 | 90/68 | 6e<sup>-32</sup>/3e<sup>-11</sup> |
| BR:Sti35:11 | II | 162/54 | 2470 | 2308 | 82/81 | 99/98 | 6e<sup>-57</sup>/5e<sup>-20</sup> |
| BR:Crb1:11 | III | 285/95 | 2439 | 2154 | 48/47 | 99/100 | 1e<sup>-61</sup>/1e<sup>-23</sup> |
| BR:Crb2:11 | III | 285/95 | 2439 | 2154 | 48/47 | 99/100 | 1e<sup>-61</sup>/1e<sup>-23</sup> |

\* Numbering starts at the first nucleotide after the cleavage site at the origin of replication and increases clockwise.
<sup>#</sup> Percentage of tRep ORF (nt, nucleotides; aa, amino acids) which is homologous to the Rep protein N-terminal region of the cognate DNA-A. Analyses performed using the BLASTn and BLASTp algorithms for nt and aa sequences, respectively.

### Reassortment analysis

As mentioned above for phylogenetic reconstruction, PACo analysis provided significant evidence for the global congruence between DNA-A and DNA-B trees (Table 1). However, it does not rule out the possibility of reassortment, since distance-based methods do not take into account the degree of congruence between links when accepting the hypothesis of global congruence between two trees (52). Indeed, the residual square sum (*m^2^_XY_*) values were variable across the different species data sets (Table 1), ranging from 0.1079 for ToSRV to 0.6039 for MaYSV, indicating that congruence levels are variable among different species data sets and suggesting that reassortment can occur at different frequencies across data sets.

To verify the occurrence of potential reassortment events, the individual contribution of each DNA-A and DNA-B link for the global square sum was assessed (Figure 6). This approach assumes that squared residuals between DNA-A and DNA-B links that are significantly high reflect phylogenetic incongruence, indicating the occurrence of reassortment. Analysis of the estimated jackknifed squared residuals for each link clearly indicated the occurrence of reassortment in all species data sets (Figure 6). Tanglegram plots depicting pairs of DNA-A and DNA-B phylogenetic trees also suggest the occurrence of reassortment in all data sets (Figure 7). Interestingly, some groups within the same species data set are apparently more prone to reassortment, as evidenced by a higher entanglement in corresponding clades of the phylogenies (Figure 7). To test this possibility, jackknifed squared residuals values were grouped according to sampling location and compared by estimating their 95% bootstrap confidence intervals from 1,000 nonparametric simulations. Indeed, some groups showed significantly higher residual values (Suppl. Figure S6), indicative of a higher contribution to incongruence and a higher frequency of reassortant genomes. For example, for the EuYMV data set, significantly higher residual values were observed for sequences sampled in Rio Grande do Sul and Mato Grosso do Sul compared to Góias and Paraná (Figure 6). Conversely, for the BlYSV data set there was no difference in residual values between sequences sampled at different locations, indicating no differences in reassortment frequency (Figure 6). Furthermore, most reassortment events occurred between isolates from the same or close regions.

**Figure 6.**
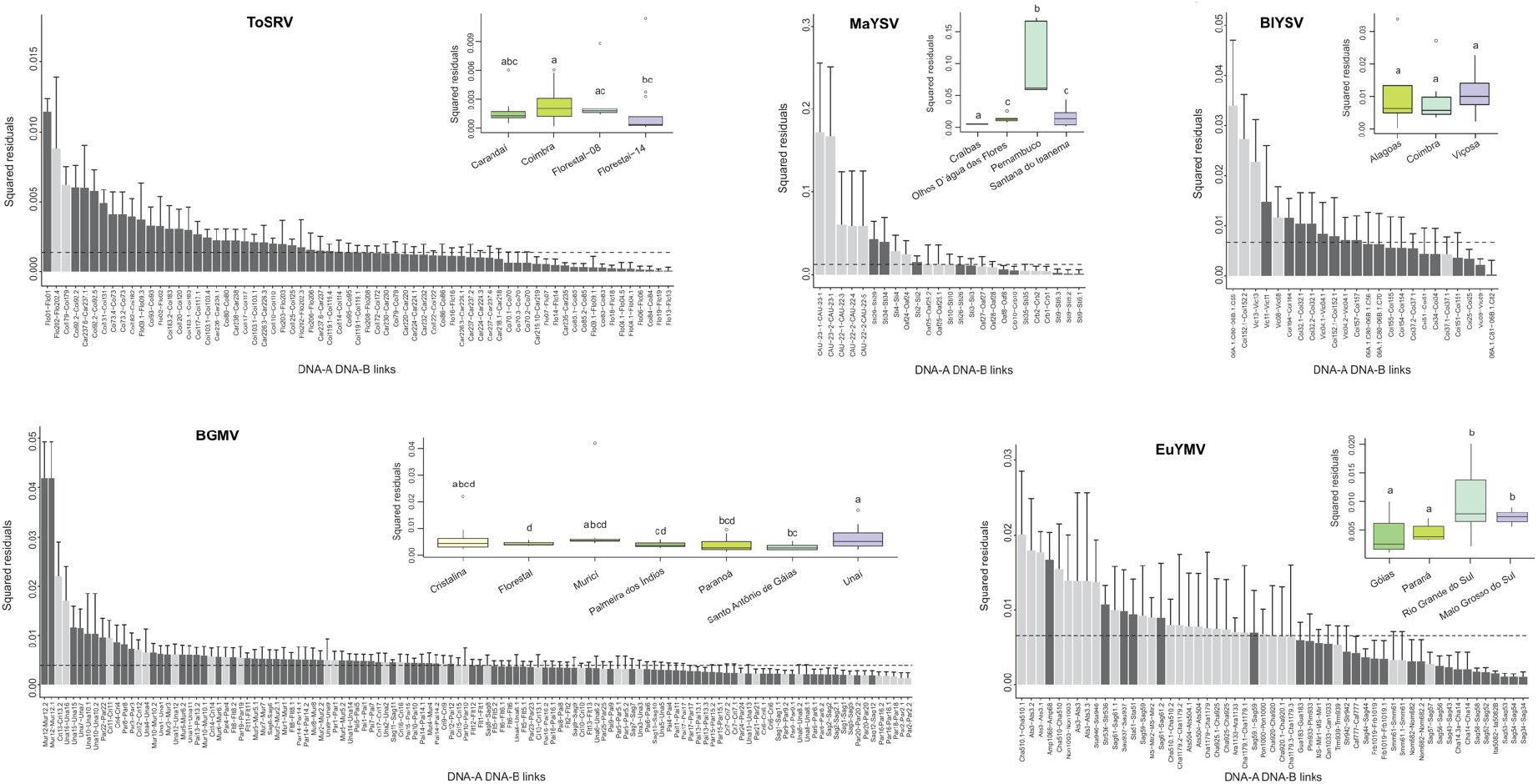
Evidence of reassortment events viewed through individual contributions of DNA-A-DNA-B links for the global congruence statistic. Estimated jackknifed squared residuals (bars) and 95% confidence intervals (error bars) for *Bean golden mosaic virus* (BGMV), *Blainvillea yellow spot virus* (BlYSV), *Euphorbia yellow mosaic virus* (EuYMV), *Macroptilium yellow spot virus* (MaYSV) and *Tomato severe rugose virus* (ToSRV). X-axis labels refer to isolate names for each pair of DNA-A and DNA-B segments. Horizontal dotted lines indicate the median squared residual values. Light gray bars correspond to sequences containing potential reassortment events detected by concatenated analysis in RDP. Box plots represent the jackknifed squared residuals grouped by sampling location. Ninety-five percent bootstrap confidence intervals (CIs) for the jackknifed squared residuals were estimated from 1,000 nonparametric simulations. Box plots with the same letter indicate that the confidence interval includes the value “zero”, denoting no statistically significant difference between means. The horizontal line inside the box corresponds to the median. This analysis was performed using data sets without recombinant blocks.

**Figure 7.**
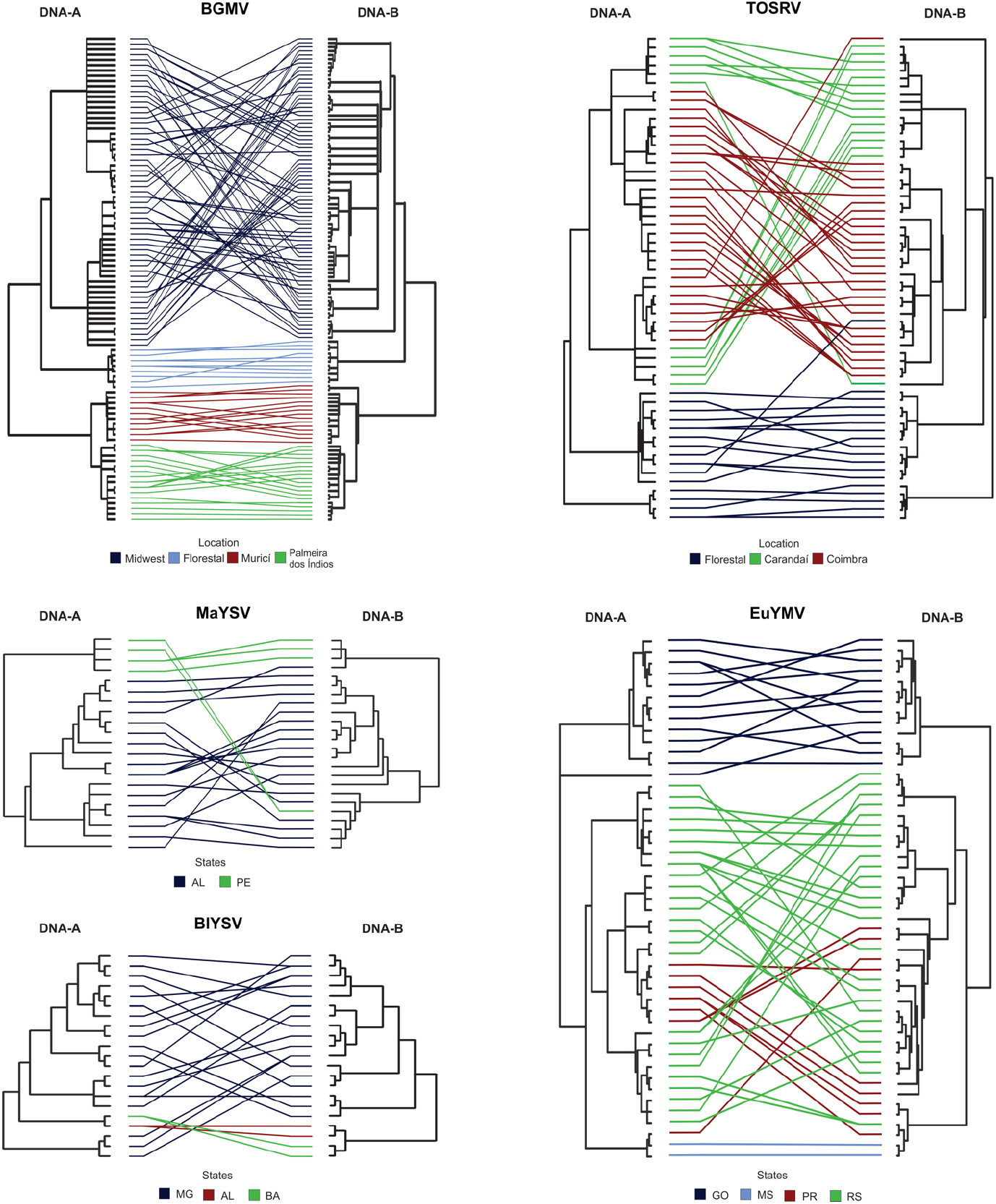
Tanglegram plots depicting phylogenetic relationships between inferred Bayesian phylogenetic trees of the DNA-A and DNA-B of *Bean golden mosaic virus* (BGMV), *Blainvillea yellow spot virus* (BlYSV), *Euphorbia yellow mosaic virus* (EuYMV), *Macroptilium yellow spot virus* (MaYSV) and *Tomato severe rugose virus* (ToSRV). The association among cognate DNA-A and DNA-B segments is showed as connecting lines colored according to sampling location. For BGMV, Midwest represents isolates sampled in Paranoá, Santo Antônio de Góias, Cristalina and Unaí. AL, Alagoas; BA, Bahia; GO, Góias; MS, Mato Grosso do Sul; PE, Pernambuco; PR, Paraná; RS, Rio Grande do Sul.

Analysis of concatenated sequences confirmed the results obtained by the topological congruence test. For almost all data sets, most of the reassortment events detected by RDP were supported by PACo analysis, displaying high squared residuals (Figure 6). The frequency of reassortment sequences detected by RDP, in agreement with PACo global test, was higher for MaYSV (19/25; 76%), followed by EuYMV (29/55; 53%), BGMV (35/117; 29%) and BlYSV (5/24; 21%) (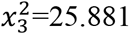, *P*=1.01×10^−05^; Table 4). For ToSRV only two sequences were detected as putatively reassortant by RDP. However, PACo analysis suggests a greater number of reassortment events, with 16/67 links (24%) showing squared residual values greater than two times the median value (indicative of strong incongruence). This discrepancy between RDP and PACo analyses may be due to the high sequence identity among isolates of the ToSRV data set (97-100% and 95-100% for DNA-A and DNA-B, respectively), impairing the capacity of RDP to detect reliable reassortment events.

**Table 4.** Reassortment events detected by RDP analysis of concatenated genomic components and proportion of reassortant sequences in *Bean golden mosaic virus* (BGMV), *Blainvillea yellow spot virus* (BlYSV), *Euphorbia yellow mosaic virus* (EuYMV), *Macroptilium yellow spot virus* (MaYSV) and *Tomato severe rugose virus* (ToSRV) data sets.

| **Virus** | **Event<sup>\*</sup>** | **Number of reassortant sequences by event/total number of sequences (%)** | **Number of reassortant sequences/total number of sequences (%)** |
| --- | --- | --- | --- |
| BGMV | 1 | 4/119 (3.36) | 35/119 (29.41) |
|  | 2 | 31/119 (26.05) |  |
| BIYSV | 1 | 1/24 (4.16) | 5/24 (20.83) |
|  | 2 | 1/24 (4.16) |  |
|  | 3 | 1/24 (4.16) |  |
|  | 4 | 2/24 (8.33) |  |
| EuYMV | 1 | 4/55 (7.28) | 29/55 (52.72) |
|  | 2 | 1/55 (1.82) |  |
|  | 3 | 3/55 (5.46) |  |
|  | 4 | 2/55 (3.63) |  |
|  | 5 | 5/55 (9.10) |  |
|  | 6 | 2/55 (3.63) |  |
|  | 7 | 3/55 (5.46) |  |
|  | 8 | 1/55 (1.82) |  |
|  | 9 | 1/55 (1.82) |  |
|  | 10 | 1/55 (1.82) |  |
|  | 11 | 5/55 (9.10) |  |
|  | 12 | 1/55 (1.82) |  |
| MaYSV | 1 | 9/25 (36.00) | 19/25 (76.00) |
|  | 2 | 10/25 (40.00) |  |
| ToSRV | 1 | 1/67 (1.49) | 2/67 (2.98) |
|  | 2 | 1/67 (1.49) |  |

<sup>\*</sup> For detailed information of the reassortment events see Suppl. Table S7.

Interestingly and surprisingly, when the BGMV and MaYSV data sets including recombinant blocks were analyzed, the global square sum was lower compared to the same data sets without recombinant blocks (Table 1; there was no difference in the case of the EuYMV and BlYSV data sets). Although the differences were small, this result suggests that recombination may somehow be restoring the congruence between phylogenies. To test the consistency of this result, the topological congruence between the phylogenetic trees in data sets with and without recombinant blocks was evaluated using a topology-based method, normalized PH85 distance (nPH85) (53). While for MaYSV the nPH85 distance was also lower for recombinant data set (nPH85_rec_=0.789 vs. nPH85_no_rec_=0.828), there was no difference for BGMV, EuYMV and BlYSV (data not shown).

### Selection analysis

To understand the possible effect of selection pressure on the uneven distribution of variation observed between DNA-A and DNA-B, the ratio of non-synonymous to synonymous substitutions (dN/dS) was compared for each gene/genomic component. The dN/dS ratios were <1 for almost all genes, except *ac4* (Table 5), indicating the predominance of negative selection acting on both components. In agreement with these results, a higher number of individual sites under negative selection was detected in all species data sets (Table 5). The *ac4* gene has a dN/dS ratio >1 for BGMV (2.53), EuYMV (1.15) and MaYSV (1.45) (Table 5).

**Table 5.** Non-synonymous to synonymous substitution ratios (dN/dS) and number of negatively and positively selected sites in each gene of the DNA-A and DNA-B segments of *Bean golden mosaic virus* (BGMV), *Blainvillea yellow spot virus* (BlYSV), *Euphorbia yellow mosaic virus* (EuYMV), *Macroptilium yellow spot virus* (MaYSV) and *Tomato severe rugose virus* (ToSRV).

| Virus | Gene | dN/dS | Negative selection |  |  |  | Positive selection |  |  |  |
| --- | --- | --- | --- | --- | --- | --- | --- | --- | --- | --- |
|  |  |  | SLAC | FEL | REL | Total | SLAC | FEL | REL | Total<sup>&</sup> |
| BGMV | CP | 0.1037 | 0 | 39 | 86 | 86 | 0 | 0 | 0 | 0 |
|  | Rep | 0.2490 | 8 | 41 | 3 | 41 | 0 | 0 | 2 | 2 |
|  | TrAP | 0.4007 | 0 | 6 | 4 | 10 | 0 | 0 | 0 | 0 |
|  | REn | 0.4261 | 1 | 6 | 1 | 6 | 0 | 0 | 0 | 0 |
|  | AC4 | 2.5300 | 0 | 3 | 3 | 3 | 0 | 1 | 0 | **1** |
|  | MP | 0.1183 | 20 | 67 | -* | 67 | 0 | 0 | -* | 0 |
|  | NSP | 0.2290 | 14 | 48 | 4 | 48 | 0 | 0 | 6 | 6 |
| BIYSV | CP | 0.0282 | 23 | 56 | 96 | 96 | 0 | 0 | 0 | 0 |
|  | Rep | 0.1837 | 21 | 59 | 27 | 67 | 0 | 2 | 5 | 5 |
|  | TrAP | 0.4378 | 5 | 13 | 2 | 13 | 0 | 1 | 10 | **10** |
|  | REn | 0.1902 | 7 | 15 | 33 | 33 | 0 | 0 | 0 | 0 |
|  | AC4 | 0.8575 | 1 | 4 | 12 | 12 | 0 | 0 | 0 | 0 |
|  | MP | 0.0667 | 63 | 103 | 47 | 104 | 0 | 1 | 7 | 7 |
|  | NSP | 0.1803 | 48 | 83 | 36 | 51 | 0 | 1 | 1 | 1 |
| EuYMV | CP | 0.0611 | 22 | 47 | -<sup>#</sup> | 47 | 0 | 0 | 0 | 0 |
|  | Rep | 0.2589 | 21 | 36 | 14 | 36 | 1 | 3 | 14 | 14 |
|  | TrAP | 0.6470 | 2 | 8 | 0 | 10 | 0 | 1 | 1 | 2 |
|  | REn | 0.2710 | 2 | 11 | 26 | 26 | 0 | 0 | 0 | 0 |
|  | AC4 | 1.1500 | 1 | 5 | 8 | 8 | 0 | 2 | 0 | **2** |
|  | MP | 0.0835 | 35 | 65 | 27 | 72 | 0 | 0 | 1 | 1 |
|  | NSP | 0.2284 | 28 | 56 | 8 | 56 | 0 | 1 | 3 | 3 |
| MaYSV | CP | 0.0943 | 11 | 41 | 78 | 78 | 0 | 0 | 0 | 0 |
|  | Rep | 0.2284 | 13 | 79 | 70 | 106 | 0 | 2 | 6 | 6 |
|  | TrAP | 0.5240 | 2 | 7 | 4 | 11 | 0 | 1 | 0 | 1 |
|  | REn | 0.3078 | 3 | 9 | 3 | 9 | 0 | 0 | 3 | 3 |
|  | AC4 | 1.4500 | 1 | 6 | 0 | 6 | 0 | 1 | 0 | 1 |
|  | MP | 0.0952 | 16 | 41 | 10 | 41 | 0 | 1 | 4 | 4 |
|  | NSP | 0.1983 | 11 | 38 | 8 | 38 | 0 | 2 | 12 | 12 |
| ToSRV | CP | 0.2610 | 2 | 13 | 0 | 13 | 0 | 0 | 6 | 6 |
|  | Rep | 0.1644 | 8 | 38 | 0 | 38 | 0 | 0 | 6 | 6 |
|  | TrAP | 0.5467 | 0 | 2 | 0 | 2 | 0 | 0 | 3 | 3 |
|  | REn | 0.3366 | 1 | 7 | 1 | 7 | 0 | 0 | 1 | 1 |
|  | AC4 | 0.2377 | 0 | 7 | 12 | 12 | 0 | 0 | 0 | 0 |
|  | MP | 0.1319 | 4 | 26 | 0 | 26 | 0 | 0 | 3 | 3 |
|  | NSP | 0.2311 | 5 | 22 | 2 | 22 | 0 | 0 | 0 | 0 |
| Total |  |  | 399 | 1107 | 625 | 1301 | 1 | 20 | 94 | 87 |

\* Analysis was not performed as the number of haplotypes was above the program's limit).
<sup>#</sup> No rates with dN>dS were inferred for this data set, suggesting that all sites are under negative selection.
<sup>&</sup> Numbers in bold indicate evidence of positive selection at $P<0.1$ according to PARRIS.

Although negative selection is predominant, dN/dS ratios were highly variable among genes/genomic components (0.028-2.530 for the DNA-A, 0.0667-0.2311 for the DNA-B) (Table 5). These results indicate that different genes/genomic components can evolve under distinct selection pressures and, unexpectedly, that DNA-B coding regions evolve under equal or more strict selection compared to DNA-A coding regions.

To compare the statistical significance of the differences in selection pressure, 95% confidence intervals were estimated for the mean dN/dS ratio and for the average number of negatively and positively selected sites (Figure 8). Considering data grouped by genomic component, the average dN/dS values were statistically lower for DNA-B (0.1562) compared to DNA-A (0.4778) (Figure 8A). In addition, the absolute number of individual sites under negative selection was statistically higher for the DNA-B (Figure 8B), suggesting a stronger negative selection acting on this component. However, these results should be treated with caution. Overlapping genes (which occur in the DNA-A but not in the BNA-B) may limit the accumulation of synonymous substitutions or a higher accumulation of non-synonymous substitution in one of the overlapping genes, leading to an increase in the dN/dS ratio (54 20326{Simon-Loriere, 2013 #21046)}. Thus, the higher dN/dS ratios in the DNA-A can be wrongly interpreted as evidence of positive selection or more relaxed evolution. Indeed, the *ac4* and *TrAP* genes, which have the higher dN/dS ratios (Figure 8A; Table 5), are both overlapping genes in the DNA-A. Therefore, the possibility of this result being an artifact cannot be excluded. When the dN/dS ratio was compared between DNA-A and DNA-B data set excluding the *ac4* and *TrAP* genes, there was no significant difference between dN/dS ratio (data not shown). Thus, it is suggested that DNA-B coding regions evolve at least under equivalent selection pressure as those of the DNA-A. Considering data grouped by genes, the dN/dS ratio was statistically higher for *ac4* and *TrAP* (Figure 8A). The *CP* and *MP* genes showed the lowest dN/dS ratios (Figure 8A). Moreover, *ac4* and *TrAP* (and also *REn*) showed a similar number of individual sites under negative selection, statistically lower than *CP, Rep, MP and NSP* (Figure 8B).

**Figure 8.**
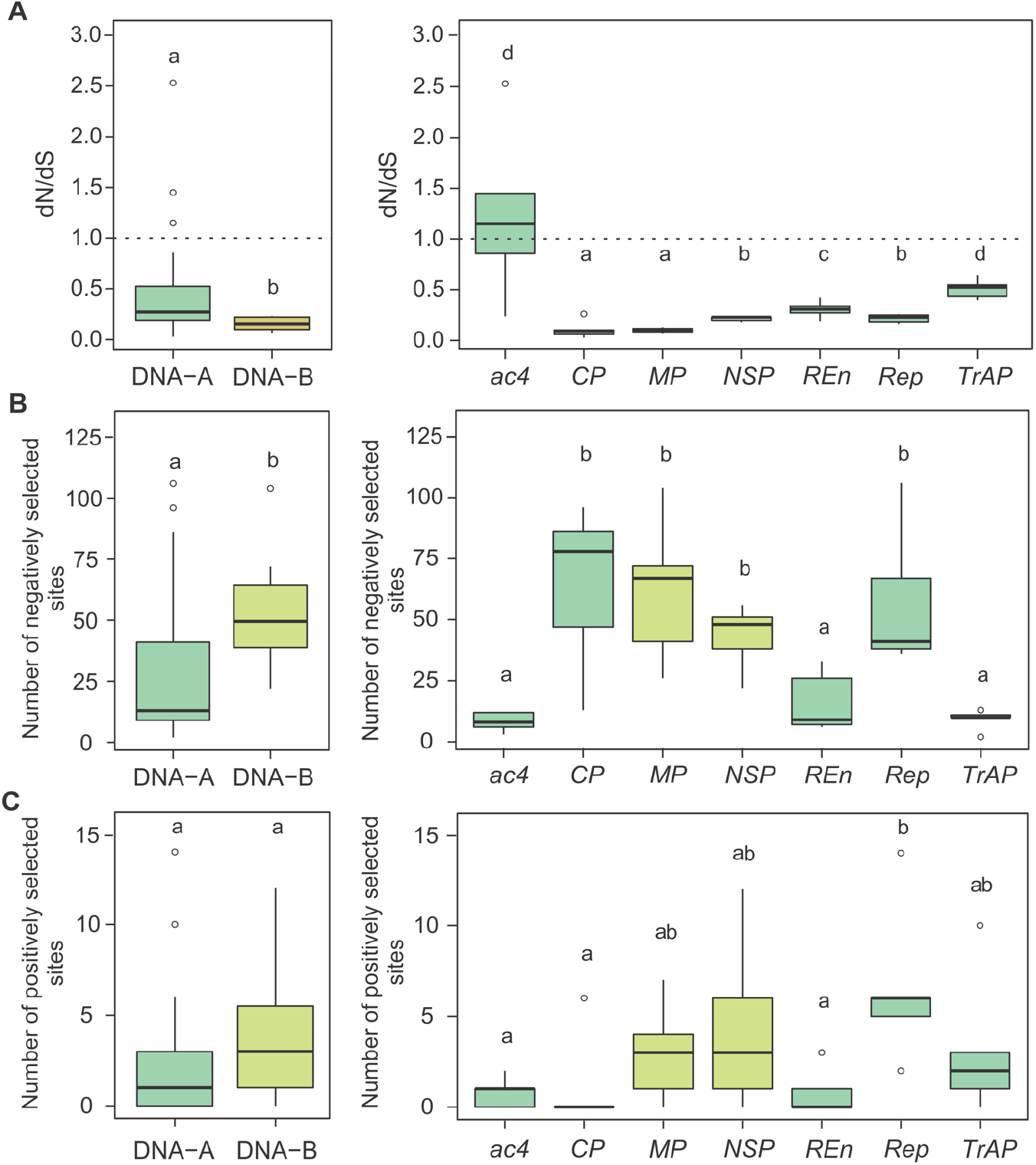
Statistical significance of the difference between selection pressure parameters calculated for DNA-A and DNA-B data sets, considering data grouped by genomic segment and coding region. Box-plots correspond to: **(A)** non-synonymous to synonymous substitution ratios (d*N*/d*S*); **(B)** the number of negatively selected sites; **(C)** the number of positively selected sites. Ninety-five percent bootstrap confidence intervals (CIs) for the difference between mean d*N*/d*S* values, number of negatively and positively selected sites were estimated from 1,000 nonparametric simulations. Box plots with the same letter indicate that the confidence interval includes the value “zero”, denoting no statistically significant difference between means. The horizontal line inside the box corresponds to the median.

The number of sites under positive selection was very low for both components, with no significant difference (Figure 8C). In agreement with the high dN/dS ratio of the *ac4* gene of BGMV and EuYMV (Table 5), we found evidence of positive selection by PARRIS (*P*<0.04), and one positively selected site for BGMV (codon 54) and two sites for EuYMV (codons 93 and 110) were detected by FEL. For *ac4* of MaYSV, which also exhibited dN/dS>1, only one positively selected site (codon 39) was detected by FEL, without evidence by PARRIS (Table 5). In addition, for BlYSV *TrAP* we found evidence of positive selection by PARRIS (*P*<0.043), and one (codon 90) and ten (codons 34, 79, 85, 88, 89, 90, 93, 95, 97, 128) positively selected sites were detected by FEL and REL, respectively. Other genes showed sites with weaker evidence of positive selection, detected by only one or two methods (Table 5).

## Discussion

Little is known about the significance of genome multipartition for viral biology and evolution (14, 17, 19, 23). Independence among segments, in addition to functional and structural differences, might heavily impact the evolutionary processes acting on the different segments, especially the generation and maintenance of diversity (22, 30, 55–58).

Several key events marked the evolution of geminiviruses, culminating with the emergence of segmented (bipartite) genomes in viruses classified in the genus *Begomovirus* (59). Expansion of the genome through the capture of a new component by an ancestral non-segmented virus, together with the loss of the *V2* gene [present only in Old World (OW) begomoviruses] are the main events related to the emergence of NW bipartite begomoviruses. Although NW begomoviruses are predominantly bipartite, very few studies have addressed evolutionary aspects of the complete (DNA-A and DNA-B) genome (31, 60).

Our results, based on analysis of five NW bipartite begomoviruses, indicate that DNA-A and DNA-B components respond differently to evolutionary processes, with the DNA-B being more permissive to variation and more prone to recombination than the DNA-A. Although the DNA-B is usually more variable than DNA-A at the population level, this is not an absolute rule, as MaYSV DNA-A is more variable than its DNA-B.

It has been proposed that the DNA-B can tolerate greater variation since it does not contain overlapping genes, while the DNA-A encodes four overlapping genes (*Rep/ac4* and *TrAP/Ren*) (31). With overlapping genes, a higher proportion of mutations will cause amino acid changes, leading to fitness trade-offs and negative selection. In addition, the proteins encoded by the DNA-A are involved in several *cis* and *trans* interactions which could be negatively affected by changes in their amino acid sequences, placing this component under stricter evolutionary constraints. However, using an interspecific data set, Ho *et al*. (61) demonstrated that while DNA-B coding regions of OW begomoviruses are indeed more variable than DNA-A coding regions, this pattern was not consistently observed for NW begomoviruses. In agreement with this obsevation, our results indicate that patterns of variation for NW begomovirus DNA components coding regions are species-specific. For example, there is no difference in variation among ToSRV *CP*, *Rep*, *MP* and *NSP* genes, although the DNA-B as a whole is more variable than the DNA-A. For MaYSV not only is the DNA-A more variable than the DNA-B, but *Rep* is the most variable gene followed by ac*4*, with *CP* more variable than *MP* and no difference between *CP* and *NSP*. In contrast, the BlYSV *MP* and *NSP* genes are its most variable genes. We also show that coding regions of the DNA-B evolve under at least the same selection pressure than DNA-A coding regions. This pattern might reflect the specialized movement function, encoded exclusively by the DNA-B in NW begomoviruses, impairing its ability to tolerate non-synonymous substitutions (62, 63). Conversely, bipartite OW begomoviruses, whose movement functions may partially be provided by DNA-A, can allow the DNA-B to evolve in a more relaxed fashion, being more permissive to variation (64–66). Moreover, NW DNA-B components accommodate additional functions besides movement, such as suppression of plant defense mechanisms (67), which may impose additional constraints.

In contrast to the ssDNA *Faba bean necrotic stunt virus* (FBNSV, family *Nanoviridae*), where there is no difference in variability between coding and non-coding regions (56), a clear difference was observed here, where intergenic regions (IRs) are more relaxed to variation than coding regions. Thus, in absolute terms, when the whole component is considered, the DNA-B may support a greater accumulation of variation compared to the DNA-A, since it has two IRs comprising approximately 1/3 of its length, in contrast to the DNA-A IR encompassing only 1/8 of its length. On this regard, our results show that the SIR-B is the most variable genome region, followed by the LIR-B and the IR-A. The SIR-B has no known *cis*-elements involved in replication or transcription, thus being more permissive to variation. On the contrary, the LIR-B and IR-A contain important *cis*-elements, therefore being under greater selective pressure. However, the LIR-B is almost three times longer than the IR-A, thus supporting a greater accumulation of variation without interference in regulatory *cis*-elements.

Asymmetric accumulation among the different genomic components in multipartite viruses has been shown, and seems to be a common trait shared by RNA and DNA viruses infecting animals and plants (22, 30, 68–70). During the infection cycle, each segment reaches a stable relative frequency, referred to as the “setpoint genome formula” (22, 70). Although very little is known about the biological and evolutionary meaning of this mechanism, it was proposed that it may be related to regulation of gene expression by controlling the copy number of each segment, and more interestingly, that it can directly modulate the evolutionary rates of the different segments (22, 58). Furthermore, it has been proposed that segments accumulating at higher frequency would have a greater mutation load, and consequently would evolve faster than other components present at a lower frequency (23, 30). Lozano *et al*. (30) suggested that the higher variability observed for the RNA-dependent RNA polymerase (RdRp) of *Tomato chlorosis virus* (ToCV; genus *Crinivirus*) is due to its location in the RNA1, which reaches a higher titer than the RNA2 and therefore accumulates a higher number of mutations in populational terms. A similar pattern was observed for the nanovirus FBNSV, where the high-titer DNA-N component is the most variable, whereas, DNA-R and DNA-S are the most conserved segments and accumulate at the lowest levels (22, 56). However, for other segments this relationship is not very clear, consistent with the joint action of other factors modulating the propensity to variation. Although no study has addressed whether the genome formula applies to bipartite begomoviruses, indirect evidence obtained using high-throughput sequencing from unamplified libraries suggest that the DNA-B can accumulate at twice the level of the DNA-A for both NW and OW begomoviruses (71; V.B. Pinto and F.M. Zerbini, *unpublished*). Considering that the two components are exposed to the same basal rate of mutation, a higher accumulation of the DNA-B could contribute to its higher variation at the populational level.

Viral populations are subject to severe bottlenecks, especially during systemic infection of the host and horizontal transmission by vectors (55, 58, 72, 73). After severe bottlenecks, a large reduction in the effective population size (*Ne*) can favor the action of genetic drift, leading to a dramatic loss of genetic diversity, as demonstrated for monopartite (72, 73) and tripartite (74) ssRNA viruses. The aphid-transmitted FBNSV (an octopartite ssDNA virus) also undergoes severe bottlenecks during vector transmission (58). Interestingly, the size of the bottleneck differs for the two FBNSV segments studied, and a direct correlation between *Ne* and the relative frequency of each segment was observed. These results suggest that genetic drift may differentially affect each genomic segment according to their relative frequencies, driving each one to distinct patterns of variation and evolution. It is reasonable to assume that differences in variability observed between begomovirus DNA-A and DNA-B components could at least in part be influenced by genetic drift, due to differences in the strength of the bottlenecks experienced by each component specially during transmission by *B. tabaci*.

Recombination is considered one of the main forces driving begomovirus evolution (41–43, 50, 75, 76). Our results showed a much greater propensity of the DNA-B to recombine compared to the DNA-A, with a higher number of unique events for DNA-B components compared to DNA-A components in all data sets. Martin *et al*. (77) demonstrated experimentally that tolerance to recombination of a given region of the genome is correlated to the degree of divergence between the segments exchanged and the number of inter- and intra-genomic interactions established. Thus, the greater propensity of the DNA-B to recombine may be explained by its lowest organizational complexity (only two non-overlapping genes, compared to one non-overlapping and four overlapping genes in the DNA-A) and the lowest number of inter- and intragenomic interactions of its encoded proteins (78).

In contrast to our results, previous studies analyzing the evolutionary dynamics of cassava mosaic geminiviruses (CMG) reported a much smaller number of unique recombinations events for the DNA-B compared to the DNA-A (79, 80). However, these results were based on the analysis of interspecific data sets with a higher number of DNA-A than DNA-B sequences. Since OW DNA-B components have a higher degree of variation compared to the DNA-A (as discussed above), obtaining a reliable interspecific alignment may be difficult, impairing the capacity of RDP to detect reliable recombination events. In addition, sampling more sequences might increase the likelihood of parental and recombinant sequences being sampled together, increasing the number of events detected.

Rodelo-Urrego *et al*. (60) observed no difference in the frequency of recombinant sequences between DNA-A and DNA-B components of *Pepper golden mosaic virus* (PepGMV) and *Pepper huasteco yellow vein virus* (PHYVV), two bipartite begomoviruses infecting chiltepin (*Capsicum annuum* var. *glabriusculum*) plants in Mexico. We found equivalent recombination frequencies for the DNA-A and DNA-B of MaYSV, however, the number of unique events was much higher for the DNA-B compared to the DNA-A. For BlYSV, both the number of unique events and the frequency of recombinant sequences were higher for the DNA-B, with 41% of the isolates carrying the same recombination event. Thus, the frequency of recombinant sequences, more than the propensity to recombination, may reflect the amplification of a few successful events in the population which are positively selected. Considering the five species, our results indicate a higher propensity for recombination in the DNA-B compared to the DNA-A, but also that selective advantage of specific recombination events may vary across DNA components and viruses.

Although less frequent, intercomponent homologous recombination involving the exchange of replication origin among heterologous components in multipartite viruses has been previously reported (29, 32, 81, 82). Interestingly, Gregorio-Jorge *et al*. (83) reported the presence of small fragments of *Rep* gene-derived sequences in the DNA-B intergenic region of several begomoviruses from NW. These *Rep* fragments, ranging from 35 to 51 nt in length, were located in the viral-sense strand and it was suggested that they could be involved in the posttranscriptional regulation of the cognate *Rep* gene. This suggests that non-homologous recombination among heterologous components may also occur. We detected small truncated Rep (tRep) ORFs in the complementary-sense strand of the LIR-B of several MaYSV isolates, ranging from 35 to 95 amino acids (significantly longer than the short regions identified by Gregorio-Jorge *et al*. (83)). The presence of these tRep ORFs is a strong indication of intercomponent recombination. The tRep ORFs were located downstream of the *MP* gene promoter, suggesting that they can be expressed. Moreover, they include the coding region for Motif I and the IRD, which are involved in the process of recognition and binding of Rep to the origin of replication (39). If the tRep ORFs are expressed, one possibility is that the truncated proteins can compete with Rep for binding at the origin of replication, negatively interfering in the replication process. Another possibility is that tRep ORFs might interfere with the translation of MP, due to their location upstream of the MP initiation codon. However, these possibilities need to be experimentally tested. Although non-homologous recombination may be deleterious due to the breakage of important coding or regulatory regions, these results suggest that it can occur in a less congested component such as the DNA-B and might accommodate the emergence of new functions, as suggested above.

While the cophylogenetic analysis suggests a global congruence between DNA-A and DNA-B trees across all data sets, congruence levels were variable among them, consistent with the role of reassortment in the evolution of multipartite viruses (84, 85). We were not able to estimate and compare evolutionary rates between DNA-A and DNA-B data sets, since the sequences were sampled on too short a time span. However, the phylogenetic concordance observed, which would indicate a co-evolution scenario, may be caused by biogeographic processes and limited gene flow among subpopulations, rather than similar evolutionary rates driven by mutual selection between the components. Indeed, DNA-A and DNA-B phylogenies depict the formation of large clades according to geographical origin, providing evidence of population subdivision based on geography, as confirmed by DAPC analysis. Structural differences (mainly the presence of overlapping genes in the DNA-A) and the specialized functions of each component would argue against the assumption of similar evolutionary rates, but nevertheless it needs to be tested. In spite of the selection analysis, which suggests that coding regions from each component may evolve under at least the same selection pressure (which could lead to similar evolutionary rates), different segments in multipartite viruses may be differentially influenced by recombination and genetic drift (58). That could overcome, to some extent, the homogenizing effect of selection and contribute to distinct rates. In addition, although molecular clock-based methods to estimate substitution rates consider evolution to be driven exclusively by mutational dynamics, recombination constitutes an important mechanism that may speed up begomovirus evolution (50, 75). Differences in the propensity for recombination between DNA-A and DNA-B may drastically affect their evolution, guiding each component to distinct patterns of variation. Therefore, even with DNA-A and DNA-B working together as a single functional unit, with both components necessary to systemically infect the plant, only those regulatory elements required to maintain the integrity and functionality of genome may show similar evolutionary rates driven by reciprocal selection, unlike the remaining of the genome.

Previous studies, based only on the DNA-A analysis, have shown that begomoviruses populations segregate based on geographical location on a global as well as on a local scale (41, 44, 48, 86). Our results showed this to occur for the DNA-B as well. Although a direct comparison among different data sets should be treated with caution, due to differences in geographic scale and number of sequences sampled, our results demonstrate that geography-based subdivision is clear for both components of BGMV, EuYMV and ToSRV, and to a lesser extent for the DNA-A of MaYSV. For BlYSV, a consistent clustering pattern was not verified by either phylogenetic or DAPC analyses, even analyzing isolates sampled in distant geographical regions. This result suggests an incipient pattern of differentiation, probably due a connection between the two sampled regions, allowing gene flow between them or a recent founder effect.

Slight differences in the genetic structure between DNA-A and DNA-B were observed for BGMV, EuYMV and ToSRV, consistent a role of reassortment. Indeed, we were able to detect reassortment events across all data sets using two different methods. However, all events detected were intraspecific and most exchanged components were restricted among geographically and genetically close subpopulations, probably contributing little for the genetic makeup of the subpopulation. These results may reflect a limited dispersion capacity of the virus, which is primarily dependent on the vector. If this is the case, the genetic structure of begomovirus population would match the genetic structure of *B. tabaci* populations. Thus, studies addressing the genetic structure of both the virus and its vector might provide insights on the ecological and genetic processes that affect begomovirus populations.

The extent of evolutionary congruence among distinct genomic components of segmented and multipartite viruses depends on the interplay between evolutionary and biological process acting at both the inter- and intragenomic levels. Thus, different structural and functional constraints along genomic regions might drive such regions and even whole segments to distinct patterns of evolution. Although it is expected that different segments will follow a similar evolutionary way, given the functional dependence among segments, previous studies showed that different components in multipartite viruses might experiment distinct evolutionary routes (28–32, 85). Our results demonstrate the DNA-A and DNA-B segments of NW begomoviruses, as well as different regions in a gene and genomic segment, display different evolutionary patterns, with significant differences in variation and recombination levels. Thus, while regulatory regions responsible for maintaining the integrity and functionality of the multipartite genome might be under strong selection pressure in different segments, most of the genome may evolve in a more relaxed fashion, driving distinct patterns of variation. This more relaxed evolution may be an advantage of multipartition, as each component may behave as a single independent entity, exploring a larger portion of sequence space and allowing for faster adaptation to a constantly changing environment.

## Methods

### Cloning and sequencing of DNA-B components

Begomovirus DNA-B clones were obtained from total DNA extracted from the samples collected for the studies of Lima *et al*. (42), Rocha *et al*. (41) and Ramos-Sobrinho *et al*. (48), which have been stored in our laboratory at −80°C. Total DNA was used as a template for rolling-circle amplification (RCA) of viral genomes as described by Inoue-Nagata *et al*. (87). To facilitate the cloning of DNA-B components, restriction analysis of the previously cloned DNA-As was performed using Ape 2.0 Plasmid Editor, and only restriction enzymes that do not cleave the cognate DNA-A component were chosen. Unit genome-length fragments (approximately 2,600 nucleotides) were excised and ligated into the pBLUESCRIPT-KS+ (pKS+) plasmid vector (Stratagene), previously cleaved with the same enzyme. Viral inserts were sequenced commercially (Macrogen Inc.) by primer walking. Full-length begomovirus genomes were assembled using Geneious v. 8.1 (88). Sequences were initially analyzed with the BLAST*n* algorithm (89) to confirm that they corresponded to a DNA-B. The genomic organization and integrity of the open reading frames (ORF) were checked using the Fangorn Forest method (90) implemented in Geminivirus Data Warehouse (91). Sequences with truncated ORFs or with atypical DNA-B organization were discarded.

### Multiple sequence alignments and phylogenetic analysis

All genome sequences were organized to begin at the nicking site in the invariant nonanucleotide at the origin of replication (5’-TAATATT//AC-3’). Multiple sequence alignments were prepared for the full-length nucleotide sequences of DNA-A and DNA-B, of the *CP* (capsid protein), *Rep* (replication-associated protein), *REn* (replication enhancer protein), *TrAP* (trans-activating protein), *AC4*, *MP* (movement protein) and *NSP* (nuclear shuttle protein) genes, and of the DNA-A intergenic region (IR-A), DNA-B large intergenic region (LIR-B) and DNA-B small intergenic region (SIR-B), using the MUSCLE option in MEGA6 (92). Alignments were manually checked and adjusted when necessary. The same alignments were used for all subsequently performed analyzes.

Phylogenetic trees were constructed based on the full-length DNA-A and DNA-B nucleotide sequence using Bayesian inference performed with MrBayes v. 3.0b4 (93). The program MrModeltest v. 2.2 (94) was used to select the nucleotide substitution model with the best fit for each data set in the Akaike Information Criterion (AIC). The analyses were carried out running 10,000,000 generations and excluding the first 2,500,000 generations as burn-in. Trees were visualized and edited using FigTree (tree.bio.ed.ac.uk/software/figtree/). For data sets that showed evidence of recombination, trees with and without recombinant blocks were generated. The regions corresponding to the recombinant blocks were replaced by missing data (?), using a script (available from the authors upon request) written in R software (95 8112).

### Variability indices and genetic structure

The average pairwise number of nucleotide differences per site (nucleotide diversity, π) was calculated using a script (available from the authors upon request) written in R software. The statistical significance of the differences amongst the mean π obtained from different data sets was calculated by estimating their 95% bootstrap confidence intervals from 1,000 nonparametric simulations using the simpleboot statistical package in R software (96) as previously described (43). This method was chosen because it does not make assumptions about the distribution of data and allows comparing data sets with different sample sizes (43). Nucleotide diversity was also calculated on a 100-nucleotide sliding window with a step size of 10 nucleotides across the full-length DNA-A and DNA-B nucleotide sequence for each data set. The number of segregating sites (S), number of haplotypes (H) and haplotype diversity (Hd) were estimated for each data set using DnaSP v. 5.10 (97).

Inferences about population genetic structure was performed using Discriminant Analysis of Principal Components (DAPC) (98), a nonparametric multivariate static implemented in the adegenet package (99) in R software. DAPC is a model-free method, with no assumptions about Hardy-Weinberg equilibrium or linkage disequilibrium, thus being ideal for viruses. For the full-length DNA-A and DNA-B, the number of genetic clusters (*k*) were pre-defined using a *k*-means algorithm. To find the best number of genetic clusters to describe the data, the *k*-means was run sequentially with values of *k* varying from 2 to 10, retaining all principal components (PC). The choice of the best clustering model was based primarily on the minimum number of *k* after which the Bayesian Information Criterion (BIC) decreases by an insignificant amount, as proposed by Jombart *et al*. (98). Since there is not always a clear limit at which the BIC value decreases by an insignificant amount, the percentage of variation among genetic clusters inferred by *k*-means was plotted with BIC according to analysis of molecular variance (AMOVA) performed with the Poppr package (100) in R software. After assigning individuals to each inferred genetic cluster, the optimum PC values to run discriminant analysis were investigated through visual analysis of the cumulative variance graph explained by PC and a-score optimization. In addition, to support and verify the degree of differentiation between genetic clusters inferred by DAPC, the coefficient of nucleotide differentiation (Nst) was estimated using DnaSP v. 5 (97).

Subpopulation will be used here to define a group of individuals sampled at the same location without previous knowledge of population structure, and genetic cluster as a set of individuals genetically similar defined based on the DAPC analysis.

### Recombination and reassortment analysis

The occurrence of recombination was analyzed for the DNA-A and DNA-B separately using intraspecific data sets. Recombination analysis was performed using the Rdp, Geneconv, Bootscan, Maximum Chi Square, Chimaera, SisterScan and 3Seq methods implemented in Recombination Detection Program (RDP) v. 4 (101). Alignments were scanned with default settings for the different methods. Statistical significance was inferred by *P*-values lower than a Bonferroni-corrected cut-off of 0.05. Only recombination events detected by at least four different methods were considered reliable.

To assess the occurrence of reassortment (or pseudorecombination), the topological congruence between the inferred Bayesian phylogenetic trees based on full-length nucleotide sequences of the DNA-A and DNA-B were analyzed with Procrustean Approach to Cophylogeny (PACo). PACo is a distance-based method which assesses the global congruence between trees and the contribution of each individual DNA-A and DNA-B association to the global congruence statistic (102). The method uses matrices of patristic distances calculated from inferred Bayesian DNA-A and DNA-B phylogenies, testing the null hypothesis of no congruence between trees, a through permutation test. Matrices of patristic distances from DNA-A and DNA-B phylogenies were calculated using the ape package (103) and transformed into principle coordinates (PCo) using the paco package (104), both in R software. The PCo and DNA-A-DNA-B association matrices were used for procrustean superimposition analysis to produce the residual square sum (*m^2^_XY_*), which is inversely proportional to the level of global congruence between the phylogenies. The contribution of each individual DNA-A and DNA-B link for the global congruence and 95% confidence intervals were estimated using a jackknife method. Individual links that highly contribute to global congruence have a small contribution to the global square sum. The statistical significance for global test was assessed with 999 permutations in R software using the paco package (104). The assumption behind this approach is that incongruence between DNA-A and DNA-B phylogenies is due to reassortment events. Since recombination can also lead to incongruence between phylogenies, analyzes were performed in data sets with and without recombinant blocks. In addition, reassortment was analyzed by concatenating the sequences of the DNA-A and DNA-B and scanning sequences for recombination breakpoints near the artificial joint between the two genomic components using RDP as described previously.

### Selection analysis

Potentials sites under positive and negative selection in coding regions of the *CP*, *Rep*, *TrAP, REn, AC4, MP* and *NSP* genes were identified using four distinct methods: single likelihood ancestor counting (SLAC), fixed effects likelihood (FEL), random effects likelihood (REL) (105) and partitioning for robust inference of selection (PARRIS) (106), implemented in the DataMonkey webserver (107). The mean ratios of non-synonymous to synonymous substitutions (d*N*/d*S*) were estimated for all genes using the SLAC method. All methods were applied using nucleotide substitution models with the best fit for each data set determined in the Datamonkey webserver. To avoid misleading selection results, we searched for recombination breakpoints in each data set using Genetic Algorithm Recombination Detection (GARD) (108). All analyses were based on the inferred GARD-corrected phylogenetic trees. To access the statistical significance of the differences in selection pressure, 95% confidence intervals were estimated for the mean dN/dS ratio and for the average number of negatively and positively selected sites, from 1,000 nonparametric simulations in R software using the Simpleboot statistical package.

## Acknowledgements

This work was funded by CAPES (Finance Code 001), CNPq (grant 409599/2016-6 to FMZ) and FAPEMIG (grant APQ-03444-16 to FMZ). The authors wish to thank Eduardo S.G. Mizubuti and Siobain Duffy for helpful discussions, and Fernando García-Arenal for critical review of the manuscript.

## Notes

### Competing Interest Statement

The authors have declared no competing interest.

