## Supplementary Tables S1-S7; Supplementary Figures S1-S5 for "Evolutionary dynamics of bipartite begomoviruses revealed by complete genome analysis"

**Supplementary Table S1.** Begomovirus isolates obtained in this study.

| Sample code | Date of collection | Location | Geographical coordinates | | Host | Enzyme | Isolate name | GenBank access number |
|---|---|---|---|---|---|---|---|---|
| *Bean golden mosaic virus* (BGMV) | | | | | | | | |
| MG-146 | June, 2012 | Unaí, MG | S16°43'02.2" | W046°45'14" | *Phaseolus vulgaris* | *Spe*I | BR:Una1:12 | MT626860 |
| MG-147 | June, 2012 | Unaí, MG | S16°43'02.2" | W046°45'14" | *Phaseolus vulgaris* | *Spe*I | BR:Una2:12 | MT626861 |
| MG-148 | June, 2012 | Unaí, MG | S16°43'02.2" | W046°45'14" | *Phaseolus vulgaris* | *Spe*I | BR:Una3:12 | MT626862 |
| MG-149 | June, 2012 | Unaí, MG | S16°43'02.2" | W046°45'14" | *Phaseolus vulgaris* | *Spe*I | BR:Una4:12 | MT626863 |
| MG-150 | June, 2012 | Unaí, MG | S16°43'02.2" | W046°45'14" | *Phaseolus vulgaris* | *Spe*I | BR:Una5:12 | MT626864 |
| MG-152 | June, 2012 | Unaí, MG | S16°43'02.2" | W046°45'14" | *Phaseolus vulgaris* | *Spe*I | BR:Una6.1:12 | MT626865 |
| | | | | | | *Spe*I | BR:Una6.2:12 | MT626866 |
| MG-153 | June, 2012 | Unaí, MG | S16°43'02.2" | W046°45'14" | *Phaseolus vulgaris* | *Spe*I | BR:Una7:12 | MT626867 |
| MG-154 | June, 2012 | Unaí, MG | S16°43'02.2" | W046°45'14" | *Phaseolus vulgaris* | *Spe*I | BR:Una8.1:12 | MT626868 |
| | | | | | | *Spe*I | BR:Una8.2:12 | MT626869 |
| MG-155 | June, 2012 | Unaí, MG | S16°43'02.2" | W046°45'14" | *Phaseolus vulgaris* | *Spe*I | BR:Una9:12 | MT626870 |
| MG-156 | June, 2012 | Unaí, MG | S16°43'02.2" | W046°45'14" | *Phaseolus vulgaris* | *Spe*I | BR:Una10.1:12 | MT626871 |
| | | | | | | *Spe*I | BR:Una10.2:12 | MT626872 |
| MG-157 | June, 2012 | Unaí, MG | S16°43'02.2" | W046°45'14" | *Phaseolus vulgaris* | *Spe*I | BR:Una11:12 | MT626873 |
| MG-158 | June, 2012 | Unaí, MG | S16°43'02.2" | W046°45'14" | *Phaseolus vulgaris* | *Spe*I | BR:Una12:12 | MT626874 |
| MG-161 | June, 2012 | Unaí, MG | S16°43'02.2" | W046°45'14" | *Phaseolus vulgaris* | *Spe*I | BR:Una13:12 | MT626875 |
| MG-162 | June, 2012 | Unaí, MG | S16°43'02.2" | W046°45'14" | *Phaseolus vulgaris* | *Spe*I | BR:Una14:12 | MT626876 |
| MG-163 | June, 2012 | Unaí, MG | S16°43'02.2" | W046°45'14" | *Phaseolus vulgaris* | *Spe*I | BR:Una15:12 | MT626877 |
| MG-165 | June, 2012 | Unaí, MG | S16°43'02.2" | W046°45'14" | *Phaseolus vulgaris* | *Spe*I | BR:Una16:12 | MT626878 |
| GO-2 | June, 2012 | Santo Antônio de Góias, GO | S16°30'22.5" | W049°17'06.8" | *Phaseolus vulgaris* | *Spe*I | BR:Sag11:12 | MT626917 |
| GO-3 | June, 2012 | Santo Antônio de Góias, GO | S16°30'22.5" | W049°17'06.8" | *Phaseolus vulgaris* | *Spe*I | BR:Sag1.1:12 | MT626907 |
| | | | | | | *Spe*I | BR:Sag1.2:12 | MT626908 |
| GO-4 | June, 2012 | Santo Antônio de Góias, GO | S16°30'22.5" | W049°17'06.8" | *Phaseolus vulgaris* | *Spe*I | BR:Sag2:12 | MT626909 |
| | | | | | | *Spe*I | BR:Sag2.1:12 | MT626910 |
| GO-9 | June, 2012 | Santo Antônio de Góias, GO | S16°30'22.5" | W049°17'06.8" | *Phaseolus vulgaris* | *Spe*I | BR:Sag7:12 | MT626913 |
| GO-15 | June, 2012 | Santo Antônio de Góias, GO | S16°30'22.5" | W049°17'06.8" | *Phaseolus vulgaris* | *Spe*I | BR:Sag5:12 | MT626911 |
| GO-22 | June, 2012 | Santo Antônio de Góias, GO | S16°30'22.5" | W049°17'06.8" | *Phaseolus vulgaris* | *Spe*I | BR:Sag6:12 | MT626912 |
| GO-25 | June, 2012 | Santo Antônio de Góias, GO | S16°30'22.5" | W049°17'06.8" | *Phaseolus vulgaris* | *Spe*I | BR:Sag12:12 | MT626918 |
| GO-26 | June, 2013 | Santo Antônio de Góias, GO | S16°30'22.5" | W049°17'06.8" | *Phaseolus vulgaris* | *Spe*I | BR:Sag8:12 | MT626914 |
| GO-27 | June, 2012 | Santo Antônio de Góias, GO | S16°30'22.5" | W049°17'06.8" | *Phaseolus vulgaris* | *Spe*I | BR:Sag9:12 | MT626915 |
| GO-29 | June, 2012 | Santo Antônio de Góias, GO | S16°30'22.5" | W049°17'06.8" | *Phaseolus vulgaris* | *Spe*I | BR:Sag10:12 | MT626916 |

| | | | | | | | | |
|---|---|---|---|---|---|---|---|---|
| GO-63 | June, 2012 | Paranoá, DF | S16°00'47" | W047°33'21.5" | *Phaseolus vulgaris* | *Spe*I | BR:Par14.1:12 | MT626932 |
| | | | | | | *Hind*III | BR:Par14.2:12 | MT626933 |
| GO-64 | June, 2012 | Paranoá, DF | S16°00'47" | W047°33'21.5" | *Phaseolus vulgaris* | *Spe*I | BR:Par1:12 | MT626919 |
| GO-65 | June, 2012 | Paranoá, DF | S16°00'47" | W047°33'21.5" | *Phaseolus vulgaris* | *Spe*I | BR:Par2.1:12 | MT626920 |
| | | | | | | *Spe*I | BR:Par2.2:12 | MT626921 |
| GO-66 | June, 2012 | Paranoá, DF | S16°00'47" | W047°33'21.5" | *Phaseolus vulgaris* | *Spe*I | BR:Par15.1:12 | MT626934 |
| | | | | | | *Hind*III | BR:Par15.2:12 | MT626935 |
| GO-67 | June, 2012 | Paranoá, DF | S16°00'47" | W047°33'21.5" | *Phaseolus vulgaris* | *Hind*III | BR:Par67:12 | MT626948 |
| GO-68 | June, 2012 | Paranoá, DF | S16°00'47" | W047°33'21.5" | *Phaseolus vulgaris* | *Spe*I | BR:Par16.1:12 | MT626936 |
| | | | | | | *Hind*III | BR:Par16.2:12 | MT626937 |
| GO-70 | June, 2012 | Paranoá, DF | S16°00'47" | W047°33'21.5" | *Phaseolus vulgaris* | *Spe*I | BR:Par3:12 | MT626922 |
| GO-71 | June, 2012 | Paranoá, DF | S16°00'47" | W047°33'21.5" | *Phaseolus vulgaris* | *Spe*I | BR:Par17:12 | MT626938 |
| GO-72 | June, 2012 | Paranoá, DF | S16°00'47" | W047°33'21.5" | *Phaseolus vulgaris* | *Spe*I | BR:Par4:12 | MT626923 |
| GO-73 | June, 2012 | Paranoá, DF | S16°00'47" | W047°33'21.5" | *Phaseolus vulgaris* | *Hind*III | BR:Par5.1:12 | MT626924 |
| | | | | | | *Spe*I | BR:Par5.2:12 | MT626925 |
| GO-75 | June, 2012 | Paranoá, DF | S15°57'57.4" | W047°31'35.8" | *Phaseolus vulgaris* | *Spe*I | BR:Par6.1:12 | MT626926 |
| | | | | | | *Hind*III | BR:Par6.2:12 | MT626927 |
| GO-76 | June, 2012 | Paranoá, DF | S15°57'57.4" | W047°31'35.8" | *Phaseolus vulgaris* | *Spe*I | BR:Par18:12 | MT626939 |
| GO-78 | June, 2012 | Paranoá, DF | S15°57'57.4" | W047°31'35.8" | *Phaseolus vulgaris* | *Spe*I | BR:Par19:12 | MT626940 |
| GO-79 | June, 2012 | Paranoá, DF | S15°57'57.4" | W047°31'35.8" | *Phaseolus vulgaris* | *Spe*I | BR:Par8:12 | MT626928 |
| GO-80 | June, 2012 | Paranoá, DF | S15°57'57.4" | W047°31'35.8" | *Phaseolus vulgaris* | *Spe*I | BR:Par20.1:12 | MT626941 |
| | | | | | | *Hind*III | BR:Par20.2:12 | MT626942 |
| G0-81 | June, 2012 | Paranoá, DF | S15°57'57.4" | W047°31'35.8" | *Phaseolus vulgaris* | *Spe*I | BR:Par21:12 | MT626943 |
| GO-82 | June, 2013 | Paranoá, DF | S15°57'57.4" | W047°31'35.8" | *Phaseolus vulgaris* | *Spe*I | BR:Par22:12 | MT626944 |
| GO-85 | June, 2012 | Paranoá, DF | S15°57'57.4" | W047°31'35.8" | *Phaseolus vulgaris* | *Spe*I | BR:Par23:12 | MT626945 |
| GO-88 | June, 2012 | Paranoá, DF | S15°57'57.4" | W047°31'35.8" | *Phaseolus vulgaris* | *Spe*I | BR:Par9.1:12 | MT626929 |
| | | | | | | *Spe*I | BR:Par9.2:12 | MT626930 |
| GO-89 | June, 2012 | Paranoá, DF | S15°57'57.4" | W047°31'35.8" | *Phaseolus vulgaris* | *Spe*I | BR:Par10:12 | MT626931 |
| GO-91 | June, 2012 | Paranoá, DF | S15°57'57.4" | W047°31'35.8" | *Phaseolus vulgaris* | *Spe*I | BR:Par24:12 | MT626946 |
| GO-92 | June, 2012 | Paranoá, DF | S15°57'57.4" | W047°31'35.8" | *Phaseolus vulgaris* | *Spe*I | BR:Par25:12 | MT626947 |
| GO-200 | June, 2012 | Cristalina, GO | S17°05'29.2" | W047°37'01.4" | *Phaseolus vulgaris* | *Spe*I | BR:Cri9:12 | MT626897 |
| GO-201 | June, 2012 | Cristalina, GO | S17°05'29.2" | W047°37'01.4" | *Phaseolus vulgaris* | *Spe*I | BR:Cri10:12 | MT626898 |
| GO-204 | June, 2012 | Cristalina, GO | S17°05'29.2" | W047°37'01.4" | *Phaseolus vulgaris* | *Spe*I | BR:Cri11:12 | MT626899 |
| GO-207 | June, 2012 | Cristalina, GO | S17°05'29.2" | W047°37'01.4" | *Phaseolus vulgaris* | *Spe*I | BR:Cri12:12 | MT626900 |
| GO-214 | June, 2012 | Cristalina, GO | S17°05'29.2" | W047°37'01.4" | *Phaseolus vulgaris* | *Spe*I | BR:Cri4:12 | MT626892 |
| GO-221 | June, 2012 | Cristalina, GO | S17°05'29.2" | W047°37'01.4" | *Phaseolus vulgaris* | *Spe*I | BR:Cri6:12 | MT626893 |

| | | | | | | | | |
|---|---|---|---|---|---|---|---|---|
| | | | | | | *Spe*I | BR:Cri6.1:12 | MT626894 |
| GO-222 | June, 2012 | Cristalina, GO | S17°05'29.2" | W047°37'01.4" | *Phaseolus vulgaris* | *Spe*I | BR:Cri7.1:12 | MT626895 |
| | | | | | | *Spe*I | BR:Cri7.2:12 | MT626896 |
| GO-227 | June, 2012 | Cristalina, GO | S17°05'29.2" | W047°37'01.4" | *Phaseolus vulgaris* | *Spe*I | BR:Cri13.1:12 | MT626901 |
| | | | | | | *Spe*I | BR:Cri13.2:12 | MT626902 |
| GO-228 | June, 2012 | Cristalina, GO | S17°05'29.2" | W047°37'01.4" | *Phaseolus vulgaris* | *Spe*I | BR:Cri14:12 | MT626903 |
| GO-236 | June, 2012 | Cristalina, GO | S17°05'29.2" | W047°37'01.4" | *Phaseolus vulgaris* | *Spe*I | BR:Cri15:12 | MT626904 |
| GO-238 | June, 2012 | Cristalina, GO | S17°05'29.2" | W047°37'01.4" | *Phaseolus vulgaris* | *Spe*I | BR:Cri16:12 | MT626905 |
| GO241 | June, 2012 | Cristalina, GO | S17°05'29.2" | W047°37'01.4" | *Phaseolus vulgaris* | *Spe*I | BR:Cri17:12 | MT626906 |
| 159AL | July, 2011 | Palmeira dos Índios, AL | S09°27'11.3" | W036°36'34.3" | *Phaseolus lunatus* | *Spe*I | BR:Pai2:11 | MT626949 |
| 162AL | July, 2011 | Palmeira dos Índios, AL | S09°27'11.3" | W036°36'34.3" | *Phaseolus lunatus* | *Spe*I | BR:Pai4:11 | MT626950 |
| 163AL | July, 2011 | Palmeira dos Índios, AL | S09°27'11.3" | W036°36'34.3" | *Phaseolus lunatus* | *Spe*I | BR:Pai5:11 | MT626951 |
| 164AL | July, 2011 | Palmeira dos Índios, AL | S09°27'11.3" | W036°36'34.3" | *Phaseolus lunatus* | *Spe*I | BR:Pai6:11 | MT626952 |
| 165AL | July, 2011 | Palmeira dos Índios, AL | S09°27'11.3" | W036°36'34.3" | *Phaseolus lunatus* | *Spe*I | BR:Pai7:11 | MT626953 |
| 166AL | July, 2011 | Palmeira dos Índios, AL | S09°27'11.3" | W036°36'34.3" | *Phaseolus lunatus* | *Spe*I | BR:Pai8:11 | MT626954 |
| 167AL | July, 2011 | Palmeira dos Índios, AL | S09°27'11.3" | W036°36'34.3" | *Phaseolus lunatus* | *Spe*I | BR:Pai9:11 | MT626955 |
| 168AL | July, 2011 | Palmeira dos Índios, AL | S09°27'11.3" | W036°36'34.3" | *Phaseolus lunatus* | *Spe*I | BR:Pai10:11 | MT626956 |
| 169AL | July, 2011 | Palmeira dos Índios, AL | S09°27'11.3" | W036°36'34.3" | *Phaseolus lunatus* | *Sac*I | BR:Pai11:11 | MT626957 |
| 170AL | July, 2011 | Palmeira dos Índios, AL | S09°27'11.3" | W036°36'34.3" | *Phaseolus lunatus* | *Spe*I | BR:Pai12:11 | MT626958 |
| 173AL | July, 2011 | Palmeira dos Índios, AL | S09°27'11.3" | W036°36'34.3" | *Phaseolus lunatus* | *Spe*I | BR:Pai13.1:11 | MT626959 |
| | | | | | | *Spe*I | BR:Pai13.2:11 | MT626960 |
| | | | | | | *Spe*I | BR:Pai13.3:11 | MT626961 |
| 174AL | July, 2011 | Palmeira dos Índios, AL | S09°27'11.3" | W036°36'34.3" | *Phaseolus lunatus* | *Spe*I | BR:Pai14.1:11 | MT626962 |
| | | | | | | *Spe*I | BR:Pai14.2:11 | MT626963 |
| 176AL | July, 2011 | Palmeira dos Índios, AL | S09°27'11.3" | W036°36'34.3" | *Phaseolus lunatus* | *Sac*I | BR:Pai15:11 | MT626964 |
| 177AL | July, 2011 | Palmeira dos Índios, AL | S09°27'11.3" | W036°36'34.3" | *Phaseolus lunatus* | *Sac*I | BR:Pai16.1:11 | MT626965 |
| | | | | | | *Spe*I | BR:Pai16.2:11 | MT626966 |
| 178AL | July, 2011 | Palmeira dos Índios, AL | S09°27'11.3" | W036°36'34.3" | *Phaseolus lunatus* | *Sac*I | BR:Pai17:11 | MT626967 |
| 254AL | July, 2011 | Murici, AL | S09°17'10.0" | W035°57'43.3" | *Phaseolus lunatus* | *Spe*I | BR:Mur1:11 | MT626968 |
| 255AL | July, 2011 | Murici, AL | S09°17'10.0" | W035°57'43.3" | *Phaseolus lunatus* | *Spe*I | BR:Mur6.1:11 | MT626975 |
| | | | | | | *Spe*I | BR:Mur6.2:11 | MT626976 |
| 257AL | July, 2011 | Murici, AL | S09°17'10.0" | W035°57'43.3" | *Phaseolus lunatus* | *Sac*I | BR:Mur7:11 | MT626977 |
| 258AL | July, 2011 | Murici, AL | S09°17'10.0" | W035°57'43.3" | *Phaseolus lunatus* | *Spe*I | BR:Mur8:11 | MT626978 |
| 262AL | July, 2011 | Murici, AL | S09°17'10.0" | W035°57'43.3" | *Phaseolus lunatus* | *Spe*I | BR:Mur10.1:11 | MT626979 |
| | | | | | | *Spe*I | BR:Mur10.2:11 | MT626980 |
| 265AL | July, 2011 | Murici, AL | S09°17'10.0" | W035°57'43.3" | *Phaseolus lunatus* | *Sac*I | BR:Mur2.1:11 | MT626969 |

| | | | | | | | | |
|---|---|---|---|---|---|---|---|---|
| | | | | | | *Spe*I | BR:Mur2.2:11 | MT626970 |
| 269AL | July, 2011 | Murici, AL | S09°17'10.0" | W035°57'43.3" | *Phaseolus lunatus* | *Sac*I | BR:Mur3:11 | MT626971 |
| 272AL | July, 2011 | Murici, AL | S09°17'10.0" | W035°57'43.3" | *Phaseolus lunatus* | *Spe*I | BR:Mur12.1:11 | MT626982 |
| | | | | | | *Spe*I | BR:Mur12.2:11 | MT626981 |
| 274AL | July, 2011 | Murici, AL | S09°17'10.0" | W035°57'43.3" | *Phaseolus lunatus* | *Spe*I | BR:Mur4:11 | MT626972 |
| 275AL | July, 2011 | Murici, AL | S09°17'10.0" | W035°57'43.3" | *Phaseolus lunatus* | *Spe*I | BR:Mur5.1:11 | MT626973 |
| | | | | | | *Spe*I | BR:Mur5.2:11 | MT626974 |
| 310MG | March, 2011 | Florestal, MG | S19°52'58.4" | W44°25'14.6" | *Macroptilium lathyroides* | *Xba*I | BR:Flt1:11 | MT626879 |
| 311MG | March, 2011 | Florestal, MG | S19°52'58.4" | W44°25'14.6" | *Macroptilium lathyroides* | *Xba*I | BR:Flt2:11 | MT626880 |
| 321MG | March, 2011 | Florestal, MG | S19°52'58.4" | W44°25'14.6" | *Macroptilium lathyroides* | *Hind*III | BR:Flt20:11 | MT626890 |
| 323MG | | Florestal, MG | S19°52'58.4" | W44°25'14.6" | *Macroptilium lathyroides* | *Xba*I | BR:Flt323:11 | MT626891 |
| 325MG | March, 2011 | Florestal, MG | S19°52'58.4" | W44°25'14.6" | *Macroptilium lathyroides* | *Hind*III | BR:Flt5.1:11 | MT626881 |
| | | | | | | *Xba*I | BR:Flt5.2:11 | MT626882 |
| 326MG | March, 2011 | Florestal, MG | S19°52'58.4" | W44°25'14.6" | *Macroptilium lathyroides* | *Hind*III | BR:Flt6:11 | MT626883 |
| | | | | | | *Xba*I | BR:Flt6.1:11 | MT626884 |
| 330MG | March, 2011 | Florestal, MG | S19°52'58.4" | W44°25'14.6" | *Macroptilium lathyroides* | *Xba*I | BR:Flt8.1:11 | MT626885 |
| | | | | | | *Xba*I | BR:Flt8.2:11 | MT626886 |
| 332MG | March, 2011 | Florestal, MG | S19°52'58.4" | W44°25'14.6" | *Macroptilium lathyroides* | *Xba*I | BR:Flt13:11 | MT626889 |
| 337MG | March, 2011 | Florestal, MG | S19°52'58.4" | W44°25'14.6" | *Macroptilium lathyroides* | *Xba*I | BR:Flt11:11 | MT626887 |
| 338MG | March, 2011 | Florestal, MG | S19°52'58.4" | W44°25'14.6" | *Macroptilium lathyroides* | *Xba*I | BR:Flt12:11 | MT626888 |
| *Blainvillea yellow spot virus* (BlYSV) | | | | | | | | |
| HV4 | May, 2010 | Viçosa, MG* | S20°45'22.9" | W42°50'58.0" | *Blainvillea rhomboidea* | *Sac*I | BR:Vic04.1:10 | MT626997 |
| HV9 | May, 2010 | Viçosa, MG | S20°45'22.9" | W42°50'58.0" | *Blainvillea rhomboidea* | *Sac*I | BR:Vic09:10 | MT626998 |
| HV13 | May, 2010 | Viçosa, MG | S20°45'22.9" | W42°50'58.0" | *Blainvillea rhomboidea* | *Sac*I | BR:Vic13:10 | MT626999 |
| OS32 | July, 2013 | Coimbra, MG | S20°51'29.9" | W042°51'42.2" | *Blainvillea rhomboidea* | *Cla*I | BR:Coi32.1:13 | MT627000 |
| OS33 | July, 2013 | Coimbra, MG | S20°51'29.9" | W042°51'42.2" | *Blainvillea rhomboidea* | *Sac*I | BR:Coi33:13 | MT627001 |
| OS34 | July, 2013 | Coimbra, MG | S20°51'29.9" | W042°51'42.2" | *Blainvillea rhomboidea* | *Sac*I | BR:Coi34:13 | MT627002 |
| OS35 | July, 2013 | Coimbra, MG | S20°51'29.9" | W042°51'42.2" | *Blainvillea rhomboidea* | *Sac*I | BR:Coi35:13 | MT627003 |
| OS36 | July, 2013 | Coimbra, MG | S20°51'29.9" | W042°51'42.2" | *Blainvillea rhomboidea* | *Sac*I | BR:Coi36:13 | MT627004 |
| OS37 | July, 2013 | Coimbra, MG | S20°51'29.9" | W042°51'42.2" | *Blainvillea rhomboidea* | *Xba*I | BR:Coi37.1:13 | MT627005 |
| OS51 | July, 2013 | Coimbra, MG | S20°51'29.9" | W042°51'42.2" | *Blainvillea rhomboidea* | *Sac*I | BR:Coi51:13 | MT627006 |
| OS142 | February, 2014 | Coimbra, MG | S20°50'58.3" | W042°53'14.4" | *Blainvillea rhomboidea* | *Sac*I | BR:Coi142.1:14 | MT627007 |
| | | | | | | *Sac*I | BR:Coi142.2:14 | MT627008 |
| OS145 | February, 2014 | Coimbra, MG | S20°50'58.3" | W042°53'14.4" | *Blainvillea rhomboidea* | *Sac*I | BR:Coi145:14 | MT627009 |
| OS147 | February, 2014 | Coimbra, MG | S20°50'58.3" | W042°53'14.4" | *Blainvillea rhomboidea* | *Sac*I | BR:Coi147:14 | MT627010 |
| OS148 | February, 2014 | Coimbra, MG | S20°50'58.3" | W042°53'14.4" | *Blainvillea rhomboidea* | *Sac*I | BR:Coi148:14 | MT627011 |

| | | | | | | | | |
|---|---|---|---|---|---|---|---|---|
| OS150 | February, 2014 | Coimbra, MG | S20°50'58.3" | W042°53'14.4" | *Blainvillea rhomboidea* | *Sac*I | BR:Coi150:14 | MT627012 |
| OS151 | February, 2014 | Coimbra, MG | S20°50'58.3" | W042°53'14.4" | *Blainvillea rhomboidea* | *Sac*I | BR:Coi151:14 | MT627013 |
| OS152 | February, 2014 | Coimbra, MG | S20°50'58.3" | W042°53'14.4" | *Blainvillea rhomboidea* | *Sac*I | BR:Coi152.1:14 | MT627014 |
| | | | | | | *Sac*I | BR:Coi152.2:14 | MT627015 |
| OS154 | February, 2014 | Coimbra, MG | S20°50'58.3" | W042°53'14.4" | *Blainvillea rhomboidea* | *Sac*I | BR:Coi154:14 | MT627016 |
| OS155 | February, 2014 | Coimbra, MG | S20°50'58.3" | W042°53'14.4" | *Blainvillea rhomboidea* | *Sac*I | BR:Coi155:14 | MT627017 |
| OS157 | February, 2014 | Coimbra, MG | S20°50'58.3" | W042°53'14.4" | *Blainvillea rhomboidea* | *Sac*I | BR:Coi157:14 | MT627018 |
| OS158 | February, 2014 | Coimbra, MG | S20°50'58.3" | W042°53'14.4" | *Blainvillea rhomboidea* | *Sac*I | BR:Coi158s:14 | MT627019 |
| | | | | | | *Bam*HI | BR:Coi158b:14 | MT627020 |
| OS159 | February, 2014 | Coimbra, MG | S20°50'58.3" | W042°53'14.4" | *Blainvillea rhomboidea* | *Sac*I | BR:Coi159:14 | MT627021 |
| OS160 | February, 2014 | Coimbra, MG | S20°50'58.3" | W042°53'14.4" | *Blainvillea rhomboidea* | *Sac*I | BR:Coi160.1:14 | MT627022 |
| | | | | | | *Sac*I | BR:Coi160.2:14 | MT627023 |
| OS164 | February, 2014 | Coimbra, MG | S20°50'58.3" | W042°53'14.4" | *Blainvillea rhomboidea* | *Sac*I | BR:Coi164:14 | MT627024 |
| OS165 | February, 2014 | Coimbra, MG | S20°50'58.3" | W042°53'14.4" | *Blainvillea rhomboidea* | *Sac*I | BR:Coi165:14 | MT627025 |
| *Macroptilium yellow spot virus* (MaYSV) | | | | | | | | |
| RC18 | July, 2011 | Craíbas, AL | S09°40'37.9" | W036°46'37.8" | *Phaseolus vulgaris* | *Pst*I | BR:Crb1:11 | MT627026 |
| RC19 | July, 2011 | Craíbas, AL | S09°40'39.1" | W036°46'38.4" | *Phaseolus vulgaris* | *Pst*I | BR:Crb2:11 | MT627027 |
| RC29 | July, 2011 | Craíbas, AL | S09°40'39.8" | W036°46'38.4" | *Phaseolus vulgaris* | *Pst*I | BR:Crb10:11 | MT627028 |
| RC77 | July, 2011 | Olho D'água das Flores, AL | S09°32'28.1" | W037°17'26.8" | *Phaseolus vulgaris* | *Pst*I | BR:Oaf8:11 | MT627029 |
| 46AL | July, 2011 | Olho D'água das Flores, AL | S09°32'27.8" | W037°17'22.1" | *Macroptilium lathyroides* | *Pst*I | BR:Oaf28:11 | MT627030 |
| 61AL | July, 2011 | Olho D'água das Flores, AL | S09°32'26.9" | W037°17'17.6" | *Macroptilium lathyroides* | *Pst*I | BR:Oaf27:11 | MT627031 |
| RC94 | July, 2011 | Olho D'água das Flores, AL | S09°32'59.4" | W037°18'43.5" | *Macroptilium lathyroides* | *Pst*I | BR:Oaf24:11 | MT627032 |
| RC95 | July, 2011 | Olho D'água das Flores, AL | S09°32'59.2" | W037°18'43.0" | *Macroptilium lathyroides* | *Pst*I | BR:Oaf25.1:11 | MT627033 |
| | | | | | | *Pst*I | BR:Oaf25.2:11 | MT627034 |
| 97AL | July, 2011 | Santana do Ipanema, AL | S09°23'25.1" | W037°12'46.9" | *Phaseolus lunatus* | *Pst*I | BR:Sti34:11 | MT627043 |
| 99AL | July, 2011 | Santana do Ipanema, AL | S09°23'24.7" | W037°12'48.0" | *Phaseolus lunatus* | *Bgl*II | BR:Sti2:11 | MT627035 |
| 102AL | July, 2011 | Santana do Ipanema, AL | S09°23'24.9" | W037°12'48.0" | *Phaseolus lunatus* | *Bgl*II | BR:Sti4:11 | MT627036 |
| 105AL | July, 2011 | Santana do Ipanema, AL | S09°23'24.9" | W037°12'48.3" | *Phaseolus lunatus* | *Pst*I | BR:Sti35:11 | MT627044 |
| 111AL | July, 2011 | Santana do Ipanema, AL | S09°23'24.9" | W037°12'47.5" | *Phaseolus lunatus* | *Bgl*II | BR:Sti9.1:11 | MT627037 |
| | | | | | | *Bgl*II | BR:Sti9.2:11 | MT627038 |
| | | | | | | *Spe*I | BR:Sti9.3:11 | MT627040 |
| 112AL | July, 2011 | Santana do Ipanema, AL | S09°23'24.9" | W037°12'47.6" | *Phaseolus lunatus* | *Pst*I | BR:Sti10:11 | MT627039 |
| 128AL | July, 2011 | Santana do Ipanema, AL | S09°23'24.9" | W037°12'47.6" | *Phaseolus lunatus* | *Apa*I | BR:Sti26:11 | MT627041 |
| 152AL | July, 2011 | Santana do Ipanema, AL | S09°23'24.8" | W037°12'48.6" | *Phaseolus lunatus* | *Xba*I | BR:Sti29:11 | MT627042 |
| *Tomato severe rugose virus* (ToSRV) | | | | | | | | |
| DV202 | July, 2008 | Florestal, MG | S19°52'25.4" | W44°25'00.6" | *Solanum lycopersicum* | *Kpn*I | BR:Flo202.3:08 | MT627160 |

| | | | | | | | | |
|---|---|---|---|---|---|---|---|---|
| | | | | | | *Kpn*I | BR:Flo202.4:08 | MT627161 |
| DV203 | July, 2008 | Florestal, MG | S19°52'25.4" | W44°25'00.6" | *Solanum lycopersicum* | *Kpn*I | BR:Flo203:08 | MT627162 |
| DV206 | July, 2008 | Florestal, MG | S19°52'25.4" | W44°25'00.6" | *Solanum lycopersicum* | *Kpn*I | BR:Flo206:08 | MT627163 |
| DV208 | July, 2008 | Florestal, MG | S19°52'25.4" | W44°25'00.6" | *Solanum lycopersicum* | *Kpn*I | BR:Flo208:08 | MT627164 |
| DV218 | July, 2008 | Carandaí, MG | S20°56'56.5" | W43°47'42.2" | *Solanum lycopersicum* | *Kpn*I | BR:Car218:08 | MT627098 |
| DV219 | July, 2008 | Carandaí, MG | S20°56'56.5" | W43°47'42.2" | *Solanum lycopersicum* | *Kpn*I | BR:Car219:08 | MT627099 |
| DV220 | July, 2008 | Carandaí, MG | S20°56'56.5" | W43°47'42.2" | *Solanum lycopersicum* | *Kpn*I | BR:Car220:08 | MT627100 |
| DV224 | July, 2008 | Carandaí, MG | S20°56'56.5" | W43°47'42.2" | *Solanum lycopersicum* | *Kpn*I | BR:Car224.1:08 | MT627101 |
| | | | | | | *Kpn*I | BR:Car224.3:08 | MT627102 |
| DV226 | July, 2008 | Carandaí, MG | S20°56'56.5" | W43°47'42.2" | *Solanum lycopersicum* | *Kpn*I | BR:Car226.1:08 | MT627103 |
| | | | | | | *Kpn*I | BR:Car226.3:08 | MT627104 |
| DV230 | July, 2008 | Carandaí, MG | S20°56'56.5" | W43°47'42.2" | *Solanum lycopersicum* | *Kpn*I | BR:Car230:08 | MT627105 |
| DV232 | July, 2008 | Carandaí, MG | S20°56'56.5" | W43°47'42.2" | *Solanum lycopersicum* | *Kpn*I | BR:Car232:08 | MT627106 |
| DV237 | July, 2008 | Carandaí, MG | S20°56'56.5" | W43°47'42.2" | *Solanum lycopersicum* | *Kpn*I | BR:Car237.1:08 | MT627107 |
| | | | | | | *Kpn*I | BR:Car237.2:08 | MT627108 |
| DV238 | July, 2008 | Carandaí, MG | S20°56'56.5" | W43°47'42.2" | *Solanum lycopersicum* | *Kpn*I | BR:Car238.1:08 | MT627109 |
| OS70 | July, 2013 | Coimbra, MG | S20°51'43.2" | W042°51'27" | *Solanum lycopersicum* | *Kpn*I | BR:Coi70:13 | MT627110 |
| OS73 | July, 2013 | Coimbra, MG | S20°51'43.2" | W042°51'27" | *Solanum lycopersicum* | *Kpn*I | BR:Coi73:13 | MT627111 |
| OS79 | July, 2013 | Coimbra, MG | S20°51'43.2" | W042°51'27" | *Solanum lycopersicum* | *Kpn*I | BR:Coi79:13 | MT627112 |
| OS80 | July, 2013 | Coimbra, MG | S20°51'43.2" | W042°51'27" | *Solanum lycopersicum* | *Kpn*I | BR:Coi80:13 | MT627113 |
| OS83 | July, 2013 | Coimbra, MG | S20°51'43.2" | W042°51'27" | *Solanum lycopersicum* | *Kpn*I | BR:Coi83:13 | MT627114 |
| OS84 | July, 2013 | Coimbra, MG | S20°51'43.2" | W042°51'27" | *Solanum lycopersicum* | *Kpn*I | BR:Coi84:13 | MT627115 |
| OS85 | July, 2013 | Coimbra, MG | S20°51'43.2" | W042°51'27" | *Solanum lycopersicum* | *Kpn*I | BR:Coi85:13 | MT627116 |
| OS86 | July, 2013 | Coimbra, MG | S20°51'43.2" | W042°51'27" | *Solanum lycopersicum* | *Kpn*I | BR:Coi86:13 | MT627117 |
| OS92 | July, 2013 | Coimbra, MG | S20°51'43.2" | W042°51'27" | *Solanum lycopersicum* | *Kpn*I | BR:Coi92.2:13 | MT627118 |
| | | | | | | *Kpn*I | BR:Coi92.5:13 | MT627119 |
| OS93 | July, 2013 | Coimbra, MG | S20°51'43.2" | W042°51'27" | *Solanum lycopersicum* | *Kpn*I | BR:Coi93:13 | MT627120 |
| OS95 | July, 2013 | Coimbra, MG | S20°51'43.2" | W042°51'27" | *Solanum lycopersicum* | *Kpn*I | BR:Coi95:13 | MT627121 |
| OS103 | July, 2013 | Coimbra, MG | S20°51'43.2" | W042°51'27" | *Solanum lycopersicum* | *Kpn*I | BR:Coi103.1:13 | MT627122 |
| | | | | | | *Kpn*I | BR:Coi103.4:13 | MT627123 |
| OS110 | July, 2013 | Coimbra, MG | S20°51'43.2" | W042°51'27" | *Solanum lycopersicum* | *Kpn*I | BR:Coi110:13 | MT627124 |
| OS114 | July, 2013 | Coimbra, MG | S20°51'43.2" | W042°51'27" | *Solanum lycopersicum* | *Kpn*I | BR:Coi114:13 | MT627125 |
| OS117 | July, 2013 | Coimbra, MG | S20°51'43.2" | W042°51'27" | *Solanum lycopersicum* | *Kpn*I | BR:Coi117:13 | MT627126 |
| | | | | | | *Kpn*I | BR:Coi117.1:13 | MT627127 |
| OS119 | July, 2013 | Coimbra, MG | S20°36'39.3 " | W042°25'58.9" | *Solanum lycopersicum* | *Kpn*I | BR:Coi119.1:13 | MT627128 |
| | | | | | | *Kpn*I | BR:Coi119.4:13 | MT627129 |

| | | | | | | | | |
|---|---|---|---|---|---|---|---|---|
| OS120 | February, 2014 | Coimbra, MG | S20°36'39.3" | W042°25'58.9" | *Solanum lycopersicum* | *Kpn*I | BR:Coi120:14 | MT627130 |
| OS122 | February, 2014 | Coimbra, MG | S20°36'39.3" | W042°49'16.9" | *Solanum lycopersicum* | *Kpn*I | BR:Coi122:14 | MT627131 |
| OS125 | February, 2014 | Coimbra, MG | S20°50'51 " | W042°52'41.2" | *Solanum lycopersicum* | *Kpn*I | BR:Coi125:14 | MT627132 |
| OS131 | February, 2014 | Coimbra, MG | S20°50'49.4" | W042°52'41.9" | *Solanum lycopersicum* | *Kpn*I | BR:Coi131:14 | MT627133 |
| OS172 | February, 2014 | Coimbra, MG | S20°50'58.3" | W042°53'14.4" | *Solanum lycopersicum* | *Kpn*I | BR:Coi172:14 | MT627134 |
| OS179 | February, 2014 | Coimbra, MG | S20°50'58.3" | W042°53'14.4" | *Solanum lycopersicum* | *Kpn*I | BR:Coi179:14 | MT627135 |
| OS182 | February, 2014 | Coimbra, MG | S20°50'58.3" | W042°53'14.4" | *Solanum lycopersicum* | *Kpn*I | BR:Coi182:14 | MT627136 |
| OS183 | February, 2014 | Coimbra, MG | S20°50'58.3" | W042°53'14.4" | *Solanum lycopersicum* | *Kpn*I | BR:Coi183:14 | MT627137 |
| OS201 | June, 2014 | Florestal, MG | S19°55'55.8" | W044°23'52.4" | *Solanum lycopersicum* | *Kpn*I | BR:Flo01:14 | MT627138 |
| OS202 | June, 2014 | Florestal, MG | S19°55'55.8" | W044°23'52.4" | *Solanum lycopersicum* | *Kpn*I | BR:Flo02:14 | MT627139 |
| OS203 | June, 2014 | Florestal, MG | S19°55'55.8" | W044°23'52.4" | *Solanum lycopersicum* | *Kpn*I | BR:Flo03:14 | MT627140 |
| OS204 | June, 2014 | Florestal, MG | S19°55'55.8" | W044°23'52.4" | *Solanum lycopersicum* | *Kpn*I | BR:Flo04.1:14 | MT627141 |
| | | | | | | *Kpn*I | BR:Flo04.5:14 | MT627142 |
| OS205 | June, 2014 | Florestal, MG | S19°55'55.8" | W044°23'52.4" | *Solanum lycopersicum* | *Kpn*I | BR:Flo05:14 | MT627143 |
| OS206 | June, 2014 | Florestal, MG | S19°55'55.8" | W044°23'52.4" | *Solanum lycopersicum* | *Kpn*I | BR:Flo06:14 | MT627144 |
| OS207 | June, 2014 | Florestal, MG | S19°55'55.8" | W044°23'52.4" | *Solanum lycopersicum* | *Kpn*I | BR:Flo07:14 | MT627145 |
| OS208 | June, 2014 | Florestal, MG | S19°55'55.8" | W044°23'52.4" | *Solanum lycopersicum* | *Kpn*I | BR:Flo08.1:14 | MT627146 |
| | | | | | | *Kpn*I | BR:Flo08.2:14 | MT627147 |
| OS209 | June, 2014 | Florestal, MG | S19°55'55.8" | W044°23'52.4" | *Solanum lycopersicum* | *Kpn*I | BR:Flo09.1:14 | MT627148 |
| | | | | | | *Kpn*I | BR:Flo09.3:14 | MT627149 |
| OS210 | June, 2014 | Florestal, MG | S19°55'55.8" | W044°23'52.4" | *Solanum lycopersicum* | *Kpn*I | BR:Flo10:14 | MT627150 |
| OS211 | June, 2014 | Florestal, MG | S19°55'55.8" | W044°23'52.4" | *Solanum lycopersicum* | *Kpn*I | BR:Flo11:14 | MT627151 |
| OS212 | June, 2014 | Florestal, MG | S19°55'55.8" | W044°23'52.4" | *Solanum lycopersicum* | *Kpn*I | BR:Flo12:14 | MT627152 |
| OS213 | June, 2014 | Florestal, MG | S19°55'55.8" | W044°23'52.4" | *Solanum lycopersicum* | *Kpn*I | BR:Flo13:14 | MT627153 |
| OS214 | June, 2014 | Florestal, MG | S19°55'55.8" | W044°23'52.4" | *Solanum lycopersicum* | *Kpn*I | BR:Flo14:14 | MT627154 |
| OS216 | June, 2014 | Florestal, MG | S19°55'55.8" | W044°23'52.4" | *Solanum lycopersicum* | *Kpn*I | BR:Flo16:14 | MT627155 |
| OS218 | June, 2014 | Florestal, MG | S19°55'55.8" | W044°23'52.4" | *Solanum lycopersicum* | *Kpn*I | BR:Flo18:14 | MT627156 |
| OS219 | June, 2014 | Florestal, MG | S19°55'55.8" | W044°23'52.4" | *Solanum lycopersicum* | *Kpn*I | BR:Flo19:14 | MT627157 |
| OS220 | June, 2014 | Florestal, MG | S19°55'55.8" | W044°23'52.4" | *Solanum lycopersicum* | *Kpn*I | BR:Flo20:14 | MT627158 |
| OS222 | June, 2014 | Florestal, MG | S19°55'55.8" | W044°23'52.4" | *Solanum lycopersicum* | *Kpn*I | BR:Flo222:14 | MT627159 |

*State sigla: AL, Alagoas; BA, Bahia; DF, Federal District; GO, Goiás; MG, Minas Gerais; MS, Mato Grosso do Sul; PE, Pernambuco; PR, Paraná; RS, Rio Grande do Sul

**Supplementary Table S2.** Begomovirus sequences retrieved from GenBank.

| Isolate name | GenBank access number | | Location | Geographicalcoordinates | | Date | Host | Reference |
|---|---|---|---|---|---|---|---|---|
| | DNA-A | DNA-B | | Latitude | Longitude | | | |
| *Bean golden mosaic virus* (BGMV) | | | | | | | | |
| BR:Pai1:11 | KJ939737 | | Palmeira dos Índios, AL | S09°27'11.3" | W036°36'34.3" | July, 2011 | *Phaseolus lunatus* | Sobrinho et al., 2014 |
| BR:Pai2:11 | KJ939738 | | Palmeira dos Índios, AL | S09°27'11.3" | W036°36'34.3" | July, 2011 | *Phaseolus lunatus* | Sobrinho et al., 2014 |
| BR:Pai3:11 | KJ939739 | | Palmeira dos Índios, AL | S09°27'11.3" | W036°36'34.3" | July, 2011 | *Phaseolus lunatus* | Sobrinho et al., 2014 |
| BR:Pai4:11 | KJ939740 | | Palmeira dos Índios, AL | S09°27'11.3" | W036°36'34.3" | July, 2011 | *Phaseolus lunatus* | Sobrinho et al., 2014 |
| BR:Pai5:11 | KJ939741 | | Palmeira dos Índios, AL | S09°27'11.3" | W036°36'34.3" | July, 2011 | *Phaseolus lunatus* | Sobrinho et al., 2014 |
| BR:Pai6:11 | KJ939742 | | Palmeira dos Índios, AL | S09°27'11.3" | W036°36'34.3" | July, 2011 | *Phaseolus lunatus* | Sobrinho et al., 2014 |
| BR:Pai7:11 | KJ939743 | | Palmeira dos Índios, AL | S09°27'11.3" | W036°36'34.3" | July, 2011 | *Phaseolus lunatus* | Sobrinho et al., 2014 |
| BR:Pai8:11 | KJ939744 | | Palmeira dos Índios, AL | S09°27'11.3" | W036°36'34.3" | July, 2011 | *Phaseolus lunatus* | Sobrinho et al., 2014 |
| BR:Pai9:11 | KJ939745 | | Palmeira dos Índios, AL | S09°27'11.3" | W036°36'34.3" | July, 2011 | *Phaseolus lunatus* | Sobrinho et al., 2014 |
| BR:Pai10:11 | KJ939746 | | Palmeira dos Índios, AL | S09°27'11.3" | W036°36'34.3" | July, 2011 | *Phaseolus lunatus* | Sobrinho et al., 2014 |
| BR:Pai11:11 | KJ939747 | | Palmeira dos Índios, AL | S09°27'11.3" | W036°36'34.3" | July, 2011 | *Phaseolus lunatus* | Sobrinho et al., 2014 |
| BR:Pai12:11 | KJ939748 | | Palmeira dos Índios, AL | S09°27'11.3" | W036°36'34.3" | July, 2011 | *Phaseolus lunatus* | Sobrinho et al., 2014 |
| BR:Pai13:11 | KJ939749 | | Palmeira dos Índios, AL | S09°27'11.3" | W036°36'34.3" | July, 2011 | *Phaseolus lunatus* | Sobrinho et al., 2014 |
| BR:Pai14:11 | KJ939750 | | Palmeira dos Índios, AL | S09°27'11.3" | W036°36'34.3" | July, 2011 | *Phaseolus lunatus* | Sobrinho et al., 2014 |
| BR:Pai15:11 | KJ939751 | | Palmeira dos Índios, AL | S09°27'11.3" | W036°36'34.3" | July, 2011 | *Phaseolus lunatus* | Sobrinho et al., 2014 |
| BR:Pai16:11 | KJ939752 | | Palmeira dos Índios, AL | S09°27'11.3" | W036°36'34.3" | July, 2011 | *Phaseolus lunatus* | Sobrinho et al., 2014 |
| BR:Pai17:11 | KJ939753 | | Palmeira dos Índios, AL | S09°27'11.3" | W036°36'34.3" | July, 2011 | *Phaseolus lunatus* | Sobrinho et al., 2014 |
| BR:Mur1:11 | KJ939754 | | Murici, AL | S09°17'10.0" | W035°57'43.3" | July, 2011 | *Phaseolus lunatus* | Sobrinho et al., 2014 |
| BR:Mur2:11 | KJ939755 | | Murici, AL | S09°17'10.0" | W035°57'43.3" | July, 2011 | *Phaseolus lunatus* | Sobrinho et al., 2014 |
| BR:Mur3:11 | KJ939756 | | Murici, AL | S09°17'10.0" | W035°57'43.3" | July, 2011 | *Phaseolus lunatus* | Sobrinho et al., 2014 |
| BR:Mur4:11 | KJ939757 | | Murici, AL | S09°17'10.0" | W035°57'43.3" | July, 2011 | *Phaseolus lunatus* | Sobrinho et al., 2014 |
| BR:Mur5:11 | KJ939758 | | Murici, AL | S09°17'10.0" | W035°57'43.3" | July, 2011 | *Phaseolus lunatus* | Sobrinho et al., 2014 |
| BR:Mur6:11 | KJ939759 | | Murici, AL | S09°17'10.0" | W035°57'43.3" | July, 2011 | *Phaseolus lunatus* | Sobrinho et al., 2014 |
| BR:Mur7:11 | KJ939760 | | Murici, AL | S09°17'10.0" | W035°57'43.3" | July, 2011 | *Phaseolus lunatus* | Sobrinho et al., 2014 |
| BR:Mur8:11 | KJ939761 | | Murici, AL | S09°17'10.0" | W035°57'43.3" | July, 2011 | *Phaseolus lunatus* | Sobrinho et al., 2014 |
| BR:Mur9:11 | KJ939762 | | Murici, AL | S09°17'10.0" | W035°57'43.3" | July, 2011 | *Phaseolus lunatus* | Sobrinho et al., 2014 |
| BR:Mur10:11 | KJ939763 | | Murici, AL | S09°17'10.0" | W035°57'43.3" | July, 2011 | *Phaseolus lunatus* | Sobrinho et al., 2014 |
| BR:Mur11:11 | KJ939764 | | Murici, AL | S09°17'10.0" | W035°57'43.3" | July, 2011 | *Phaseolus lunatus* | Sobrinho et al., 2014 |
| BR:Mur12:11 | KJ939765 | | Murici, AL | S09°17'10.0" | W035°57'43.3" | July, 2011 | *Phaseolus lunatus* | Sobrinho et al., 2014 |
| BR:Flt1:11 | KJ939766 | | Florestal, MG | S19°52'58.4" | W44°25'14.6" | March, 2011 | *Macroptilium lathyroides* | Sobrinho et al., 2014 |
| BR:Flt2:11 | KJ939767 | | Florestal, MG | S19°52'58.4" | W44°25'14.6" | March, 2011 | *Macroptilium lathyroides* | Sobrinho et al., 2014 |
| BR:Flt3:11 | KJ939768 | | Florestal, MG | S19°52'58.4" | W44°25'14.6" | March, 2011 | *Macroptilium lathyroides* | Sobrinho et al., 2014 |
| BR:Flt4:11 | KJ939769 | | Florestal, MG | S19°52'58.4" | W44°25'14.6" | March, 2011 | *Macroptilium lathyroides* | Sobrinho et al., 2014 |
| BR:Flt5:11 | KJ939770 | | Florestal, MG | S19°52'58.4" | W44°25'14.6" | March, 2011 | *Macroptilium lathyroides* | Sobrinho et al., 2014 |
| BR:Flt6:11 | KJ939771 | | Florestal, MG | S19°52'58.4" | W44°25'14.6" | March, 2011 | *Macroptilium lathyroides* | Sobrinho et al., 2014 |
| BR:Flt7:11 | KJ939772 | | Florestal, MG | S19°52'58.4" | W44°25'14.6" | March, 2011 | *Macroptilium lathyroides* | Sobrinho et al., 2014 |

| | | | | | | | |
|---|---|---|---|---|---|---|---|
| BR:Flt8:11 | KJ939773 | Florestal, MG | S19°52'58.4" | W44°25'14.6" | March, 2011 | *Macroptilium lathyroides* | Sobrinho et al., 2014 |
| BR:Flt9:11 | KJ939774 | Florestal, MG | S19°52'58.4" | W44°25'14.6" | March, 2011 | *Macroptilium lathyroides* | Sobrinho et al., 2014 |
| BR:Flt10:11 | KJ939775 | Florestal, MG | S19°52'58.4" | W44°25'14.6" | March, 2011 | *Macroptilium lathyroides* | Sobrinho et al., 2014 |
| BR:Flt11:11 | KJ939776 | Florestal, MG | S19°52'58.4" | W44°25'14.6" | March, 2011 | *Macroptilium lathyroides* | Sobrinho et al., 2014 |
| BR:Flt12:11 | KJ939777 | Florestal, MG | S19°52'58.4" | W44°25'14.6" | March, 2011 | *Macroptilium lathyroides* | Sobrinho et al., 2014 |
| BR:Flt13:11 | KJ939778 | Florestal, MG | S19°52'58.4" | W44°25'14.6" | March, 2011 | *Macroptilium lathyroides* | Sobrinho et al., 2014 |
| BR:Sag1:12 | KJ939779 | Santo Antônio de Góias, GO | S16°30'22.5" | W049°17'06.8" | June, 2012 | *Phaseolus vulgaris* | Sobrinho et al., 2014 |
| BR:Sag2:12 | KJ939780 | Santo Antônio de Góias, GO | S16°30'22.5" | W049°17'06.8" | June, 2012 | *Phaseolus vulgaris* | Sobrinho et al., 2014 |
| BR:Sag3:12 | KJ939781 | Santo Antônio de Góias, GO | S16°30'22.5" | W049°17'06.8" | June, 2012 | *Phaseolus vulgaris* | Sobrinho et al., 2014 |
| BR:Sag4:12 | KJ939782 | Santo Antônio de Góias, GO | S16°30'22.5" | W049°17'06.8" | June, 2012 | *Phaseolus vulgaris* | Sobrinho et al., 2014 |
| BR:Sag5:12 | KJ939783 | Santo Antônio de Góias, GO | S16°30'22.5" | W049°17'06.8" | June, 2012 | *Phaseolus vulgaris* | Sobrinho et al., 2014 |
| BR:Sag6:12 | KJ939784 | Santo Antônio de Góias, GO | S16°30'22.5" | W049°17'06.8" | June, 2012 | *Phaseolus vulgaris* | Sobrinho et al., 2014 |
| BR:Sag7:12 | KJ939785 | Santo Antônio de Góias, GO | S16°30'22.5" | W049°17'06.8" | June, 2012 | *Phaseolus vulgaris* | Sobrinho et al., 2014 |
| BR:Sag8:12 | KJ939786 | Santo Antônio de Góias, GO | S16°30'22.5" | W049°17'06.8" | June, 2012 | *Phaseolus vulgaris* | Sobrinho et al., 2014 |
| BR:Sag9:12 | KJ939787 | Santo Antônio de Góias, GO | S16°30'22.5" | W049°17'06.8" | June, 2012 | *Phaseolus vulgaris* | Sobrinho et al., 2014 |
| BR:Sag10:12 | KJ939788 | Santo Antônio de Góias, GO | S16°30'22.5" | W049°17'06.8" | June, 2012 | *Phaseolus vulgaris* | Sobrinho et al., 2014 |
| BR:Sag11:12 | KJ939789 | Santo Antônio de Góias, GO | S16°30'22.5" | W049°17'06.8" | June, 2012 | *Phaseolus vulgaris* | Sobrinho et al., 2014 |
| BR:Sag12:12 | KJ939790 | Santo Antônio de Góias, GO | S16°30'22.5" | W049°17'06.8" | June, 2012 | *Phaseolus vulgaris* | Sobrinho et al., 2014 |
| BR:Par14:12 | KJ939791 | Paranoá, DF | S16°00'47" | W047°33'21.5" | June, 2012 | *Phaseolus vulgaris* | Sobrinho et al., 2014 |
| BR:Par1:12 | KJ939792 | Paranoá, DF | S16°00'47" | W047°33'21.5" | June, 2012 | *Phaseolus vulgaris* | Sobrinho et al., 2014 |
| BR:Par2:12 | KJ939793 | Paranoá, DF | S16°00'47" | W047°33'21.5" | June, 2012 | *Phaseolus vulgaris* | Sobrinho et al., 2014 |
| BR:Par15:12 | KJ939794 | Paranoá, DF | S16°00'47" | W047°33'21.5" | June, 2012 | *Phaseolus vulgaris* | Sobrinho et al., 2014 |
| BR:Par16:12 | KJ939795 | Paranoá, DF | S16°00'47" | W047°33'21.5" | June, 2012 | *Phaseolus vulgaris* | Sobrinho et al., 2014 |
| BR:Par3:12 | KJ939796 | Paranoá, DF | S16°00'47" | W047°33'21.5" | June, 2012 | *Phaseolus vulgaris* | Sobrinho et al., 2014 |
| BR:Par17:12 | KJ939797 | Paranoá, DF | S16°00'47" | W047°33'21.5" | June, 2012 | *Phaseolus vulgaris* | Sobrinho et al., 2014 |
| BR:Par4:12 | KJ939798 | Paranoá, DF | S16°00'47" | W047°33'21.5" | June, 2012 | *Phaseolus vulgaris* | Sobrinho et al., 2014 |
| BR:Par5:12 | KJ939799 | Paranoá, DF | S16°00'47" | W047°33'21.5" | June, 2012 | *Phaseolus vulgaris* | Sobrinho et al., 2014 |
| BR:Par6:12 | KJ939800 | Paranoá, DF | S16°00'47" | W047°33'21.5" | June, 2012 | *Phaseolus vulgaris* | Sobrinho et al., 2014 |
| BR:Par18:12 | KJ939801 | Paranoá, DF | S16°00'47" | W047°33'21.5" | June, 2012 | *Phaseolus vulgaris* | Sobrinho et al., 2014 |
| BR:Par7:12 | KJ939802 | Paranoá, DF | S16°00'47" | W047°33'21.5" | June, 2012 | *Phaseolus vulgaris* | Sobrinho et al., 2014 |
| BR:Par19:12 | KJ939803 | Paranoá, DF | S16°00'47" | W047°33'21.5" | June, 2012 | *Phaseolus vulgaris* | Sobrinho et al., 2014 |
| BR:Par8:12 | KJ939804 | Paranoá, DF | S16°00'47" | W047°33'21.5" | June, 2012 | *Phaseolus vulgaris* | Sobrinho et al., 2014 |
| BR:Par20:12 | KJ939805 | Paranoá, DF | S16°00'47" | W047°33'21.5" | June, 2012 | *Phaseolus vulgaris* | Sobrinho et al., 2014 |
| BR:Par21:12 | KJ939806 | Paranoá, DF | S16°00'47" | W047°33'21.5" | June, 2012 | *Phaseolus vulgaris* | Sobrinho et al., 2014 |
| BR:Par22:12 | KJ939807 | Paranoá, DF | S16°00'47" | W047°33'21.5" | June, 2012 | *Phaseolus vulgaris* | Sobrinho et al., 2014 |
| BR:Par23:12 | KJ939808 | Paranoá, DF | S16°00'47" | W047°33'21.5" | June, 2012 | *Phaseolus vulgaris* | Sobrinho et al., 2014 |
| BR:Par9:12 | KJ939809 | Paranoá, DF | S16°00'47" | W047°33'21.5" | June, 2012 | *Phaseolus vulgaris* | Sobrinho et al., 2014 |
| BR:Par10:12 | KJ939810 | Paranoá, DF | S16°00'47" | W047°33'21.5" | June, 2012 | *Phaseolus vulgaris* | Sobrinho et al., 2014 |
| BR:Par24:12 | KJ939811 | Paranoá, DF | S16°00'47" | W047°33'21.5" | June, 2012 | *Phaseolus vulgaris* | Sobrinho et al., 2014 |
| BR:Par25:12 | KJ939812 | Paranoá, DF | S16°00'47" | W047°33'21.5" | June, 2012 | *Phaseolus vulgaris* | Sobrinho et al., 2014 |
| BR:Par26:12 | KJ939813 | Paranoá, DF | S16°00'47" | W047°33'21.5" | June, 2012 | *Phaseolus vulgaris* | Sobrinho et al., 2014 |

| | | | | | | | |
|---|---|---|---|---|---|---|---|
| BR:Par27:12 | KJ939814 | Paranoá, DF | S16°00'47" | W047°33'21.5" | June, 2012 | *Phaseolus vulgaris* | Sobrinho et al., 2014 |
| BR:Par11:12 | KJ939815 | Paranoá, DF | S16°00'47" | W047°33'21.5" | June, 2012 | *Phaseolus vulgaris* | Sobrinho et al., 2014 |
| BR:Par28:12 | KJ939816 | Paranoá, DF | S16°00'47" | W047°33'21.5" | June, 2012 | *Phaseolus vulgaris* | Sobrinho et al., 2014 |
| BR:Par29:12 | KJ939817 | Paranoá, DF | S16°00'47" | W047°33'21.5" | June, 2012 | *Phaseolus vulgaris* | Sobrinho et al., 2014 |
| BR:Par30:12 | KJ939818 | Paranoá, DF | S16°00'47" | W047°33'21.5" | June, 2012 | *Phaseolus vulgaris* | Sobrinho et al., 2014 |
| BR:Par12:12 | KJ939819 | Paranoá, DF | S16°00'47" | W047°33'21.5" | June, 2012 | *Phaseolus vulgaris* | Sobrinho et al., 2014 |
| BR:Par13:12 | KJ939820 | Paranoá, DF | S16°00'47" | W047°33'21.5" | June, 2012 | *Phaseolus vulgaris* | Sobrinho et al., 2014 |
| BR:Una1:12 | KJ939821 | Unaí, MG | S16°43'02.2" | W046°45'14" | June, 2012 | *Phaseolus vulgaris* | Sobrinho et al., 2014 |
| BR:Una2:12 | KJ939822 | Unaí, MG | S16°43'02.2" | W046°45'14" | June, 2012 | *Phaseolus vulgaris* | Sobrinho et al., 2014 |
| BR:Una3:12 | KJ939823 | Unaí, MG | S16°43'02.2" | W046°45'14" | June, 2012 | *Phaseolus vulgaris* | Sobrinho et al., 2014 |
| BR:Una4:12 | KJ939824 | Unaí, MG | S16°43'02.2" | W046°45'14" | June, 2012 | *Phaseolus vulgaris* | Sobrinho et al., 2014 |
| BR:Una5:12 | KJ939825 | Unaí, MG | S16°43'02.2" | W046°45'14" | June, 2012 | *Phaseolus vulgaris* | Sobrinho et al., 2014 |
| BR:Una6:12 | KJ939826 | Unaí, MG | S16°43'02.2" | W046°45'14" | June, 2012 | *Phaseolus vulgaris* | Sobrinho et al., 2014 |
| BR:Una7:12 | KJ939827 | Unaí, MG | S16°43'02.2" | W046°45'14" | June, 2012 | *Phaseolus vulgaris* | Sobrinho et al., 2014 |
| BR:Una8:12 | KJ939828 | Unaí, MG | S16°43'02.2" | W046°45'14" | June, 2012 | *Phaseolus vulgaris* | Sobrinho et al., 2014 |
| BR:Una9:12 | KJ939829 | Unaí, MG | S16°43'02.2" | W046°45'14" | June, 2012 | *Phaseolus vulgaris* | Sobrinho et al., 2014 |
| BR:Una10:12 | KJ939830 | Unaí, MG | S16°43'02.2" | W046°45'14" | June, 2012 | *Phaseolus vulgaris* | Sobrinho et al., 2014 |
| BR:Una11:12 | KJ939831 | Unaí, MG | S16°43'02.2" | W046°45'14" | June, 2012 | *Phaseolus vulgaris* | Sobrinho et al., 2014 |
| BR:Una12:12 | KJ939832 | Unaí, MG | S16°43'02.2" | W046°45'14" | June, 2012 | *Phaseolus vulgaris* | Sobrinho et al., 2014 |
| BR:Una13:12 | KJ939833 | Unaí, MG | S16°43'02.2" | W046°45'14" | June, 2012 | *Phaseolus vulgaris* | Sobrinho et al., 2014 |
| BR:Una14:12 | KJ939834 | Unaí, MG | S16°43'02.2" | W046°45'14" | June, 2012 | *Phaseolus vulgaris* | Sobrinho et al., 2014 |
| BR:Una15:12 | KJ939835 | Unaí, MG | S16°43'02.2" | W046°45'14" | June, 2012 | *Phaseolus vulgaris* | Sobrinho et al., 2014 |
| BR:Una16:12 | KJ939836 | Unaí, MG | S16°43'02.2" | W046°45'14" | June, 2012 | *Phaseolus vulgaris* | Sobrinho et al., 2014 |
| BR:Cri1:12 | KJ939837 | Cristalina, GO | S17°05'29.2" | W047°37'01.4" | June, 2012 | *Phaseolus vulgaris* | Sobrinho et al., 2014 |
| BR:Cri2:12 | KJ939838 | Cristalina, GO | S17°05'29.2" | W047°37'01.4" | June, 2012 | *Phaseolus vulgaris* | Sobrinho et al., 2014 |
| BR:Cri3:12 | KJ939839 | Cristalina, GO | S17°05'29.2" | W047°37'01.4" | June, 2012 | *Phaseolus vulgaris* | Sobrinho et al., 2014 |
| BR:Cri9:12 | KJ939840 | Cristalina, GO | S17°05'29.2" | W047°37'01.4" | June, 2012 | *Phaseolus vulgaris* | Sobrinho et al., 2014 |
| BR:Cri10:12 | KJ939841 | Cristalina, GO | S17°05'29.2" | W047°37'01.4" | June, 2012 | *Phaseolus vulgaris* | Sobrinho et al., 2014 |
| BR:Cri11:12 | KJ939842 | Cristalina, GO | S17°05'29.2" | W047°37'01.4" | June, 2012 | *Phaseolus vulgaris* | Sobrinho et al., 2014 |
| BR:Cri12:12 | KJ939844 | Cristalina, GO | S17°05'29.2" | W047°37'01.4" | June, 2012 | *Phaseolus vulgaris* | Sobrinho et al., 2014 |
| BR:Cri4:12 | KJ939844 | Cristalina, GO | S17°05'29.2" | W047°37'01.4" | June, 2012 | *Phaseolus vulgaris* | Sobrinho et al., 2014 |
| BR:Cri5:12 | KJ939845 | Cristalina, GO | S17°05'29.2" | W047°37'01.4" | June, 2012 | *Phaseolus vulgaris* | Sobrinho et al., 2014 |
| BR:Cri6:12 | KJ939846 | Cristalina, GO | S17°05'29.2" | W047°37'01.4" | June, 2012 | *Phaseolus vulgaris* | Sobrinho et al., 2014 |
| BR:Cri7:12 | KJ939847 | Cristalina, GO | S17°05'29.2" | W047°37'01.4" | June, 2012 | *Phaseolus vulgaris* | Sobrinho et al., 2014 |
| BR:Cri8:12 | KJ939848 | Cristalina, GO | S17°05'29.2" | W047°37'01.4" | June, 2012 | *Phaseolus vulgaris* | Sobrinho et al., 2014 |
| BR:Cri13:12 | KJ939849 | Cristalina, GO | S17°05'29.2" | W047°37'01.4" | June, 2012 | *Phaseolus vulgaris* | Sobrinho et al., 2014 |
| BR:Cri14:12 | KJ939850 | Cristalina, GO | S17°05'29.2" | W047°37'01.4" | June, 2012 | *Phaseolus vulgaris* | Sobrinho et al., 2014 |
| BR:Cri15:12 | KJ939851 | Cristalina, GO | S17°05'29.2" | W047°37'01.4" | June, 2012 | *Phaseolus vulgaris* | Sobrinho et al., 2014 |
| BR:Cri16:12 | KJ939852 | Cristalina, GO | S17°05'29.2" | W047°37'01.4" | June, 2012 | *Phaseolus vulgaris* | Sobrinho et al., 2014 |
| BR:Cri17:12 | KJ939853 | Cristalina, GO | S17°05'29.2" | W047°37'01.4" | June, 2012 | *Phaseolus vulgaris* | Sobrinho et al., 2014 |
| *Blainvillea yellow spot virus* (BlYSV) | | | | | | | |

| | | | | | | | | |
|---|---|---|---|---|---|---|---|---|
| BR:Vic04.1:10 | KC706516 | | Viçosa, MG* | S20°45'22.9" | W42°50'58.0" | May, 2010 | *Blainvillea rhomboidea* | Rocha et al., 2013 |
| BR:Vic04.2:10 | KC706517 | | Viçosa, MG | S20°45'22.9" | W42°50'58.0" | May, 2010 | *Blainvillea rhomboidea* | Rocha et al., 2013 |
| BR:Vic07:10 | | KC706523 | Viçosa, MG | S20°45'22.9" | W42°50'58.0" | May, 2010 | *Blainvillea rhomboidea* | Rocha et al., 2013 |
| BR:Vic08:10 | KC706518 | KC706524 | Viçosa, MG | S20°45'22.9" | W42°50'58.0" | May, 2010 | *Blainvillea rhomboidea* | Rocha et al., 2013 |
| BR:Vic09:10 | KC706519 | | Viçosa, MG | S20°45'22.9" | W42°50'58.0" | May, 2010 | *Blainvillea rhomboidea* | Rocha et al., 2013 |
| BR:Vic11:10 | KC706520 | KC706525 | Viçosa, MG | S20°45'22.9" | W42°50'58.0" | May, 2010 | *Blainvillea rhomboidea* | Rocha et al., 2013 |
| BR:Vic13:10 | KC706521 | | Viçosa, MG | S20°45'22.9" | W42°50'58.0" | May, 2010 | *Blainvillea rhomboidea* | Rocha et al., 2013 |
| BR:Vic18:10 | | KC706526 | Viçosa, MG | S20°45'22.9" | W42°50'58.0" | May, 2010 | *Blainvillea rhomboidea* | Rocha et al., 2013 |
| BR:Vic20:10 | KC706522 | | Viçosa, MG | S20°45'22.9" | W42°50'58.0" | May, 2010 | *Blainvillea rhomboidea* | Rocha et al., 2013 |
| BR:Vic21:10 | | KC706527 | Viçosa, MG | S20°45'22.9" | W42°50'58.0" | May, 2010 | *Blainvillea rhomboidea* | Rocha et al., 2013 |
| BR:Vic26s:10 | | KC706529 | Viçosa, MG | S20°45'22.9" | W42°50'58.0" | May, 2010 | *Blainvillea rhomboidea* | Rocha et al., 2013 |
| BR:Vic26c:10 | | KC706528 | Viçosa, MG | S20°45'22.9" | W42°50'58.0" | May, 2010 | *Blainvillea rhomboidea* | Rocha et al., 2013 |
| BR:Coi25:07 | EU710756 | EU710757 | Coimbra, MG | S20°51'29.9" | W042°51'42.2" | July, 2007 | *Blainvillea rhomboidea* | Rocha et al., 2013 |
| BR:Coi32.1:13 | MT626983 | | Coimbra, MG | S20°51'29.9" | W042°51'42.2" | July, 2013 | *Blainvillea rhomboidea* | Sande, 2014 |
| BR:Coi32.2:13 | MT626984 | | Coimbra, MG | S20°51'29.9" | W042°51'42.2" | July, 2013 | *Blainvillea rhomboidea* | Sande, 2014 |
| BR:Coi34:13 | MT626985 | | Coimbra, MG | S20°51'29.9" | W042°51'42.2" | July, 2013 | *Blainvillea rhomboidea* | Sande, 2014 |
| BR:Coi37.1:13 | MT626986 | | Coimbra, MG | S20°51'29.9" | W042°51'42.2" | July, 2013 | *Blainvillea rhomboidea* | Sande, 2014 |
| BR:Coi37.2:13 | MT626987 | | Coimbra, MG | S20°51'29.9" | W042°51'42.2" | July, 2013 | *Blainvillea rhomboidea* | Sande, 2014 |
| BR:Coi51:13 | MT626988 | | Coimbra, MG | S20°51'29.9" | W042°51'42.2" | July, 2013 | *Blainvillea rhomboidea* | Sande, 2014 |
| BR:Coi151:14 | MT626989 | | Coimbra, MG | S20°50'58.3" | W042°53'14.4" | February, 2014 | *Blainvillea rhomboidea* | Sande, 2014 |
| BR:Coi152.1:14 | MT626990 | | Coimbra, MG | S20°50'58.3" | W042°53'14.4" | February, 2014 | *Blainvillea rhomboidea* | Sande, 2014 |
| BR:Coi153:14 | MT626991 | | Coimbra, MG | S20°50'58.3" | W042°53'14.4" | February, 2014 | *Blainvillea rhomboidea* | Sande, 2014 |
| BR:Coi154:14 | MT626992 | | Coimbra, MG | S20°50'58.3" | W042°53'14.4" | February, 2014 | *Blainvillea rhomboidea* | Sande, 2014 |
| BR:Coi155:14 | MT626993 | | Coimbra, MG | S20°50'58.3" | W042°53'14.4" | February, 2014 | *Blainvillea rhomboidea* | Sande, 2014 |
| BR:Coi156:14 | MT626994 | | Coimbra, MG | S20°50'58.3" | W042°53'14.4" | February, 2014 | *Blainvillea rhomboidea* | Sande, 2014 |
| BR:Coi157:14 | MT626995 | | Coimbra, MG | S20°50'58.3" | W042°53'14.4" | February, 2014 | *Blainvillea rhomboidea* | Sande, 2014 |
| BR:Coi164:14 | MT626996 | | Coimbra, MG | S20°50'58.3" | W042°53'14.4" | February, 2014 | *Blainvillea rhomboidea* | Sande, 2014 |
| BR:Jun1:09 | JX871394 | | Junqueiro, AL | S9°54'24.6" | W36°28'30.1" | November, 2009 | *Blainvillea rhomboidea* | Tavares et al., 2012 |
| BR:Lim1:09 | JX871393 | | Limoeiro, AL | S9°46'05.4" | W36°30'41.6" | November, 2009 | *Blainvillea rhomboidea* | Tavares et al., 2012 |
| BR:Rla6:09 | JX871392 | | Rio Largo, AL | S9°28'43.7" | W35°49'44.7" | November, 2009 | *Blainvillea rhomboidea* | Tavares et al., 2012 |
| BR:Rla5:10 | JX871391 | | Rio Largo, AL | S9°28'43.7" | W35°49'44.7" | January, 2010 | *Blainvillea rhomboidea* | Tavares et al., 2012 |
| BR:Rla4:10 | JX871390 | | Rio Largo, AL | S9°28'43.7" | W35°49'44.7" | January, 2010 | *Blainvillea rhomboidea* | Tavares et al., 2012 |
| BR:Rla3:10 | JX871389 | | Rio Largo, AL | S9°28'43.7" | W35°49'44.7" | January, 2010 | *Blainvillea rhomboidea* | Tavares et al., 2012 |
| BgV06A.1.C80 | JF694468 | | Rio Largo, AL | S9°28'43.7" | W35°49'44.7" | February, 2012 | *Blainvillea rhomboidea* | Wyant et al., 2012 |
| BgV06A.1.C81 | JF694476 | | BA | S12°56'42.6" | W38°26'52.2" | February, 2012 | *Blainvillea rhomboidea* | Wyant et al., 2012 |
| BgV06B.1.C55 | | JF694470 | BA | S12°56'42.6" | W38°26'52.2" | February, 2012 | *Blainvillea rhomboidea* | Wyant et al., 2012 |
| BgV06B.1.C56 | | JF694470 | Rio Largo, AL | S9°28'43.7" | W35°49'44.7" | February, 2012 | *Blainvillea rhomboidea* | Wyant et al., 2012 |
| BgV06B.1.C70 | | JF694478 | Rio Largo, AL | S9°28'43.7" | W35°49'44.7" | February, 2012 | *Blainvillea rhomboidea* | Wyant et al., 2012 |
| BgV06B.1.C82 | | JF694469 | BA | S12°56'42.6" | W38°26'52.2" | February, 2012 | *Blainvillea rhomboidea* | Wyant et al., 2012 |
| *Euphorbia yellow mosaic virus* (EuYMV) | | | | | | | | |
| BR:GO:Ita5082:07 | JF756671 | JF756677 | Itaberaí, GO | S16°01'23.9" | W49°47'31.0" | 2007 | *Euphorbia heterophylla* | Mar et al., 2017 |

| | | | | | | | | |
|---|---|---|---|---|---|---|---|---|
| BR:Sag32:12 | KY559431 | | Santo Antônio de Goiás, GO | S16°30'22.50" | W49°17'06.80" | July, 2012 | *Euphorbia heterophylla* | Mar et al., 2017 |
| BR:Sag33:12 | KY559432 | KY559580 | Santo Antônio de Goiás, GO | S16°30'22.50" | W49°17'06.80" | July, 2012 | *Euphorbia heterophylla* | Mar et al., 2017 |
| BR:Sag35:12 | KY559433 | | Santo Antônio de Goiás, GO | S16°30'22.50" | W49°17'06.80" | July, 2012 | *Euphorbia heterophylla* | Mar et al., 2017 |
| BR:Sag34:12 | KY559434 | KY559581 | Santo Antônio de Goiás, GO | S16°30'22.50" | W49°17'06.80" | July, 2012 | *Euphorbia heterophylla* | Mar et al., 2017 |
| BR:Sag40.1:12 | | KY559582 | Santo Antônio de Goiás, GO | S16°30'22.50" | W49°17'06.80" | July, 2012 | *Euphorbia heterophylla* | Mar et al., 2017 |
| BR:Sag40.2:12 | | KY559583 | Santo Antônio de Goiás, GO | S16°30'22.50" | W49°17'06.80" | July, 2012 | *Euphorbia heterophylla* | Mar et al., 2017 |
| BR:Sag43:12 | KY559437 | KY559584 | Santo Antônio de Goiás, GO | S16°30'22.50" | W49°17'06.80" | July, 2012 | *Euphorbia heterophylla* | Mar et al., 2017 |
| BR:Sag44:12 | KY559438 | KY559585 | Santo Antônio de Goiás, GO | S16°30'22.50" | W49°17'06.80" | July, 2012 | *Euphorbia heterophylla* | Mar et al., 2017 |
| BR:Sag52:12 | KY559441 | KY559586 | Santo Antônio de Goiás, GO | S16°30'22.50" | W49°17'06.80" | July, 2012 | *Euphorbia heterophylla* | Mar et al., 2017 |
| BR:Sag53:12 | KY559442 | KY559587 | Santo Antônio de Goiás, GO | S16°30'22.50" | W49°17'06.80" | July, 2012 | *Euphorbia heterophylla* | Mar et al., 2017 |
| BR:Sag54:12 | KY559443 | KY559588 | Santo Antônio de Goiás, GO | S16°30'22.50" | W49°17'06.80" | July, 2012 | *Euphorbia heterophylla* | Mar et al., 2017 |
| BR:Sag56:12 | KY559444 | KY559589 | Santo Antônio de Goiás, GO | S16°30'22.50" | W49°17'06.80" | July, 2012 | *Euphorbia heterophylla* | Mar et al., 2017 |
| BR:Sag57:12 | KY559445 | KY559590 | Santo Antônio de Goiás, GO | S16°30'22.50" | W49°17'06.80" | July, 2012 | *Euphorbia heterophylla* | Mar et al., 2017 |
| BR:Sag58:12 | KY559446 | KY559591 | Santo Antônio de Goiás, GO | S16°30'22.50" | W49°17'06.80" | July, 2012 | *Euphorbia heterophylla* | Mar et al., 2017 |
| BR:Sag59:12 | KY559447 | KY559592 | Santo Antônio de Goiás, GO | S16°30'22.50" | W49°17'06.80" | July, 2012 | *Euphorbia heterophylla* | Mar et al., 2017 |
| BR:Sag59.1:12 | KY559448 | | Santo Antônio de Goiás, GO | S16°30'22.50" | W49°17'06.80" | July, 2012 | *Euphorbia heterophylla* | Mar et al., 2017 |
| BR:Sag61.1:12 | KY559449 | KY559593 | Santo Antônio de Goiás, GO | S16°30'22.50" | W49°17'06.80" | July, 2012 | *Euphorbia heterophylla* | Mar et al., 2017 |
| BR:Sag61.2:12 | | KY559594 | Santo Antônio de Goiás, GO | S16°30'22.50" | W49°17'06.80" | July, 2012 | *Euphorbia heterophylla* | Mar et al., 2017 |
| BR:MS:Mir2:07 | FN435997 | FN435998 | Miranda, MS | S20°15'41.9" | W56°22'14.9" | 2007 | *Euphorbia heterophylla* | Mar et al., 2017 |
| BR:MS:Mir:1:07 | FN435995 | FN435996 | Miranda, MS | S20°15'41.9" | W56°22'14.9" | 2007 | *Euphorbia heterophylla* | Mar et al., 2017 |
| BR:Gua183:09 | KY559475 | KY559599 | Guamiranga, PR | S25°11'15.28" | W50°52'36.98" | September, 2009 | *Euphorbia heterophylla* | Mar et al., 2017 |
| BR:Nom682:10 | KY559478 | KY559600 | Nova Mercedes, PR | S24°30'54.70" | W54°07'0.22" | June, 2010 | *Euphorbia heterophylla* | Mar et al., 2017 |
| BR:Caf777:10 | KY559482 | KY559602 | Cafelândia, PR | S24°39'21.30" | W53°13'07.40" | September, 2010 | *Euphorbia heterophylla* | Mar et al., 2017 |
| BR:Frb1019:11 | KY559484 | KY559603 | Francisco Beltrão, PR | S26°04'01" | W53°00'42" | April, 2011 | *Euphorbia heterophylla* | Mar et al., 2017 |
| BR:Frb1019.1:11 | | KY559604 | Francisco Beltrão, PR | S26°04'01" | W53°00'42" | April, 2011 | *Euphorbia heterophylla* | Mar et al., 2017 |
| BR:Can1033:11 | KY559486 | KY559605 | Candói, PR | S25°32'07.74" | W52°00'03.55" | April, 2011 | *Euphorbia heterophylla* | Mar et al., 2017 |
| BR:Amp1068:11 | KY559488 | KY559606 | Ampére, PR | S25°57'06.90" | W53°24'25" | April, 2011 | *Euphorbia heterophylla* | Mar et al., 2017 |
| BR:Ara1133:11 | KY559491 | KY559607 | Araruna, PR | S24°03'50" | W52°33'52" | April, 2011 | *Euphorbia heterophylla* | Mar et al., 2017 |
| BR:Ats03:09 | KY559502 | KY559608 | Almirante Tamandaré, RS | S28°06'35.02" | W52°54'29.57" | September, 2009 | *Euphorbia heterophylla* | Mar et al., 2017 |
| BR:Ats3.1:09 | | KY559609 | Almirante Tamandaré, RS | S28°06'35.02" | W52°54'29.57" | September, 2009 | *Euphorbia heterophylla* | Mar et al., 2017 |
| BR:Ats3.2:09 | | KY559610 | Almirante Tamandaré, RS | S28°06'35.02" | W52°54'29.57" | September, 2009 | *Euphorbia heterophylla* | Mar et al., 2017 |
| BR:Ats3.3:09 | | KY559611 | Almirante Tamandaré, RS | S28°06'35.02" | W52°54'29.57" | September, 2009 | *Euphorbia heterophylla* | Mar et al., 2017 |
| BR:Cha14:09 | KY559503 | KY559612 | Chapada, RS | S28°02'39.56" | W53°04'53.95" | March, 2009 | *Euphorbia heterophylla* | Mar et al., 2017 |
| BR:Cha14.3a:09 | KY559504 | | Chapada, RS | S28°02'39.56" | W53°04'53.95" | March, 2009 | *Euphorbia heterophylla* | Mar et al., 2017 |
| BR:Sta51:09 | KY559508 | KY559613 | Santo Ângelo, RS | S28°22'54.70" | W54°18'17.23" | March, 2009 | *Euphorbia heterophylla* | Mar et al., 2017 |
| BR:Smm61:09 | KY559509 | KY559614 | São Miguel das Missões, RS | S28°29'35.59" | W54°33'37.15" | March, 2009 | *Euphorbia heterophylla* | Mar et al., 2017 |
| BR:Smm61.1:09 | KY559510 | | São Miguel das Missões, RS | S28°29'35.59" | W54°33'37.15" | March, 2009 | *Euphorbia heterophylla* | Mar et al., 2017 |
| BR:Ats504:10 | KY559517 | KY559615 | Almirante Tamandaré, RS | S28°06'35.02305" | 52°54'29.57" | March, 2010 | *Euphorbia heterophylla* | Mar et al., 2017 |
| BR:Ats504.1:10 | | KY559616 | Almirante Tamandaré, RS | S28°06'35.02305" | 52°54'29.57" | March, 2010 | *Euphorbia heterophylla* | Mar et al., 2017 |
| BR:Cha510:10 | KY559518 | KY559617 | Chapada, RS | S28°02'39" | W53°04'53" | March, 2010 | *Euphorbia heterophylla* | Mar et al., 2017 |
| BR:Cha510.1:10 | KY559519 | KY559618 | Chapada, RS | S28°02'39" | W53°04'53" | March, 2010 | *Euphorbia heterophylla* | Mar et al., 2017 |

| | | | | | | | | |
|---|---|---|---|---|---|---|---|---|
| BR:Cha510.2:10 | | KY559619 | Chapada, RS | S28°02'39" | W53°04'53" | March, 2010 | *Euphorbia heterophylla* | Mar et al., 2017 |
| BR:Cha517-10 | KY559520 | | Chapada, RS | S28°02'39" | W53°04'53" | March, 2010 | *Euphorbia heterophylla* | Mar et al., 2017 |
| BR:Str536:10 | KY559522 | KY559620 | Santa Rosa, RS | S27°52'52.94" | W54°26'02.50" | March, 2010 | *Euphorbia heterophylla* | Mar et al., 2017 |
| BR:Cha920:11 | KY559525 | KY559621 | Chapada, RS | S28°02'39.56" | W53°04'53.95" | March, 2011 | *Euphorbia heterophylla* | Mar et al., 2017 |
| BR:Cha920.1:11 | KY559526 | KY559622 | Chapada, RS | S28°02'39.56" | W53°04'53.95" | March, 2011 | *Euphorbia heterophylla* | Mar et al., 2017 |
| BR:Cha925:11 | KY559530 | KY559623 | Chapada, RS | S28°01'08.80" | W53°05'52.31" | March, 2011 | *Euphorbia heterophylla* | Mar et al., 2017 |
| BR:Plm933:11 | KY559531 | KY559624 | Palmeira das Missões, RS | S27°45'57.35" | W53°27'29.49" | March, 2011 | *Euphorbia heterophylla* | Mar et al., 2017 |
| BR:Sau937:11 | KY559532 | KY559625 | Santo Augusto, RS | S27°44'12.44" | W53°51'27.91" | March, 2011 | *Euphorbia heterophylla* | Mar et al., 2017 |
| BR:Trm939:11 | KY559533 | KY559626 | Três de Maio, RS | S27°45'39.44" | W54°15'42.87" | March, 2011 | *Euphorbia heterophylla* | Mar et al., 2017 |
| BR:Str942:11 | KY559534 | KY559627 | Santa Rosa, RS | S27°45'39.44" | W54°15'42.87" | March, 2011 | *Euphorbia heterophylla* | Mar et al., 2017 |
| BR:Sta946:11 | KY559535 | KY559628 | Santo Ângelo, RS | S28°22'54.70" | W54°18'17.23" | March, 2011 | *Euphorbia heterophylla* | Mar et al., 2017 |
| BR:Pon1000:11 | KY559540 | KY559631 | Pontão, RS | S27°58'09.60" | W52°43'47.90" | April, 2011 | *Euphorbia heterophylla* | Mar et al., 2017 |
| BR:Non1003:11 | KY559541 | KY559632 | Nonoai, RS | S27°29'49.71" | W52°54'07.23" | April, 2011 | *Euphorbia heterophylla* | Mar et al., 2017 |
| BR:Cha1179:14 | KY559543 | KY559633 | Chapada, RS | S28°02'39.56" | W53°04'53.95" | July, 2014 | *Euphorbia heterophylla* | Mar et al., 2017 |
| BR:Cha1179.1:14 | KY559544 | KY559634 | Chapada, RS | S28°02'39.56" | W53°04'53.95" | July, 2014 | *Euphorbia heterophylla* | Mar et al., 2017 |
| BR:Cha1179.2:14 | KY559545 | KY559635 | Chapada, RS | S28°02'39.56" | W53°04'53.95" | July, 2014 | *Euphorbia heterophylla* | Mar et al., 2017 |
| BR:Cha1179.3:14 | KY559546 | KY559636 | Chapada, RS | S28°02'39.56" | W53°04'53.95" | July, 2014 | *Euphorbia heterophylla* | Mar et al., 2017 |
| *Macroptilium yellow spot virus* (MaYSV) | | | | | | | | |
| BR:Crb1:11 | KC004111 | | Craíbas, AL | S09°40'37.9" | W036°46'37.8" | July, 2011 | *Phaseolus vulgaris* | Sobrinho et al., 2014 |
| BR:Crb2:11 | KC004116 | | Craíbas, AL | S09°40'39.1" | W036°46'38.4" | July, 2011 | *Phaseolus vulgaris* | Sobrinho et al., 2014 |
| BR:Crb10:11 | KC004093 | | Craíbas, AL | S09°40'39.8" | W036°46'38.4" | July, 2011 | *Phaseolus vulgaris* | Sobrinho et al., 2014 |
| BR:Oaf8:11 | KC004121 | | Olho D'água das Flores, AL | S09°32'28.1" | W037°17'26.8" | July, 2011 | *Phaseolus vulgaris* | Sobrinho et al., 2014 |
| BR:Oaf28:11 | KJ939857 | | Olho D'água das Flores, AL | S09°32'27.8" | W037°17'22.1" | July, 2011 | *Macroptilium lathyroides* | Sobrinho et al., 2014 |
| BR:Oaf27:11 | KJ939856 | | Olho D'água das Flores, AL | S09°32'26.9" | W037°17'17.6" | July, 2011 | *Macroptilium lathyroides* | Sobrinho et al., 2014 |
| BR:Oaf24:11 | KC004132 | | Olho D'água das Flores, AL | S09°32'59.4" | W037°18'43.5" | July, 2011 | *Macroptilium lathyroides* | Sobrinho et al., 2014 |
| BR:Oaf25:11 | KC004133 | | Olho D'água das Flores, AL | S09°32'59.2" | W037°18'43.0" | July, 2011 | *Macroptilium lathyroides* | Sobrinho et al., 2014 |
| BR:Sti34:11 | KJ939891 | | Santana do Ipanema, AL | S09°23'25.1" | W037°12'46.9" | July, 2011 | *Phaseolus lunatus* | Sobrinho et al., 2014 |
| BR:Sti2:11 | KJ939860 | | Santana do Ipanema, AL | S09°23'24.7" | W037°12'48.0" | July, 2011 | *Phaseolus lunatus* | Sobrinho et al., 2014 |
| BR:Sti3:11 | KJ939861 | | Santana do Ipanema, AL | S09°23'24.8" | W037°12'48.0" | July, 2011 | *Phaseolus lunatus* | Sobrinho et al., 2014 |
| BR:Sti4:11 | KJ939862 | | Santana do Ipanema, AL | S09°23'24.9" | W037°12'48.0" | July, 2011 | *Phaseolus lunatus* | Sobrinho et al., 2014 |
| BR:Sti35:11 | KJ939892 | | Santana do Ipanema, AL | S09°23'24.9" | W037°12'48.3" | July, 2011 | *Phaseolus lunatus* | Sobrinho et al., 2014 |
| BR:Sti9:11 | KJ939867 | | Santana do Ipanema, AL | S09°23'24.9" | W037°12'47.5" | July, 2011 | *Phaseolus lunatus* | Sobrinho et al., 2014 |
| BR:Sti10:11 | KJ939868 | | Santana do Ipanema, AL | S09°23'24.9" | W037°12'47.6" | July, 2011 | *Phaseolus lunatus* | Sobrinho et al., 2014 |
| BR:Sti26:11 | KJ939883 | | Santana do Ipanema, AL | S09°23'24.9" | W037°12'47.6" | July, 2011 | *Phaseolus lunatus* | Sobrinho et al., 2014 |
| BR:Sti29:11 | KJ939886 | | Santana do Ipanema, AL | S09°23'24.8" | W037°12'48.6" | July, 2011 | *Phaseolus lunatus* | Sobrinho et al., 2014 |
| BR:PE:CAU:22_1 | KT779562 | | Caruaru, PE | S8°17'47.6" | W35°58'56.1" | August , 2013 | *Desmodium glabrum* | Fontenele et al., 2016 |
| BR:PE:CAU:22_2 | KT779561 | | Caruaru, PE | S8°17'47.6" | W35°58'56.1" | August , 2013 | *Desmodium glabrum* | Fontenele et al., 2016 |
| BR:PE:CAU:22_3 | | KT779560 | Caruaru, PE | S8°17'47.6" | W35°58'56.1" | August , 2013 | *Desmodium glabrum* | Fontenele et al., 2016 |
| BR:PE:CAU:22_4 | | KT779559 | Caruaru, PE | S8°17'47.6" | W35°58'56.1" | August , 2013 | *Desmodium glabrum* | Fontenele et al., 2016 |
| BR:PE:CAU:22_5 | | KT779558 | Caruaru, PE | S8°17'47.6" | W35°58'56.1" | August , 2013 | *Desmodium glabrum* | Fontenele et al., 2016 |
| BR:PE:CAU:23_1 | KT779565 | KT779563 | Caruaru, PE | S8°17'47.6" | W35°58'56.1" | August , 2013 | *Desmodium glabrum* | Fontenele et al., 2016 |

| | | | | | | | | |
|---|---|---|---|---|---|---|---|---|
| BR:PE:CAU:23_2 | KT779564 | KT779563 | Caruaru, PE | S8°17'47.6" | W35°58'56.1" | August , 2013 | *Desmodium glabrum* | Fontenele et al., 2016 |
| *Tomato severe rugose virus* (ToSRV) | | | | | | | | |
| BR:Coi70.1:13 | MT627045 | | Coimbra, MG | S20°51'43.2" | W042°51'27" | July, 2013 | *Solanum lycopersicum* | Sande, 2014 |
| BR:Coi70.2:13 | MT627046 | | Coimbra, MG | S20°51'43.2" | W042°51'27" | July, 2013 | *Solanum lycopersicum* | Sande, 2014 |
| BR:Coi70.3:13 | MT627047 | | Coimbra, MG | S20°51'43.2" | W042°51'27" | July, 2013 | *Solanum lycopersicum* | Sande, 2014 |
| BR:Coi73.2:13 | MT627048 | | Coimbra, MG | S20°51'43.2" | W042°51'27" | July, 2013 | *Solanum lycopersicum* | Sande, 2014 |
| BR:Coi73.4:13 | MT627049 | | Coimbra, MG | S20°51'43.2" | W042°51'27" | July, 2013 | *Solanum lycopersicum* | Sande, 2014 |
| BR:Coi78:13 | MT627050 | | Coimbra, MG | S20°51'43.2" | W042°51'27" | July, 2013 | *Solanum lycopersicum* | Sande, 2014 |
| BR:Coi79:13 | MT627051 | | Coimbra, MG | S20°51'43.2" | W042°51'27" | July, 2013 | *Solanum lycopersicum* | Sande, 2014 |
| BR:Coi80:13 | MT627052 | | Coimbra, MG | S20°51'43.2" | W042°51'27" | July, 2013 | *Solanum lycopersicum* | Sande, 2014 |
| BR:Coi83:13 | MT627053 | | Coimbra, MG | S20°51'43.2" | W042°51'27" | July, 2013 | *Solanum lycopersicum* | Sande, 2014 |
| BR:Coi84:13 | MT627054 | | Coimbra, MG | S20°51'43.2" | W042°51'27" | July, 2013 | *Solanum lycopersicum* | Sande, 2014 |
| BR:Coi85.1:13 | MT627055 | | Coimbra, MG | S20°51'43.2" | W042°51'27" | July, 2013 | *Solanum lycopersicum* | Sande, 2014 |
| BR:Coi85.2:13 | MT627056 | | Coimbra, MG | S20°51'43.2" | W042°51'27" | July, 2013 | *Solanum lycopersicum* | Sande, 2014 |
| BR:Coi86:13 | MT627057 | | Coimbra, MG | S20°51'43.2" | W042°51'27" | July, 2013 | *Solanum lycopersicum* | Sande, 2014 |
| BR:Coi88:13 | MT627058 | | Coimbra, MG | S20°51'43.2" | W042°51'27" | July, 2013 | *Solanum lycopersicum* | Sande, 2014 |
| BR:Coi91:13 | MT627059 | | Coimbra, MG | S20°51'43.2" | W042°51'27" | July, 2013 | *Solanum lycopersicum* | Sande, 2014 |
| BR:Coi92.2:13 | MT627060 | | Coimbra, MG | S20°51'43.2" | W042°51'27" | July, 2013 | *Solanum lycopersicum* | Sande, 2014 |
| BR:Coi93:13 | MT627061 | | Coimbra, MG | S20°51'43.2" | W042°51'27" | July, 2013 | *Solanum lycopersicum* | Sande, 2014 |
| BR:Coi95:13 | MT627062 | | Coimbra, MG | S20°51'43.2" | W042°51'27" | July, 2013 | *Solanum lycopersicum* | Sande, 2014 |
| BR:Coi100.1:13 | MT627063 | | Coimbra, MG | S20°51'43.2" | W042°51'27" | July, 2013 | *Solanum lycopersicum* | Sande, 2014 |
| BR:Coi100.2:13 | MT627064 | | Coimbra, MG | S20°51'43.2" | W042°51'27" | July, 2013 | *Solanum lycopersicum* | Sande, 2014 |
| BR:Coi103.1:13 | MT627065 | | Coimbra, MG | S20°51'43.2" | W042°51'27" | July, 2013 | *Solanum lycopersicum* | Sande, 2014 |
| BR:Coi107:13 | MT627066 | | Coimbra, MG | S20°51'43.2" | W042°51'27" | July, 2013 | *Solanum lycopersicum* | Sande, 2014 |
| BR:Coi109:13 | MT627067 | | Coimbra, MG | S20°51'43.2" | W042°51'27" | July, 2013 | *Solanum lycopersicum* | Sande, 2014 |
| BR:Coi110:13 | MT627068 | | Coimbra, MG | S20°51'43.2" | W042°51'27" | July, 2013 | *Solanum lycopersicum* | Sande, 2014 |
| BR:Coi114:13 | MT627069 | | Coimbra, MG | S20°51'43.2" | W042°51'27" | July, 2013 | *Solanum lycopersicum* | Sande, 2014 |
| BR:Coi117:13 | MT627070 | | Coimbra, MG | S20°51'43.2" | W042°51'27" | July, 2013 | *Solanum lycopersicum* | Sande, 2014 |
| BR:Coi119.1:14 | MT627071 | | Coimbra, MG | S20°36'39.3" | W042°25'58.9" | February, 2014 | *Solanum lycopersicum* | Sande, 2014 |
| BR:Coi120:14 | MT627072 | | Coimbra, MG | S20°36'39.3" | W042°25'58.9" | February, 2014 | *Solanum lycopersicum* | Sande, 2014 |
| BR:Coi122:14 | MT627073 | | Coimbra, MG | S20°36'39.3" | W042°25'58.9" | February, 2014 | *Solanum lycopersicum* | Sande, 2014 |
| BR:Coi125:14 | MT627074 | | Coimbra, MG | S20°36'39.3" | W042°25'58.9" | February, 2014 | *Solanum lycopersicum* | Sande, 2014 |
| BR:Coi127:14 | MT627075 | | Coimbra, MG | S20°36'39.3" | W042°25'58.9" | February, 2014 | *Solanum lycopersicum* | Sande, 2014 |
| BR:Coi131:14 | MT627076 | | Coimbra, MG | S20°36'39.3" | W042°25'58.9" | February, 2014 | *Solanum lycopersicum* | Sande, 2014 |
| BR:Coi172:14 | MT627077 | | Coimbra, MG | S20°36'39.3" | W042°25'58.9" | February, 2014 | *Solanum lycopersicum* | Sande, 2014 |
| BR:Coi179:14 | MT627078 | | Coimbra, MG | S20°36'39.3" | W042°25'58.9" | February, 2014 | *Solanum lycopersicum* | Sande, 2014 |
| BR:Coi182:14 | MT627079 | | Coimbra, MG | S20°36'39.3" | W042°25'58.9" | February, 2014 | *Solanum lycopersicum* | Sande, 2014 |
| BR:Coi183.1:14 | MT627080 | | Coimbra, MG | S20°36'39.3" | W042°25'58.9" | February, 2014 | *Solanum lycopersicum* | Sande, 2014 |
| BR:Coi183.2:14 | MT627081 | | Coimbra, MG | S20°36'39.3" | W042°25'58.9" | February, 2014 | *Solanum lycopersicum* | Sande, 2014 |
| BR:Flo01:14 | MT627082 | | Florestal, MG | S19°55'55.8" | W044°23'52.4" | June, 2014 | *Solanum lycopersicum* | Sande, 2014 |
| BR:Flo02:14 | MT627083 | | Florestal, MG | S19°55'55.8" | W044°23'52.4" | June, 2014 | *Solanum lycopersicum* | Sande, 2014 |

| | | | | | | | | |
|---|---|---|---|---|---|---|---|---|
| BR:Flo04.1:14 | MT627084 | | Florestal, MG | S19°55'55.8" | W044°23'52.4" | June, 2014 | *Solanum lycopersicum* | Sande, 2014 |
| BR:Flo06:14 | MT627085 | | Florestal, MG | S19°55'55.8" | W044°23'52.4" | June, 2014 | *Solanum lycopersicum* | Sande, 2014 |
| BR:Flo07:14 | MT627086 | | Florestal, MG | S19°55'55.8" | W044°23'52.4" | June, 2014 | *Solanum lycopersicum* | Sande, 2014 |
| BR:Flo09.1:14 | MT627087 | | Florestal, MG | S19°55'55.8" | W044°23'52.4" | June, 2014 | *Solanum lycopersicum* | Sande, 2014 |
| BR:Flo13:14 | MT627088 | | Florestal, MG | S19°55'55.8" | W044°23'52.4" | June, 2014 | *Solanum lycopersicum* | Sande, 2014 |
| BR:Flo14:14 | MT627089 | | Florestal, MG | S19°55'55.8" | W044°23'52.4" | June, 2014 | *Solanum lycopersicum* | Sande, 2014 |
| BR:Flo15:14 | MT627090 | | Florestal, MG | S19°55'55.8" | W044°23'52.4" | June, 2014 | *Solanum lycopersicum* | Sande, 2014 |
| BR:Flo16:13 | MT627091 | | Florestal, MG | S19°55'55.8" | W044°23'52.4" | June, 2013 | *Solanum lycopersicum* | Sande, 2014 |
| BR:Flo18:13 | MT627092 | | Florestal, MG | S19°55'55.8" | W044°23'52.4" | June, 2013 | *Solanum lycopersicum* | Sande, 2014 |
| BR:Flo19:13 | MT627093 | | Florestal, MG | S19°55'55.8" | W044°23'52.4" | June, 2013 | *Solanum lycopersicum* | Sande, 2014 |
| BR:Flo22:14 | MT627094 | | Florestal, MG | S19°55'55.8" | W044°23'52.4" | June, 2014 | *Solanum lycopersicum* | Sande, 2014 |
| BR:Flo23:14 | MT627095 | | Florestal, MG | S19°55'55.8" | W044°23'52.4" | June, 2014 | *Solanum lycopersicum* | Sande, 2014 |
| BR:Flo31:14 | MT627096 | | Florestal, MG | S19°55'55.8" | W044°23'52.4" | June, 2014 | *Solanum lycopersicum* | Sande, 2014 |
| BR:Flo37:14 | MT627097 | | Florestal, MG | S19°55'55.8" | W044°23'52.4" | June, 2014 | *Solanum lycopersicum* | Sande, 2014 |
| BR:Flo165:08 | KC004070 | | Florestal, MG | S19°52'25.4" | W44°25'00.6" | July, 2008 | *Solanum lycopersicum* | Rocha et al., 2013 |
| BR:Flo202:08 | KC004071 | | Florestal, MG | S19°52'25.4" | W44°25'00.6" | July, 2008 | *Solanum lycopersicum* | Rocha et al., 2013 |
| BR:Flo203:08 | KC004072 | | Florestal, MG | S19°52'25.4" | W44°25'00.6" | July, 2008 | *Solanum lycopersicum* | Rocha et al., 2013 |
| BR:Flo206:08 | KC004073 | | Florestal, MG | S19°52'25.4" | W44°25'00.6" | July, 2008 | *Solanum lycopersicum* | Rocha et al., 2013 |
| BR:Flo208:08 | KC004074 | | Florestal, MG | S19°52'25.4" | W44°25'00.6" | July, 2008 | *Solanum lycopersicum* | Rocha et al., 2013 |
| BR:Car214:08 | KC004075 | | Carandaí, MG | S20°56'56.5" | W43°47'42.2" | July, 2008 | *Solanum lycopersicum* | Rocha et al., 2013 |
| BR:Car218.1:08 | KC004076 | | Carandaí, MG | S20°56'56.5" | W43°47'42.2" | July, 2008 | *Solanum lycopersicum* | Rocha et al., 2013 |
| BR:Car219.10:08 | KC004077 | | Carandaí, MG | S20°56'56.5" | W43°47'42.2" | July, 2008 | *Solanum lycopersicum* | Rocha et al., 2013 |
| BR:Car220:08 | KC004078 | | Carandaí, MG | S20°56'56.5" | W43°47'42.2" | July, 2008 | *Solanum lycopersicum* | Rocha et al., 2013 |
| BR:Car224:08 | KC004079 | | Carandaí, MG | S20°56'56.5" | W43°47'42.2" | July, 2008 | *Solanum lycopersicum* | Rocha et al., 2013 |
| BR:Car226.3:08 | KC004080 | | Carandaí, MG | S20°56'56.5" | W43°47'42.2" | July, 2008 | *Solanum lycopersicum* | Rocha et al., 2013 |
| BR:Car227:08 | KC004081 | | Carandaí, MG | S20°56'56.5" | W43°47'42.2" | July, 2008 | *Solanum lycopersicum* | Rocha et al., 2013 |
| BR:Car228:08 | KC004082 | | Carandaí, MG | S20°56'56.5" | W43°47'42.2" | July, 2008 | *Sida sp.* | Rocha et al., 2013 |
| BR:Car230:08 | KC004083 | | Carandaí, MG | S20°56'56.5" | W43°47'42.2" | July, 2008 | *Solanum lycopersicum* | Rocha et al., 2013 |
| BR:Car232:08 | KC004084 | | Carandaí, MG | S20°56'56.5" | W43°47'42.2" | July, 2008 | *Solanum lycopersicum* | Rocha et al., 2013 |
| BR:Car233:08 | KC004085 | | Carandaí, MG | S20°56'56.5" | W43°47'42.2" | July, 2008 | *Solanum lycopersicum* | Rocha et al., 2013 |
| BR:Car235:08 | KC004086 | KC706625 | Carandaí, MG | S20°56'56.5" | W43°47'42.2" | July, 2008 | *Solanum lycopersicum* | Rocha et al., 2013 |
| BR:Car236.1:08 | KC004087 | | Carandaí, MG | S20°56'56.5" | W43°47'42.2" | July, 2008 | *Solanum lycopersicum* | Rocha et al., 2013 |
| BR:Car237.6:08 | KC004088 | KC706626 | Carandaí, MG | S20°56'56.5" | W43°47'42.2" | July, 2008 | *Solanum lycopersicum* | Rocha et al., 2013 |
| BR:Car238:08 | KC004089 | KC706627 | Carandaí, MG | S20°56'56.5" | W43°47'42.2" | July, 2008 | *Solanum lycopersicum* | Rocha et al., 2013 |
| BR:Car237:08 | KC706620 | KC706624 | Carandaí, MG | S20°56'56.5" | W43°47'42.2" | July, 2008 | *Solanum lycopersicum* | Rocha et al., 2013 |
| BR:Car217.6:08 | | KC706621 | Carandaí, MG | S20°56'56.5" | W43°47'42.2" | July, 2008 | *Solanum lycopersicum* | Rocha et al., 2013 |
| BR:Car223:08 | | KC706622 | Carandaí, MG | S20°56'56.5" | W43°47'42.2" | July, 2008 | *Solanum lycopersicum* | Rocha et al., 2013 |
| BR:Car234.5:08 | | KC706623 | Carandaí, MG | S20°56'56.5" | W43°47'42.2" | July, 2008 | *Solanum lycopersicum* | Rocha et al., 2013 |

*State sigla: AL, Alagoas; BA, Bahia; DF, Federal District; GO, Goiás; MG, Minas Gerais; MS, Mato Grosso do Sul; PE, Pernambuco; PR, Paraná; RS, Rio Grande do Sul

**Supplementary Table S3**. Recombination events detected in the DNA-A and DNA-B components of *Bean golden mosaic virus* (BGMV), *Blainvillea yellow spot virus* (BlYSV), *Euphorbia yellow mosaic virus* (EuYMV), *Macroptilium yellow spot virus* (MaYSV) and *Tomato severe rugose virus* (ToSRV).

| **Virus** | **Component** | **Event** | **Recombinant** | **Recombination breakpoints*** | | **Parents** | | **Method**[#] | ***P*-value**[&] |
|---|---|---|---|---|---|---|---|---|---|
| | | | | **Begin** | **End** | **Minor** | **Major** | | |
| BGMV | DNA-B | 1 | BR:Cri15:12 | 179 | 53 | Unknown | BR:Cri14:12 | GMCS**3** | $5.147 \times 10^{-30}$ |
| | | 2 | BR:Cri13.2:12 | 1013 | 2379 | BR:Una10.2:12 | BR:Par8:12 | MC**S**3 | $2.712 \times 10^{-11}$ |
| | | 3 | BR:Una16:12 | 988 | 2368 | BR:Una10.2:12 | BR:Una2:12 | MCS**3** | $1.629 \times 10^{-03}$ |
| | | 4 | BR:Mur12.1:11, BR:Mur12.2:11 | 544 | 2330 | Unknown | BR:Mur3:11 | RGMCS**3** | $1.748 \times 10^{-18}$ |
| BlYSV | DNA-A | 1 | BgV06A.1.C80 | 642 | 2059 | BR:Lim1:09 | BgV06A.1.C81 | GMC**S**3 | $5.488 \times 10^{-06}$ |
| | | 2 | BR:Coi32.2:13, BR:Coi32.1:13, BR:Rla6:09, BR:Rla5:10, BR:Rla4:10, BR:Rla3:10, BgV06A.1.C81 | 2403 | 2641 | Unknown | BR:Vic04.2:10 | RGMC**S**3 | $5.346 \times 10^{-07}$ |
| | | 3 | BgV06A.1.C80, BR:Jun1:09, BR:Lim1:09, BR:Rla6:09 | 2060(?) | 202 | Unknown | BR:Coi37.2:13 | GMC**S**3 | $5.488 \times 10^{-06}$ |
| | DNA-B | 1 | BR:Vic07:10, BR:Vic04.1:10, BR:Vic08:10, BR:Vic09:10, BR:Vic18:10, BR:Vic21:10, BR:Coi25:07, BR:Vic26c:10, BR:Vic26s:10, BR:Coi165:14, BR:Coi164:14, BR:Coi155:14, BR:Coi154:14, BR:Coi150:14, BR:Coi145:14, BR:Coi37.1:13, BR:Coi36:13, BR:Coi33:13, BR:Coi32.1:13 | 2578 | 1865 | Unknown | BR:Vic13:10 | RGMC**S**3 | $3.364 \times 10^{-22}$ |
| | | 2 | BR:Coi33:13, BR:Coi142.1:14, BR:Coi151:14, BR:Coi152.1:14, BR:Coi155:14, BR:Coi154:14 | 2577(?) | 464 | BR:Coi158s:14 | BR:Vic07:10 | RGMC**3** | $1.345 \times 10^{-07}$ |
| | | 3 | BR:Coi148:14, BR:Coi158b:14, BR:Coi158s:14 | 1970 | 2485 | BR:Vic11:10 | BR:Coi152.2:14 | **R**GMCS3 | $1.250 \times 10^{-09}$ |

| | | | | | | | | | |
|---|---|---|---|---|---|---|---|---|---|
| | | 4 | BR:Coi157:14 | 1372 | 2512 | BR:Coi160.2:14 | BR:Coi35:13 | RGMC**S**3 | $3.330 \times 10^{-14}$ |
| | | 5 | BR:Vic08:10, BR:Vic04.1:10, BR:Vic07:10, BR:Vic09:10, BR:Vic18:10, BR:Vic21:10, BR:Coi25:07, BR:Vic26c:10, BR:Vic26s:10, BR:Coi36:13, BR:Coi37.1:13, BR:Coi164:14, BR:Coi165:14 | 2579(?) | 248 | BgV06B.1.C56 | BR:Coi158s:14 | **R**GMC | $5.498 \times 10^{-05}$ |
| | | 6 | BR:Vic11:10, BR:Vic13:10, BR:Coi34:13, BR:Coi35:13, BR:Coi142.2:14, BR:Coi160.2:14, BR:Coi157:14, BR:Coi152.2:14, BR:Coi148:14, BR:Coi147:14 | 292 | 1029 | Unknown | BR:Coi164:14 | GMC**S**3 | $3.187 \times 10^{-19}$ |
| | | 7 | BR:Vic07:10, BR:Coi33:13, BR:Coi164:14 | 249 | 1034 | BR:Coi25:07 | BR:Coi145:14 | R**M**S3 | $7.821 \times 10^{-06}$ |
| | | 8 | BR:Coi145:14, BR:Coi32.1:13, BR:Coi150:14, BR:Coi154:14 | 320 | 1483(?) | Unknown | BR:Coi158s:14 | GMC**3** | $2.106 \times 10^{-03}$ |
| | | 9 | BR:Coi160.2:14, BR:Coi34:13, BR:Coi51:13, BR:Coi157:14, BR:Coi160.1:14, BR:Coi159:14 | 2422 | 2575(?) | Unknown | BR:Coi152.2:14 | RMC**S**3 | $4.049 \times 10^{-05}$ |
| EuYMV | DNA-B | 1 | BR:Frb1019:11, BR:Frb1019.1:11, | 212 | 833 | BR:Nom682.2:10 | BR:Str942:11 | RGMSC**3** | $2.478 \times 10^{-08}$ |
| | | 2 | BR:Can1033:11 | 58 | 2215 | Unknown | BR:Gua183:09 | MCS**3** | $1.892 \times 10^{-05}$ |
| | | 3 | BR:Sag61.1:12 | 1494 | 2564 | BR:Sag44:12 | BR:Sag40.2:12 | RM**S**3 | $4.046 \times 10^{-08}$ |
| | | 4 | BR:Cha1179.1:14, BR:Cha510.2:10, BR:Cha1179.2:14, | 1306 | 1956 | Unknown | BR:Sau937:11 | RGMC**S**3 | $2.632 \times 10^{-08}$ |
| | | 5 | BR:Sag34:12, BR:Sag40.1:12, BR:Sag40.2:12, BR:Sag43:12, BR:Sag44:12, BR:Sag52:12, BR:Sag53:12, BR:Sag54:12, BR:Sag56:12, BR:Sag57:12, BR:Sag58:12, BR:Sag59:12, | 1101 | 1369 | Unknown | BR:Trm939:11 | GM**S**3 | $1.116 \times 10^{-03}$ |

| | | | | | | | | | |
|---|---|---|---|---|---|---|---|---|---|
| | | | BR:Sag61.1:12, BR:Sag61.2:12, BR:Ats3.2:09 | | | | | | |
| MaYSV | DNA-A | 1 | BR:Crb2:11, BR:Crb1:11, BR:Sti4:11, BR:Sti29:11, BR:Sti34:11 | 1853 | 2643 | BR:PE:CAU:23:2 | BR:Crb10:11 | RGBMC**S**3 | $1.032 \times 10^{-35}$ |
| | | 2 | BR:Oaf25:11 | 1893 | 2619 | Unknown | BR:Sti3:11 | RGMC**S**3 | $5.847 \times 10^{-35}$ |
| | | 3 | BR:Oaf8:11 | 1941 | 401 | Unknown | BR:Oaf24:11 | RGMC**S**3 | $9.771 \times 10^{-33}$ |
| | | 4 | BR:Sti2:11 | 2626 | 1979 | Unknown | BR:Crb10:11 | RGMCS**3** | $1.540 \times 10^{-11}$ |
| | | 5 | BR:Oaf8:11, BR:Oaf25:11, BR:Sti2:11 | 1941(?) | 2157 | Unknown | BR:Sti10:11 | RGM**C**3 | $5.057 \times 10^{-11}$ |
| | | 6 | BR:PE:CAU:23:2, BR:PE:CAU:22:1, BR:PE:CAU:22:2, BR:PE:CAU:23:1 | 988 | 1763 | Unknown | BR:Crb10:11 | RGBMC**S**3 | $2.581 \times 10^{-41}$ |
| | | 7 | BR:Sti35:11, BR:Sti3:11, BR:Sti9:11, BR:Sti26:11 | 1326 | 233 | BR:Oaf27:11 | BR:Sti10:11 | MC**S**3 | 8.640 x 10-07 |
| | DNA-B | 1 | BR:Crb2:11, BR:Crb1:11 | 2596 | 2075 | Unknown | BR:Crb10:11 | RGMCS**3** | $1.035 \times 10^{-24}$ |
| | | 2 | BR:Sti34:11, BR:Sti4:11 | 1452 | 2506 | BR:Oaf8:11 | Unknown | RGMC**S**3 | $1.379 \times 10^{-13}$ |
| | | 3 | BR:Sti29:11, BR:Sti4:11, BR:PE:CAU:22:3, BR:PE:CAU:22:4, BR:PE:CAU:22:5 | 2427 | 2596 | Unknown | BR:Oaf24:11 | RGMCS**3** | $9.343 \times 10^{-19}$ |
| | | 4 | BR:Sti2:11 | 1553 | 2595 | BR:Oaf8:11 | BR:Sti26:11 | | |
| | | 5 | BR:Sti4:11, BR:Sti10:11 | 2061 | 2444(?) | BR:Sti35:11 | BR:Sti29:11 | RMS**3** | $1.977 \times 10^{-09}$ |
| | | 6 | BR:Sti29:11, BR:Oaf24:11, BR:Oaf25.1:11, BR:Oaf25.2:11 | 2597(?) | 673 | BR:Sti2:11 | BR:Sti4:11 | RGBMCS**3** | $1.397 \times 10^{-07}$ |
| | | 7 | BR:Sti9.3:11, BR:Sti9.1:11, BR:Sti9.2:11 | 2329 | 2423 | BR:Oaf27:11 | Unknown | **R**GMCS3 | $6.316 \times 10^{-09}$ |
| | | 8 | BR:Sti34:11 | 406 | 1451(?) | BR:Oaf27:11 | Unknown | RGMCS**3** | $1.511 \times 10^{-09}$ |
| | | 9 | BR:PE:CAU:23:1 | 1996 | 2434(?) | BR:PE:CAU:22:5 | BR:Oaf24:11 | RMCS**3** | $2.469 \times 10^{-09}$ |
| | | 10 | BR:Sti9.3:11 | 2597 | 537 | Unknown | BR:Sti9.2:11 | RMCS**3** | $1.046 \times 10^{-08}$ |
| | | 11 | BR:Oaf28:11 | 1143 | 2328 | BR:Sti3:11 | BR:Oaf24:11 | RMS**3** | $5.766 \times 10^{-08}$ |

| | | | | | | | | | |
|---|---|---|---|---|---|---|---|---|---|
| | | 12 | BR:Oaf28:11, BR:Oaf24:11, BR:Oaf25.1:11,BR:Oaf27:11, BR:Oaf25.2:11 | 2329(?) | 2430 | Unknown | BR:Sti3:11 | RG**<u>M</u>**S3 | $6.789 \times 10^{-06}$ |
| | | 13 | BR:Sti10:11 | 340 | 685 | BR:Oaf8:11 | BR:Sti4:11 | RGMCS**<u>3</u>** | $4.572 \times 10^{-06}$ |
| ToSRV | DNA-B | 1 | BR:Flo01:14 | 232 | 1276 | BR:Coi183:13 | BR:Flo18:14 | GMC**<u>3</u>** | $2.156 \times 10^{-14}$ |

[*] Numbering starts at the first nucleotide after the cleavage site at the origin of replication and increase clockwise. (?), Breakpoints could not be accurately located.

[#] R, Rdp; G, Geneconv; B, Boostcan; M, Maxichi; C, Chimaera; S, Siscan; 3, 3Seq.

[&] The reported *P*-value is from the method in bold and underlined, and is the lowest *P*-value calculated for the featured event.

**Supplementary Table S4.** Parameters used in the Discriminant Analysis of Principal Components (DAPC) analysis for *Bean golden mosaic virus* (BGMV), *Euphorbia yellow mosaic virus* (EuYMV), *Macroptilium yellow spot virus* (MaYSV) and *Tomato severe rugose virus* (ToSRV) data sets.

| | | | Discriminant Analysis (DA) | | |
|---|---|---|---|---|---|
| **Virus** | **Component** | **K*** | **Number of retained principal components (PCs)** | **Number of retained discriminant functions** | **Proportion of conserved variance (%)** |
| BGMV | DNA-A | 5 | 6 | 3 | 96.97 |
| | DNA-B | 4 | 6 | 3 | 90.93 |
| EuYMV | DNA-A | 3 | 4 | 2 | 61.32 |
| | DNA-B | 3 | 10 | 2 | 67.51 |
| MaYSV | DNA-A[#] | 4 | 3 | 3 | 83.75 |
| | DNA-A[&] | 2 | 5 | 1 | 86.45 |
| ToSRV | DNA-A | 4 | 8 | 3 | 74.12 |
| | DNA-B | 4 | 7 | 3 | 74.73 |

* Number of clusters inferred by the *k*-means algorithm based on the Bayesian Information Criterion (BIC) used for DAPC analysis.

[#] Analysis performed considering recombination events.

[&] Analysis performed excluding recombination events.

**Supplementary Table S5.** Genetic variability indices of the DNA-A data sets of *Bean golden mosaic virus* (BGMV), *Blainvillea yellow spot virus* (BlYSV), *Euphorbia yellow mosaic virus* (EuYMV), *Macroptilium yellow spot virus* (MaYSV) and *Tomato severe rugose virus* (ToSRV).

| **Population** | **N*** | **h** | **Hd** | **S** | **DNA-A π** | **CP π** | **Rep π** | **Trap π** | **Ren π** | **AC4 π** | **IR-A π** |
|---|---|---|---|---|---|---|---|---|---|---|---|
| BGMV (Total) | 117 | 110 | 0.998 | 439 | 0.03965 | 0.03758 | 0.03090 | 0.02156 | 0.02337 | 0.02337 | 0.09498 |
| MG-1 | 13 | 10 | 0.923 | - | 0.00183 | 0.00204 | 0.00073 | 0.00118 | 0.00347 | 0.00347 | 0.00237 |
| MW-1 | 42 | 41 | 0.999 | - | 0.00323 | 0.00211 | 0.00221 | 0.00134 | 0.00084 | 0.00084 | 0.01172 |
| MW-2 | 33 | 33 | 1.000 | - | 0.00283 | 0.00142 | 0.00205 | 0.00031 | 0.00131 | 0.00131 | 0.01103 |
| AL-1 | 18 | 17 | 0.993 | - | 0.00300 | 0.00274 | 0.00205 | 0.00199 | 0.00306 | 0.00306 | 0.00732 |
| AL-2 | 11 | 9 | 0.945 | - | 0.00107 | 0.00178 | 0.00102 | 0.00000 | 0.00046 | 0.00046 | 0.00056 |
| BlYSV (Total) | 30 | 26 | 0.991 | 433 | 0.03742 | 0.03334 | 0.03408 | 0.03616 | 0.03006 | 0.02831 | 0.05812 |
| EuYMV (Total) | 50 | 46 | 0.995 | 407 | 0.02319 | 0.01949 | 0.02546 | 0.01752 | 0.01510 | 0.01740 | 0.03648 |
| GO | 17 | 15 | 0.985 | - | 0.01067 | 0.00984 | 0.01112 | 0.01127 | 0.00944 | 0.00879 | 0.02800 |
| SO-1 | 25 | 25 | 1.000 | - | 0.01738 | 0.01761 | 0.01867 | 0.01193 | 0.01139 | 0.01223 | 0.03147 |
| SO-2 | 8 | 7 | 0.964 | - | 0.01120 | 0.00952 | 0.01023 | 0.00952 | 0.01038 | 0.01495 | 0.03539 |
| MaYSV (Total) | 21 | 20 | 0.995 | 506 | 0.07170 | 0.04451 | 0.11122 | 0.02773 | 0.02991 | 0.18584 | 0.06927 |
| AL-1 | 9 | 9 | 1.000 | - | 0.01712 | 0.01745 | 0.01258 | 0.02399 | 0.02283 | 0.00991 | 0.02207 |
| AL-2 | 3 | 3 | 1.000 | - | 0.02683 | 0.01058 | 0.03438 | 0.03248 | 0.03175 | 0.02067 | 0.03272 |
| AL-3 | 5 | 5 | 1.000 | - | 0.01472 | 0.01534 | 0.00802 | 0.01538 | 0.02155 | 0.00388 | 0.02515 |
| PE | 4 | 3 | 0.833 | - | 0.00077 | 0.00066 | 0.00048 | 0.00128 | 0.00125 | 0.00194 | 0.00153 |
| ToSRV (Total) | 74 | 62 | 0.993 | 268 | 0.01062 | 0.01111 | 0.00965 | 0.00548 | 0.00759 | 0.01208 | 0.01808 |
| MG-1 | 5 | 5 | 1.000 | - | 0.00803 | 0.00767 | 0.00812 | 0.00256 | 0.00301 | 0.00455 | 0.01584 |
| MG-2 | 16 | 13 | 0.967 | - | 0.00315 | 0.00306 | 0.00278 | 0.00263 | 0.00203 | 0.00574 | 0.00542 |
| MG-3 | 23 | 16 | 0.945 | - | 0.00558 | 0.00282 | 0.00455 | 0.00518 | 0.00506 | 0.00680 | 0.01630 |
| MG-4 | 30 | 28 | 0.995 | - | 0.00450 | 0.00320 | 0.00384 | 0.00367 | 0.00590 | 0.00515 | 0.00897 |

* N, Number of sequences; h, haplotype number; Hd, haplotype diversity; S, number of segregating sites; π, average pairwise number of nucleotide differences per site (nucleotide diversity).

**Supplementary Table S6.** Genetic variability indices of the DNA-B data sets of *Bean golden mosaic virus* (BGMV), *Blainvillea yellow spot virus* (BlYSV), *Euphorbia yellow mosaic virus* (EuYMV), *Macroptilium yellow spot virus* (MaYSV) and *Tomato severe rugose virus* (ToSRV).

| **Population** | **N*** | **h** | **Hd** | **S** | **DNA-B π** | **MP π** | **NSP π** | **LIR-B π** | **SIR-B π** |
|---|---|---|---|---|---|---|---|---|---|
| BGMV (Total) | 123 | 104 | 0.997 | 628 | 0.05454 | 0.04015 | 0.0408 | 0.08662 | 0.11303 |
| MG-1 | 13 | 11 | 0.962 | - | 0.00226 | 0.00174 | 0.0002 | 0.00374 | 0.00237 |
| MW | 76 | 62 | 0.991 | - | 0.01061 | 0.00679 | 0.0092 | 0.01779 | 0.02761 |
| AL-1 | 19 | 17 | 0.982 | - | 0.00623 | 0.00480 | 0.0096 | 0.01065 | 0.00936 |
| AL-2 | 15 | 14 | 0.990 | - | 0.01647 | 0.01775 | 0.0126 | 0.01496 | 0.05714 |
| BlYSV (Total) | 41 | 39 | 0.998 | 840 | 0.07552 | 0.05053 | 0.07761 | 0.09226 | 0.13834 |
| EuYMV (Total) | 53 | 48 | 0.996 | 677 | 0.04000 | 0.02561 | 0.03650 | 0.05375 | 0.09018 |
| GO | 15 | 13 | 0.971 | - | 0.02683 | 0.02139 | 0.01794 | 0.03870 | 0.05090 |
| South-1 | 28 | 27 | 0.997 | - | 0.02757 | 0.01816 | 0.02720 | 0.03729 | 0.04309 |
| South-2 | 10 | 8 | 0.956 | - | 0.03953 | 0.02717 | 0.03891 | 0.05224 | 0.05333 |
| MaYSV (Total) | 24 | 20 | 0.982 | 503 | 0.05567 | 0.03237 | 0.03948 | 0.08973 | 0.13541 |
| ToSRV (Total) | 74 | 72 | 0.999 | 434 | 0.02143 | 0.01134 | 0.01073 | 0.03886 | 0.05075 |
| MG-1 | 5 | 5 | 1.000 | - | 0.01224 | 0.0066 | 0.01608 | 0.01557 | 0.00488 |
| MG-2 | 22 | 20 | 0.991 | - | 0.00709 | 0.00391 | 0.00247 | 0.01454 | 0.00565 |
| MG-3 | 19 | 19 | 1.000 | - | 0.00452 | 0.0376 | 0.00349 | 0.01267 | 0.01429 |
| MG-4 | 28 | 28 | 1.000 | - | 0.01012 | 0.00532 | 0.00466 | 0.01563 | 0.02202 |

* N, Number of sequences; h, haplotype number; Hd, haplotype diversity; S, number of segregating sites; π, average pairwise number of nucleotide differences per site (nucleotide diversity).

**Supplementary Table S7.** Reassortment events detected in *Bean golden mosaic virus* (BGMV), *Blainvillea yellow spot virus* (BlYSV), *Euphorbia yellow mosaic virus* (EuYMV), *Macroptilium yellow spot virus* (MaYSV) and *Tomato severe rugose virus* (ToSRV) data sets.

| **Virus** | **Event** | **Breakpoints*** | | **Reassortants** | **Parents** | | **Method#** | ***P*-value&** |
|---|---|---|---|---|---|---|---|---|
| | | **Begin** | **End** | | **Minor** | **Major** | | |
| BGMV | 1 | 2618 | 403 | BR:Cri13.2:12, BR:Una4:12, BR:Una16:12, BR:Cri11:12 | Unknown | BR-Par4-12 | MC**S**3 | $1.98 \times 10^{-10}$ |
| | 2 | 35 | 2593 | BR:Par19:12, BR:Una2:12, BR:Una3:12, BR:Una9:12, BR:Una12:12, BR:Cri9:12, BR:Cri10:12, BR:Cri12:12, BR:Cri13.1:12, BR:Cri14:12, BR:Cri15:12, BR:Cri16:12, BR:Sag5:12, BR:Sag8:12, BR:Par1:12, BR:Par2.1:12, BR:Par2.2:12, BR:Par4:12, BR:Par5.1:12, BR:Par5.2:12, BR:Par9.1:12, BR:Par9.2:12, BR:Par10:12, BR:Par16.1:12, BR:Par16.2:12, BR:Par18:12, BR:Par20.1:12, BR:Par20.2:12, BR:Par21:12, BR:Par24:12, BR:Par25:12 | Unknown | BR-Una6.1-12 | RGBMCS**3** | $4.65 \times 10^{-6}$ |
| BlYSV | 1 | 5305 | 2657 | BgV06A.1.C80:1 | BgV06A.1.C80:2 | BgV06A.1.C81 | GBMCS**3** | $5.85 \times 10^{-69}$ |
| | 2 | 5282 | 2658 | BR:Vic13:10 | Unknown | BR:Vic11:10 | RGBMC**S** | $1.19 \times 10^{-42}$ |
| | 3 | 5201 | 2594 | BR:Coi152.2:14 | BR:Coi152.1:14 | BR:Coi51:13 | RGBMCS**3** | $1.53 \times 10^{-32}$ |
| | 4 | 1 | 2399 | BR:Vic08:10, BR:Coi37.2:13 | BR:Coi51:13 | BR:Vic04.2:10 | RGBMC**S**3 | $1.19 \times 10^{-9}$ |
| EuYMV | 1 | 2683 | 9 | BR:Cha925:11, BR:Cha920:11, BR:Cha920.1:11, BR:Cha925.1:11 | BR:Cha510:10 | BR:Ats504:10 | RGBMCS**3** | $6.3 \times 10^{-33}$ |
| | 2 | 2678 | 7 | BR:Cha510.2:10 | Unknown | BR:Cha510.1:10 | GBMCS**3** | $5.69 \times 10^{-27}$ |
| | 3 | 2618 | 5018 | BR:Ats3.1:09, BR:Ats3.2:09, BR:Ats3.4:09 | BR:Plm933:11 | BR:Ats3.3:09 | RGBMCS**3** | $2.05 \times 10^{-19}$ |

| | | | | | | | | |
|---|---|---|---|---|---|---|---|---|
| | 4 | 2702 | 106 | BR:Cha1179.3:14, BR:Cha1179:14 | BR:Str536:10 | BR:Cha1179.2:14 | MCS**3** | 1.05 x $10^{-16}$ |
| | 5 | 25 | 2620 | BR:Sta946:11, BR:Ats3.3:09, BR:Ats504:10, BR:Ats504.1:10, BR:Trm939:11 | Unknown | BR:Sau937:11 | RGBM**C**S3 | 2.36 x $10^{-9}$ |
| | 6 | 42 | 2629 | BR:Cha510:10, BR:Non1003:11 | BR:Sau937:11 | BR:Cha510.1:10 | R**M**C3 | 1.36 x $10^{-9}$ |
| | 7 | 5168 | 2689 | BR:Smm61:09, BR:Ara1133:11, BR:Smm61.1:09 | BR:Str536:10 | Unknown | RGBMC**S** | 1.45 x $10^{-14}$ |
| | 8 | 2851 | 4980 | BR:MS:Mir2:07 | BR:MS:Mir1:07 | Unknown | RGBMC**S** | 3.05 x $10^{-17}$ |
| | 9 | 2526 | 44 | BR:Sag59:12 | BR:Sag59.1:12 | BR:Sag43:12 | RG**B**M3 | 6.02 x $10^{-8}$ |
| | 10 | 40 | 2676 | BR:Sag61.1:12 | BR:Sag61.2:12 | Unknown | RGBMC**S**3 | 1.06 x $10^{-8}$ |
| | 11 | 5176 | 2692 | BR:Str942:11, BR:Plm933:11, BR:Pon1000:11, BR:Cha1179.1:14, BR:Cha1179.2:14 | BR:Cha14.3a:09 | BR:Sta51:09 | RM**C**3 | 1.20 x $10^{-4}$ |
| | 12 | 35 | 2750 | BR:Sag57:12 | BR:Sag56:12 | BR:Sag61.2:12 | GBMC**S**3 | 1.01 x $10^{-6}$ |
| MaYSV | 1 | 2629 | 5085 | BR:Sti29:11, BR:Crb1:11, BR:Crb2:11, BR:Sti4:11, BR:PE:CAU:22:1, BR:PE:CAU:22:2, BR:PE:CAU:22:3, BR:PE:CAU:23:1, BR:PE:CAU:23:2 | BR:Oaf25.1:11 | BR:Sti34:11 | RGBMCS**3** | 3.78 x $10^{-9}$ |
| | 2 | 2865 | 5100 | BR:Oaf27:11, BR:Oaf24:11, BR:Oaf25.1:11, BR:Oaf25.2:11, BR:Oaf28:11, BR:Sti9.1:11, BR:Sti9.2:11, BR:Sti9.3:11, BR:Sti10:11, BR:Sti35:11 | Unknown | BR:Sti26:11 | RGBMCS**3** | 1.04 x $10^{-18}$ |
| ToSRV | 1 | 2516 | 41 | BR:Flo202.2:08 | BR:Flo203:08 | BR:Flo202.1:08 | BMCS**3** | 1.06 x $10^{-11}$ |
| | 2 | 2620 | 5134 | BR:Coi179:14 | Unknown | BR:Coi93:13 | MC**S**3 | 7.20 x $10^{-9}$ |

[*] Breakpoints correspond to the artificial joint between the two concatenated genomic components.

[#] R, Rdp; G, Geneconv; B, Boostcan; M, Maxichi; C, Chimaera; S, Siscan; 3, 3Seq.

[&] The reported *P*-value is from the method in bold and underlined, and is the lowest *P*-value calculated for the featured event.

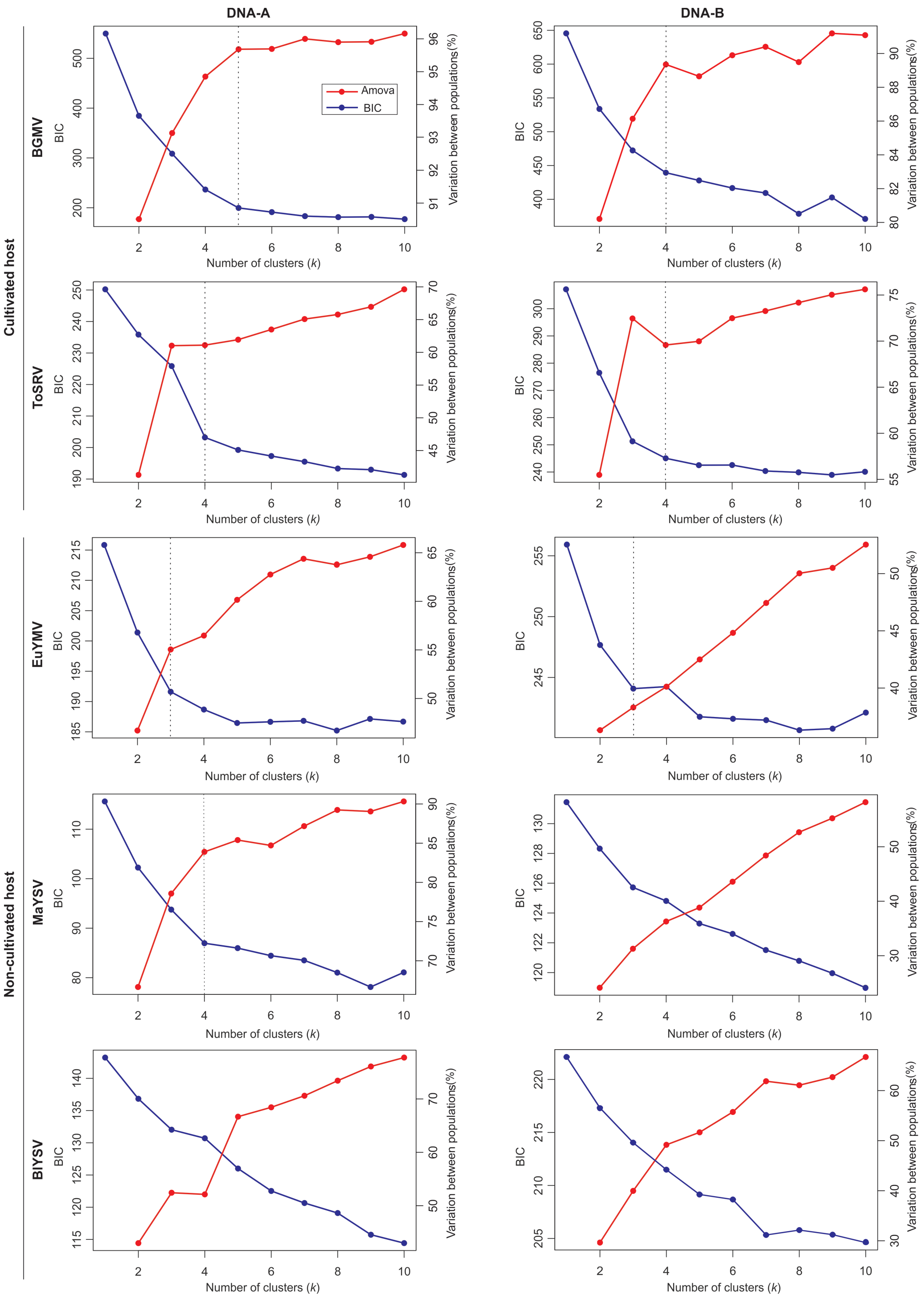


**Supplementary Figure S1.** Inference of the number of genetic clusters (*k*) using the *k*-means algorithm and analysis of molecular variance (AMOVA) for *Bean golden mosaic virus* (BGMV), *Blainvillea yellow spot virus* (BlYSV), *Euphorbia yellow mosaic virus* (EuYMV), *Macroptilium yellow spot virus* (MaYSV) and *Tomato severe rugose virus* (ToSRV) data sets. The *k*-means algorithm was run sequentially with values of *k* varying from 2 to 10. The choice of the number of clusters (indicated by the vertical dotted line) was primarily based on the minimum number of *k* after which the Bayesian Information Criterion (BIC, blue lines) decreases by an insignificant amount, as proposed by Jombart *et al.* (98). In addition, the percentage of variation among genetic clusters was plotted according to AMOVA analysis (red lines). A number of plots do not have the vertical dotted line due to unclear genetic cluster identification.

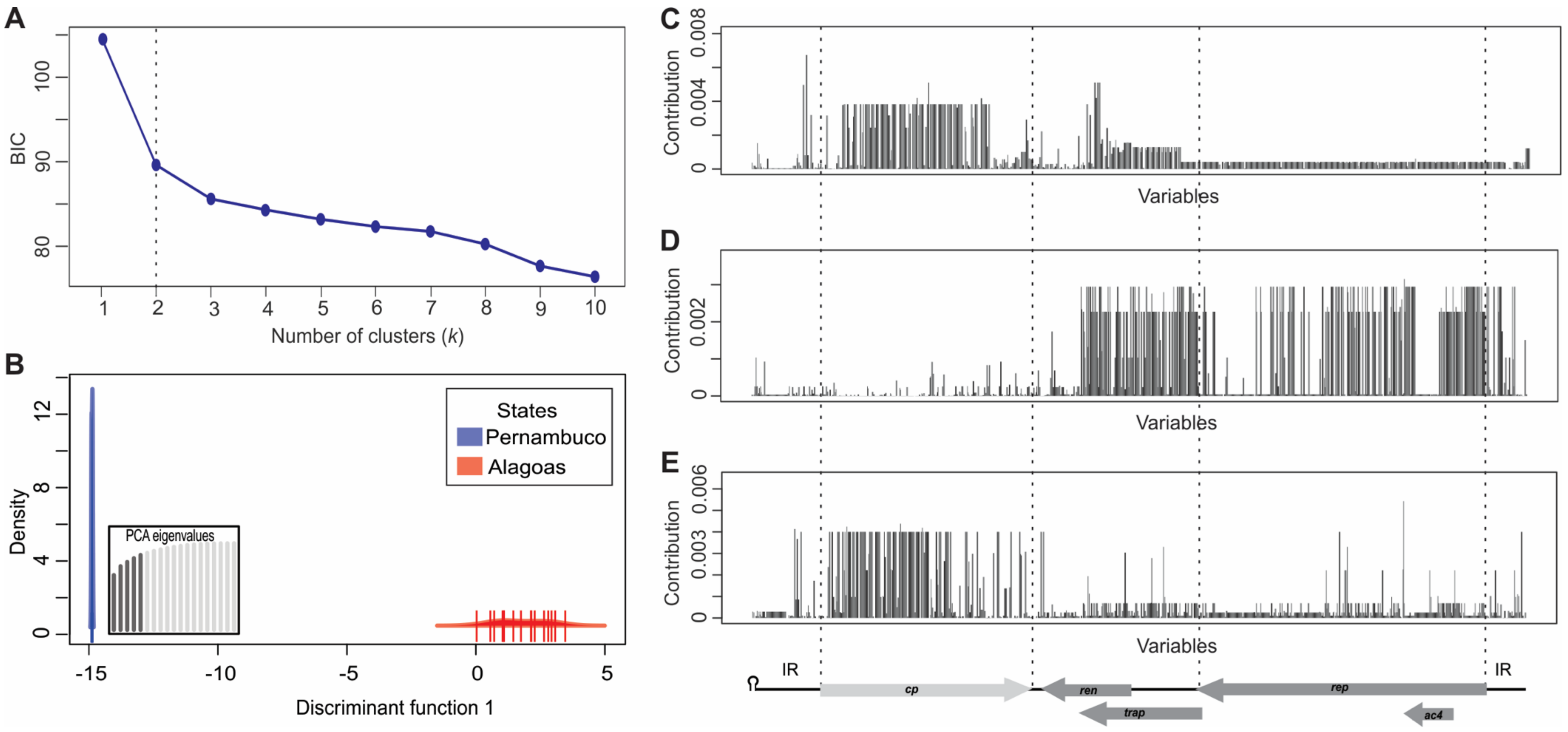


**Supplementary Figure S2.** Analysis of population subdivision using Discriminant Analysis of Principal Components (DAPC) for *Macroptilium yellow spot virus* (MaYSV) DNA-A using a data set without recombinants blocks. (**A**) Number of genetic clusters (*k*) inferred using the *k*-means algorithm. The algorithm was run sequentially with values of *k* varying from 2 to 10. The vertical dotted line indicates the number of genetic clusters chosen for DAPC analysis. (**B**) Density of isolates on discriminant function 1, colored according to the geographical origin of each genetic cluster. **(C-E)** Sites that contribute most to population subdivision when recombinants blocks are excluded (**C**) and when they are considered (**D** and **E**). The heights of the bars correspond to the contribution of the corresponding site.

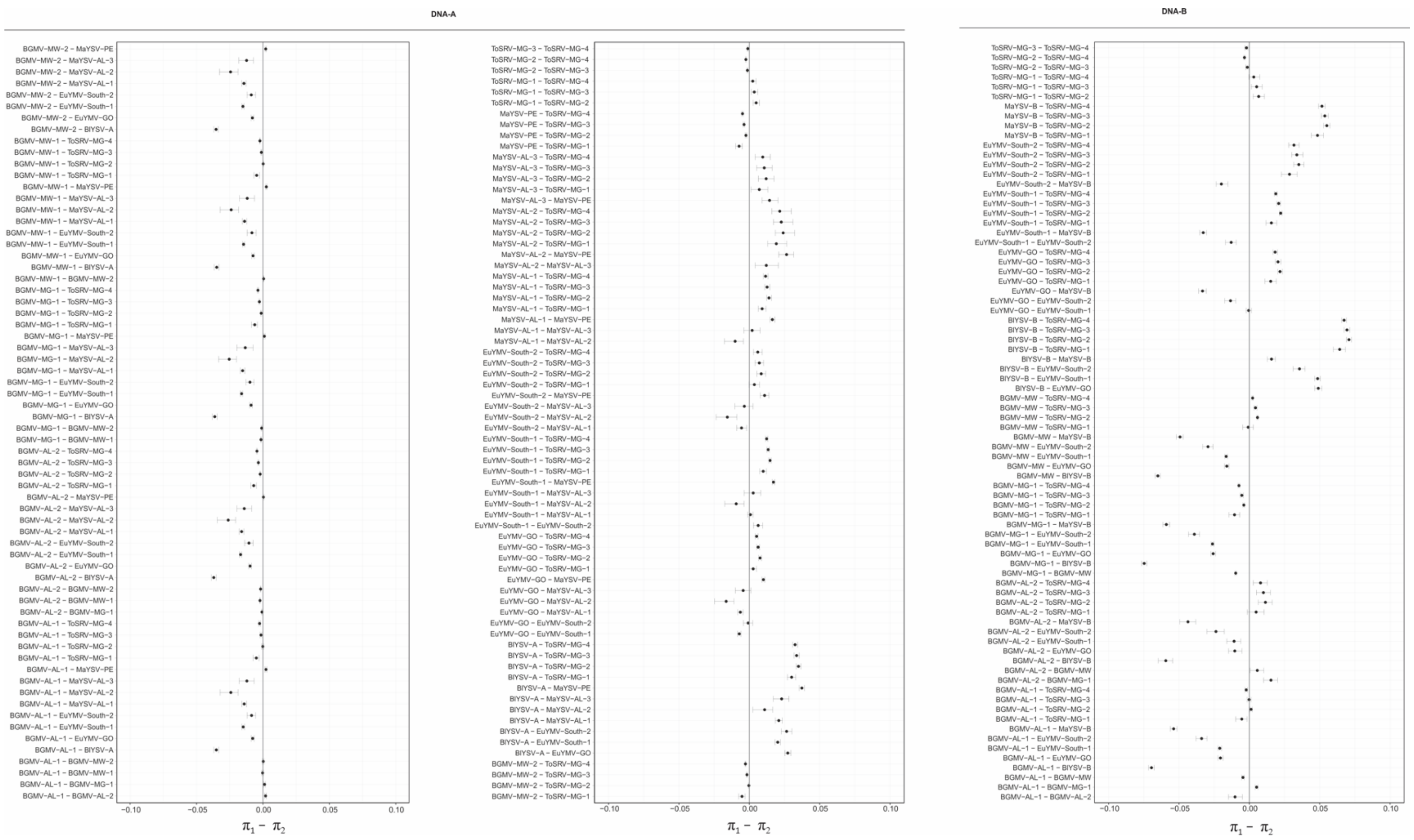


**Supplementary Figure S3.** Statistical significance of the difference between the average pairwise number of nucleotide differences per site (nucleotide diversity, π) calculated for genetic clusters inferred by Discriminant Analysis of Principal Components (DAPC) for the DNA-A and DNA-B of *Bean golden mosaic virus* (BGMV), *Blainvillea yellow spot virus* (BlYSV), *Euphorbia yellow mosaic virus* (EuYMV), *Macroptilium yellow spot virus* (MaYSV) and *Tomato severe rugose virus* (ToSRV). Ninety-five percent bootstrap confidence intervals (CIs) for the difference between π values were estimated from 1,000 nonparametric simulations. Confidence intervals that include the value "zero" denote no statistically significant difference between the means.

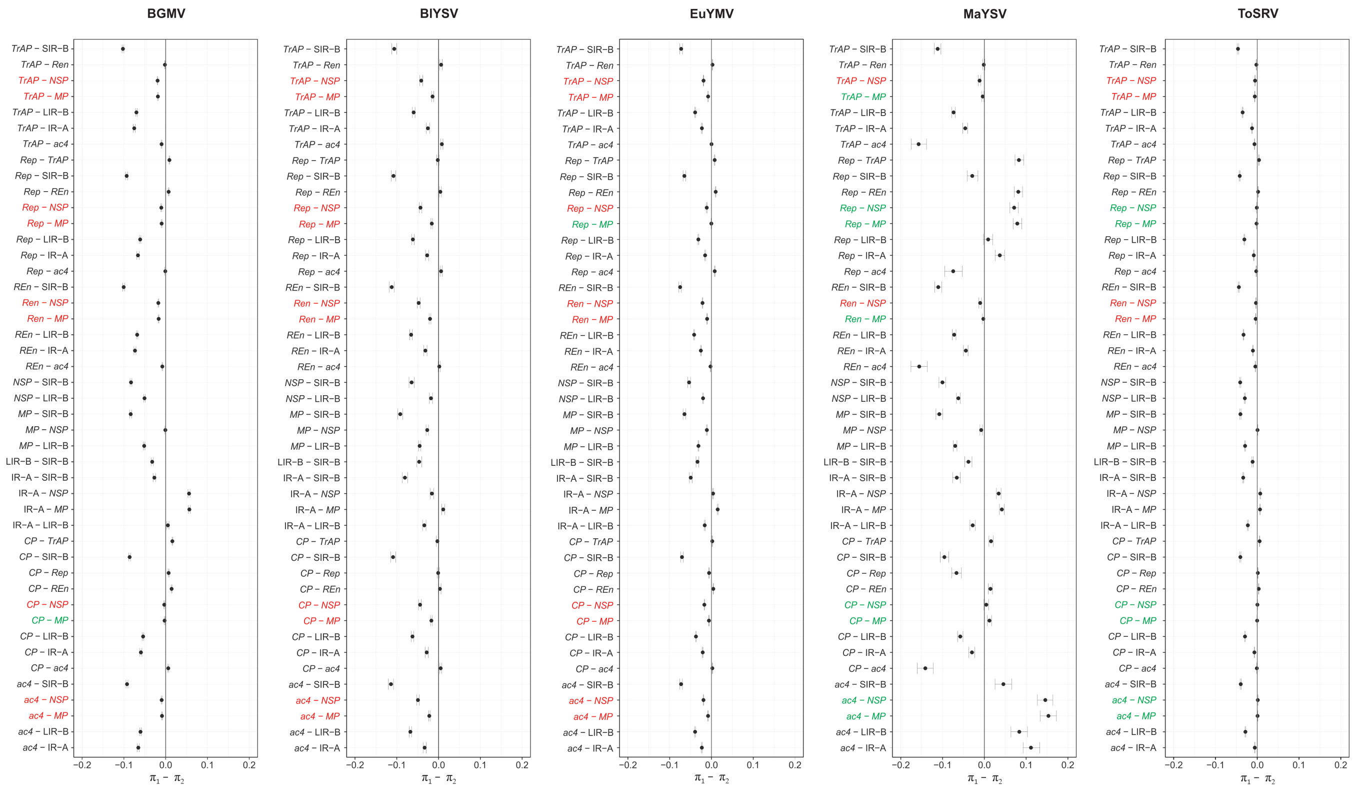


**Supplementary Figure S4.** Statistical significance of the difference between the average pairwise number of nucleotide differences per site (nucleotide diversity, π) calculated for nucleotide sequences of the *CP* (capsid protein), *Rep* (replication-associated protein), *REn* (replication enhancer protein), *TrAP* (trans-activating protein), *AC4*, *MP* (movement protein) and *NSP* (nuclear shuttle protein) genes, the DNA-A intergenic region (IR-A), DNA-B large intergenic region (LIR-B) and DNA-B small intergenic region (SIR-B) for *Bean golden mosaic virus* (BGMV), *Blainvillea yellow spot virus* (BlYSV), *Euphorbia yellow mosaic virus* (EuYMV), *Macroptilium yellow spot virus* (MaYSV) and *Tomato severe rugose virus* (ToSRV) data sets. Ninety-five percent bootstrap confidence intervals (CIs) for the difference between π values were estimated from 1,000 nonparametric simulations. Confidence intervals which include the value "zero" denote no statistically significant difference between the means. Comparisons where DNA-A genes are more variable than DNA-B genes are highlighted in red. Comparisons where there is no difference or DNA-B genes are more variable than DNA-A genes are highlighted in green.

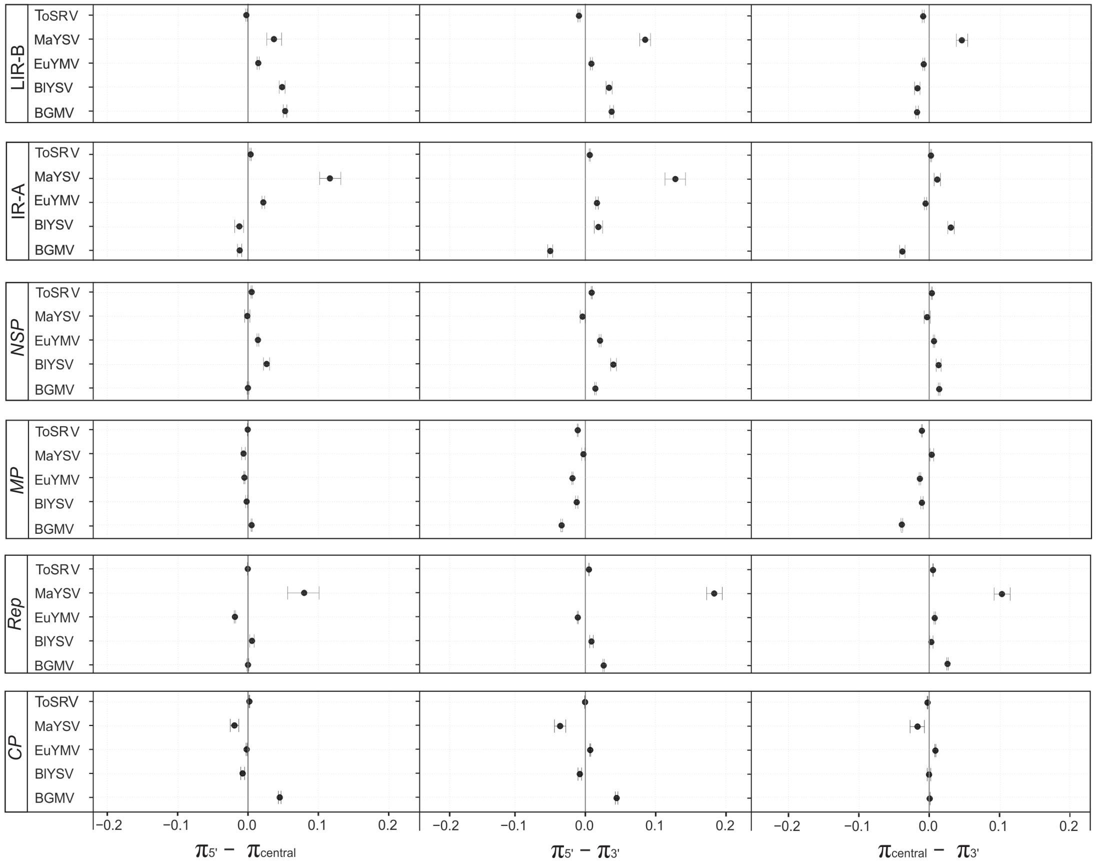


**Supplementary Figure S5.** Statistical significance of the difference between the average pairwise number of nucleotide differences per site (nucleotide diversity, π) calculated for the 5'-terminal, central and 3'-terminal regions of the *CP* (capsid protein), *Rep* (replication-associated protein), *MP* (movement protein) and *NSP* (nuclear shuttle protein) genes, the DNA-A intergenic region (IR-A) and the DNA-B large intergenic region (LIR-B) for *Bean golden mosaic virus* (BGMV), *Blainvillea yellow spot virus* (BlYSV), *Euphorbia yellow mosaic virus* (EuYMV), *Macroptilium yellow spot virus* (MaYSV) and *Tomato severe rugose virus* (ToSRV) populations. Ninety-five percent bootstrap confidence intervals (CIs) for the difference between π values were estimated from 1,000 nonparametric simulations. Confidence intervals which include the value "zero" denote no statistically significant difference between the means.
